# Geometric constraints on human brain function

**DOI:** 10.1101/2022.10.04.510897

**Authors:** James C. Pang, Kevin M. Aquino, Marianne Oldehinkel, Peter A. Robinson, Ben D. Fulcher, Michael Breakspear, Alex Fornito

**Author notes:** These authors contributed equally to this work (James C. Pang, Kevin M. Aquino).

## Abstract

The brain’s anatomy constrains its function, but precisely how remains unclear. Here, we show that human cortical and subcortical activity, measured with magnetic resonance imaging under spontaneous and diverse task-evoked conditions, can be parsimoniously understood as resulting from excitations of fundamental, resonant modes of the brain’s geometry (i.e., its shape) rather than modes from complex inter-regional connectivity, as classically assumed. We then use these modes to show that task-evoked activations across >10,000 brain maps are not confined to focal areas, as widely believed, but instead excite brain-wide modes with wavelengths spanning >60 mm. Finally, we confirm theoretical predictions that the close link between geometry and function is explained by a dominant role for wave-like dynamics, showing that such dynamics can reproduce numerous canonical spatiotemporal properties of spontaneous and evoked recordings. Our findings challenge prevailing views of brain function and identify a previously under-appreciated role of brain geometry that is predicted by a unifying and physically principled approach.

## MAIN TEXT

The dynamics of many natural systems are fundamentally constrained by their underlying structure. For instance, the shape of a drum influences its acoustic properties, the morphology of a riverbed shapes underwater currents, and the geometry of a protein constrains the molecules with which it interacts ^1,2^. The nervous system is no exception, with the rich and complex spatiotemporal dynamics of anatomically distributed neuronal populations being supported by their intricate web of axonal interconnectivity ^3,4^. Several studies have revealed correlations between various properties of brain connectivity and activity ^5^, but precisely how spatiotemporal patterns of neural dynamics are constrained by a relatively stable neuroanatomical scaffold remains unclear.

In diverse areas of physics and engineering, structural constraints on system dynamics can be understood via the system’s eigenmodes, which are fundamental spatial patterns corresponding to the natural, resonant modes of the system ^6,7^. In the linear regime, such as brain activity under normal (i.e., non-seizure-like) conditions ^8^, eigenmodes (hereafter also referred to as modes) offer a particularly powerful and rigorous formalism for linking brain anatomy with the physical processes that shape its activity. Through this lens, spatiotemporally patterned neuronal dynamics are viewed as emerging from excitations of the brain’s structural eigenmodes, much like the harmonics of a plucked violin string arise from vibrations of its resonant modes.

Just as the resonant frequencies of a violin string are determined by its length, density, and tension, the eigenmodes of the brain are determined by its structural––physical, geometric, and anatomical– –properties. Do any of these specific structural properties make a dominant contribution to dynamics? Here, we test between two influential and competing theories that make different predictions about which key elements of brain structure shape dynamics and function.

One classical perspective, which represents the dominant paradigm in neuroscience and has its roots in Ramon y Cajal’s neuron doctrine ^9^, Brodmann’s cytoarchitectonics ^10^, and over a century of work localizing functions to specific brain regions ^11,12^, is that spatiotemporal patterns of neural dynamics arise from interactions between discrete, functionally specialized cell populations connected by a topologically complex array of short- and long-range axonal connections ^13,14^. In humans, these connections can be estimated at macroscopic scales with diffusion MRI (dMRI) to yield a graph-based structural connectivity matrix or *connectome* ^15^. This approach has been used extensively to understand brain organization and dynamics ^13,15,16^, and recent work has proposed that eigenmodes derived from such discrete connectome models, referred to here as *connectome eigenmodes*, can be used to reconstruct the spatial patterns of canonical functional networks of the human cortex mapped with functional MRI (fMRI) ^17–19^.

A limitation of this discrete connectomic-based view is that it relies on an abstract representation of brain anatomy that does not directly account for its physical properties and spatial embedding (i.e., geometry and topology). These characteristics are explicitly incorporated into a broad class of *neural field theories* (NFTs) ^20–25^ that describe mean-field neural dynamics on spatial scales >0.5 mm (Supplementary Material-S1). In particular, a common, physiologically-constrained form of NFT has unified a diverse range of empirical phenomena ^25,26^ by treating cortical activity as a superposition of traveling waves propagating through a physically continuous sheet of neural tissue. In this theory, neural interactions between different cortical locations are approximated by a homogeneous spatial kernel that declines roughly exponentially with distance ^27^. This approximation is supported by experimental evidence showing that the organization of the nervous systems of different species is universally governed by an exponential distance rule (EDR) ^4,25,28,29^.

NFT predicts that the intrinsic geometry of the brain physically shapes and imposes boundary conditions on any emerging spontaneous and evoked dynamics ^30–32^. A remarkable corollary of this view is that, if we prioritize spatial and physical constraints on brain anatomy, we only need to consider the shape of the brain, and not its full array of topologically complex axonal interconnectivity, to understand dynamics. More formally, the theory predicts that eigenmodes derived from brain geometry––hereafter referred to as *geometric eigenmodes*––represent a more fundamental structural constraint on dynamics than the connectome ^30–32^. This view stands in stark contrast to the classical view that complex patterns of inter-regional anatomical connectivity shape brain activity ^33^.

Here, we test these competing views of the brain with the aim of identifying fundamental structural constraints on human brain dynamics. In line with theoretical predictions from NFT, we show that diverse experimental fMRI data from spontaneous and task-evoked recordings in the human neocortex can be more parsimoniously explained by eigenmodes derived from cortical geometry (geometric eigenmodes) rather than those obtained from a combination of topologically complex short- and long-range connectivity (connectome eigenmodes). We further confirm that stimulus-evoked activity is dominated by excitations of geometric eigenmodes with long spatial wavelengths, challenging classical views that such activity is localized to focal, spatially isolated clusters. To directly link these structural constraints to the physical processes driving brain dynamics, we use a generative model to show how wave dynamics unfolding on the geometry of the cortex can explain diverse features of functional brain organization. Finally, we show that the close relationship between geometry and function revealed by eigenmodes extends to non-neocortical structures, indicating that this link is a universal property of brain organization.

## RESULTS

### Eigenmodes of cortical geometry parsimoniously explain neocortical activity

We first examine the degree to which geometric eigenmodes can explain diverse aspects of human neocortical activity. To derive the eigenmodes, we first approximate cortical geometry using a triangular mesh representation, comprising 32,492 vertices in each hemisphere, taken from a population-averaged template of the neocortical surface ^34^ (Fig. 1A). We then construct the Laplace-Beltrami operator (LBO) from this surface mesh, which captures spatial variations of the cortical manifold by accounting for local vertex-to-vertex relations and curvature ^35^ (Supplementary Material-S2), and solve the eigenvalue problem,

**Fig. 1.**
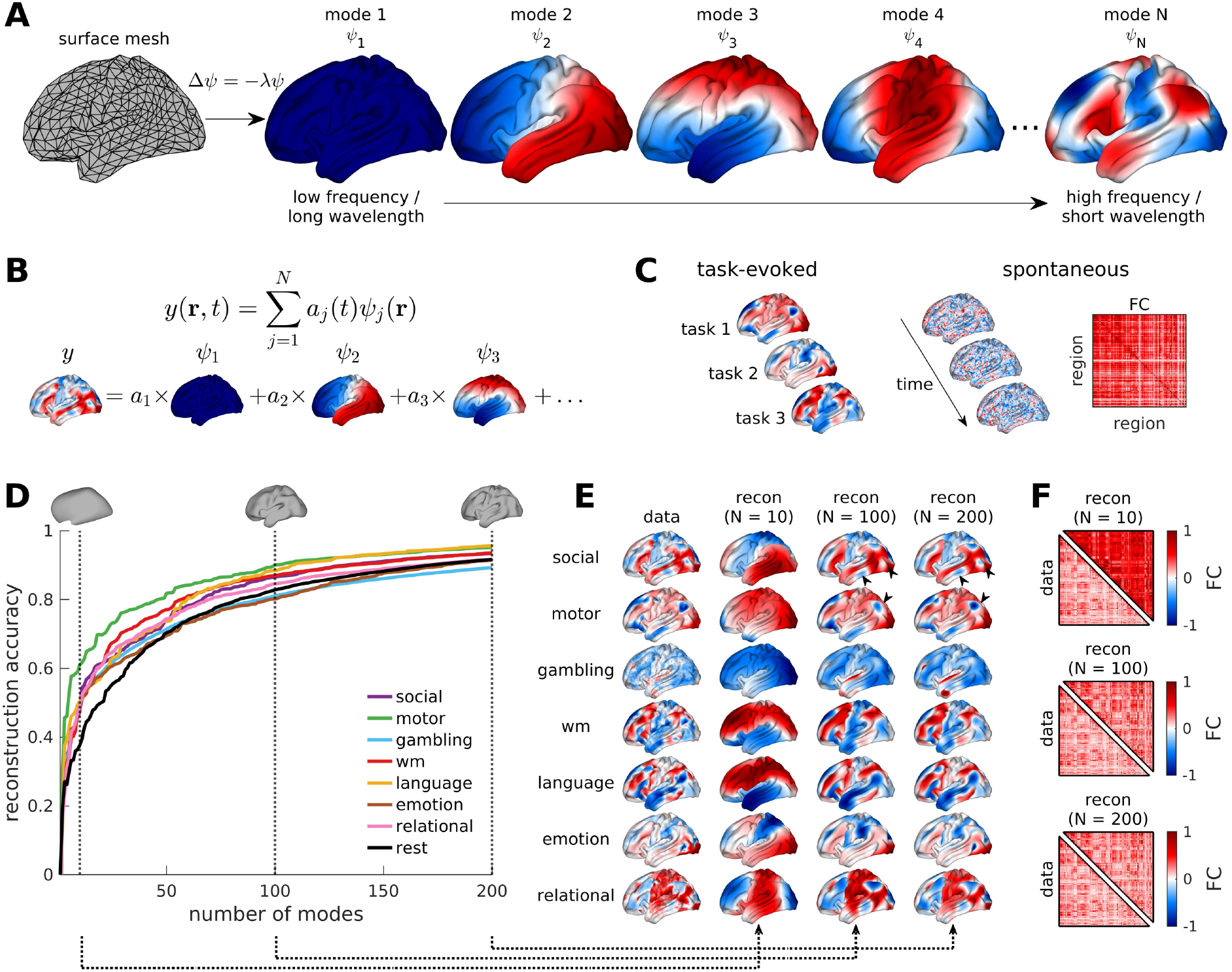
Eigenmodes of cortical geometry as a compact representation of macroscale neocortical activity. (**A**) Geometric eigenmodes are derived from a mesh representation of the cortical by solving the eigenvalue problem Δ*ψ* = −*λψ* (Eq. (1)). The modes *ψ*_1_, *ψ*_2_, *ψ*_3_, …, *ψ*_*N*_ are ordered from low to high spatial frequencies (long to short spatial wavelengths). Negative–zero–positive values are colored as blue–white–red. (**B**) Mode decomposition of brain activity data. The example shows how a spatial map, *y*(***r***), can be decomposed as a sum of modes weighted by *a*_*j*_. (**C**) We reconstruct task-evoked data using spatial maps of activation for a diverse range of stimulus contrasts (left). We also reconstruct spontaneous activity by decomposing the spatial map at each time frame and generating a region-to-region functional coupling (FC) matrix (right). (**D**) Reconstruction accuracy of 7 key HCP task-contrast maps (Supplementary Material-S4.2 and Table S2) and resting-state FC as a function of the number of modes. The insets show cortical-surface reconstructions demonstrating the spatial scales relevant to the first 10, 100, and 200 modes corresponding to spatial wavelengths of ∼120 mm, ∼40 mm, and ∼30 mm, respectively. (**E**) Group-averaged empirical task-activation maps and reconstructions obtained using 10, 100, and 200 modes of the 7 key HCP task contrasts. wm = working memory. The black arrows show localized activation patterns that are more accurately reconstructed when using short-wavelength modes. (**F**) Group-averaged empirical resting-state FC matrices and reconstructions using 10, 100, and 200 modes.

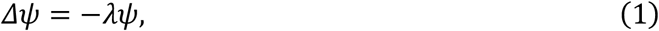

where *Δ* is the LBO and *ψ* = {*ψ*_1_(*r*), *ψ*_2_(***r***), …} is the family of geometric eigenmodes with corresponding family of eigenvalues, *λ* = {*λ*_1_, *λ*_2_, …}. The eigenvalues are ordered sequentially according to the spatial frequency or wavelength of the spatial patterns of each mode (Fig. 1A and Fig. S1); i.e., 0 ≤ *λ*_1_ ≤ *λ*_2_ ≤ ⋯, where *λ*_1_ corresponds to the mode with the longest spatial wavelength (Supplementary Material-S2).

Eigenmodes are orthogonal, forming a complete basis set to decompose spatiotemporal dynamics unfolding on the cortex into different spatial frequencies. We thus use the geometric eigenmodes to decompose empirical data, *y*(***r,*** *t*), measured at spatial location ***r*** and time *t*, as a weighted sum of modes (Fig. 1B),

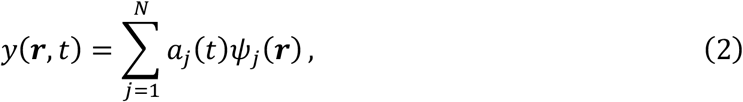

where *a*_*j*_ is the amplitude of mode *j* obtained via integration of the dot product of *y*(***r,*** *t*) and *ψ*_*j*_(***r***) over the cortical surface or via a statistical general linear model (Supplementary Material-S3) and *N* is the number of modes retained (here we use *N* = 200 modes).

Using this decomposition, we evaluate the accuracy of geometric eigenmodes in capturing both task-evoked and spontaneous brain activity (Fig. 1C) measured in 255 healthy individuals from the Human Connectome Project ^36^ (HCP; Supplementary Material-S4). For task-evoked activity, we map 47 task-based contrasts drawn from 7 different tasks, representing distinct evoked activation patterns. We then reconstruct each individual’s activation map using an increasing number of modes up to a maximum of 200 (Fig. 1D). For spontaneous, task-free (so-called “resting-state”) activity, we reconstruct the spatial map of activity at each time frame and then generate a region-to-region functional coupling (FC) matrix, describing correlations of activity between 180 discrete brain regions per hemisphere defined by a widely used parcellation based on multimodal neuroimaging data ^37^. To allow direct comparison between task-evoked and spontaneous recordings, we apply the same regional parcellation to the task-evoked data, reducing 32,492 data points at vertex resolution (millimeter scale) to 180 data points at parcel resolution (centimeter scale) (Supplementary Material-S5). Finally, we quantify reconstruction accuracy by calculating the correlation between the empirical and reconstructed task-evoked activation maps and spontaneous FC matrices (Figs. 1D–E).

We observe that reconstruction accuracy increases with an increasing number of modes across all task contrasts and in the resting-state, with *r* ≥ 0.38 already achieved using just *N* = 10 modes (Fig. 1D). Large-scale modes are also recruited distinctly across different tasks, suggesting that particular stimuli excite specific modes (Fig. 1E). Improvements in reconstruction accuracy slow down after 10 modes, reaching *r* ≥ 0.80 at approximately *N* = 100 modes with only incremental increases in reconstruction accuracy beyond this point. Since the first 100 modes have wavelengths above ∼40 mm, and the inclusion of shorter-wavelength modes serves mainly to refine reconstruction of localized patterns (arrows in Fig. 1E), our findings suggest that the data are predominantly comprised of spatial patterns with long spatial wavelengths (see next section for a more detailed analysis). These results are consistent across all 47 HCP task contrasts (Fig. S2) and parcellations of varying resolutions (Fig. S3), but data parcellated at higher resolutions require more modes to achieve high reconstruction accuracy due to the low-pass spatial filtering effect of coarser parcellations. Our results are also not affected by our use of a population-averaged cortical surface template (rather than individual-specific surfaces) to derive the geometric eigenmodes (Supplementary Material-S2; Fig. S4). However, there are some participants where individual-specific eigenmodes perform slightly better, especially in reconstructing task-activation maps (Fig. S5), but their accuracies eventually converge with template-derived eigenmodes at short wavelengths (Fig. S6). These findings indicate that cortical geometric eigenmodes form a compact representation that captures diverse aspects of task-evoked and spontaneous cortical activity. Moreover, they show that such activity is dominated by long-wavelength, large-scale eigenmodes.

We next test the hypothesis that geometric eigenmodes provide a more parsimonious and fundamental description of dynamics than eigenmodes derived from a graph-based connectome approximation. To this end, we compare the reconstruction accuracy of geometric eigenmodes against three alternative connectome-derived eigenmode basis sets (see Fig. 2A for a schematic).

**Fig. 2.**
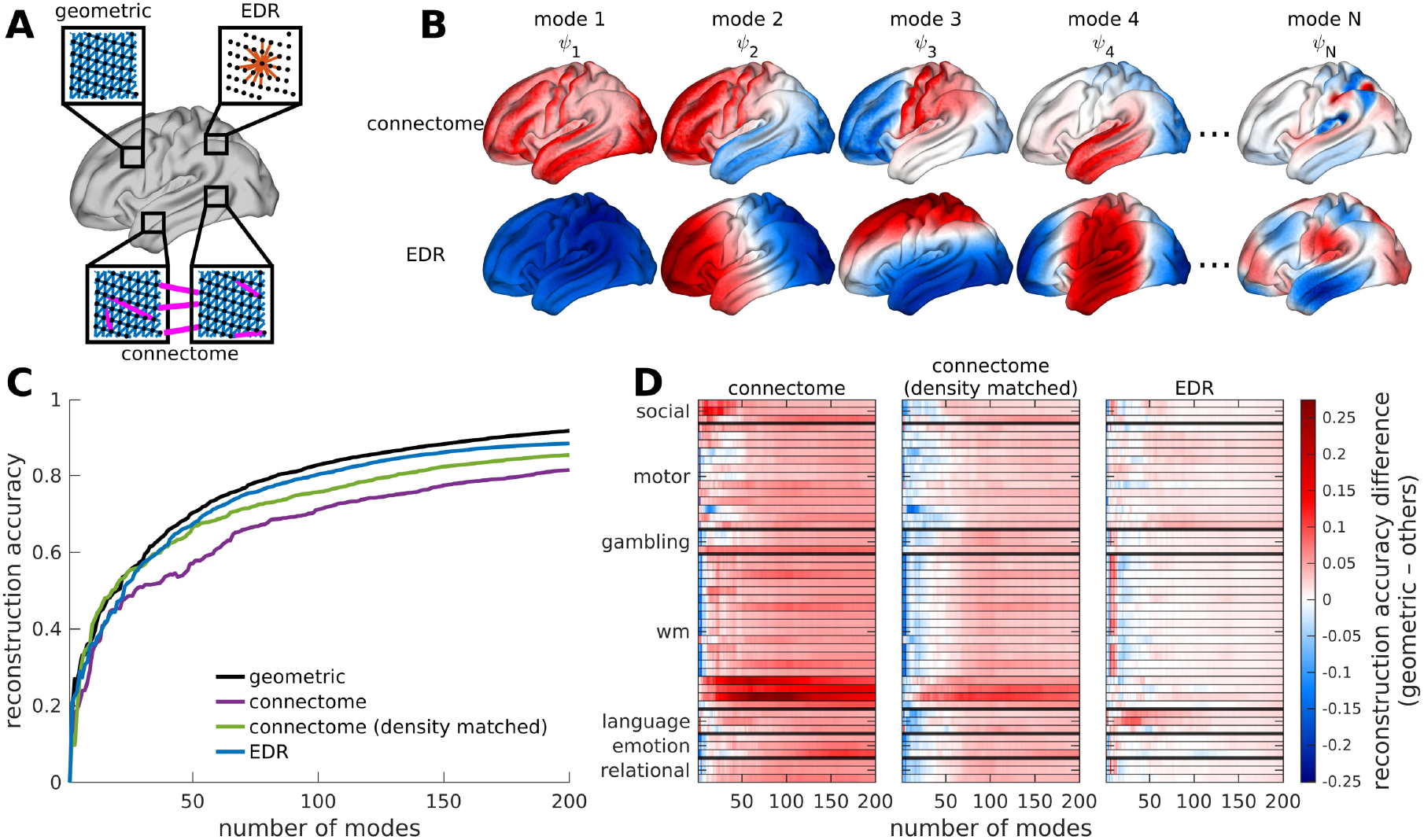
Geometric eigenmodes benchmarked against connectome-based eigenmodes. (**A**) Schematic of the anatomical properties used to derive the eigenmodes for cortical geometry, the connectome, and the exponential distance rule (EDR) connectome. Geometric eigenmodes rely on local surface mesh information such as links (blue) between neighboring surface mesh vertices (dots) and curvature. Connectome eigenmodes rely on local links between mesh vertices (blue) and short- and long-range connections (magenta) reconstructed empirically from dMRI. EDR eigenmodes rely on connections (red) generated from a stochastic wiring process where the probability of connection between vertices exponentially decays as a function of their distance (Supplementary Material-S6 and S7). (**B**) Example connectome and EDR eigenmodes. Negative–zero–positive values are colored as blue–white–red. Despite some similarities, the spatial patterns of the modes are distinct from those derived using cortical geometry (compare with Fig. 1A). (**C**) Reconstruction accuracy of resting-state FC matrices achieved by geometric, EDR, and two variants of connectome eigenmodes: one using a connectome as defined using prior methods ^38^ and the other with the same connection density as the EDR connectome to allow fair comparison (for other densities, see Fig. S9). (**D**) Difference in reconstruction accuracy of all 47 HCP task-contrast maps achieved by geometric eigenmodes and the other basis sets, as indicated by the text above each panel. Each row represents a different task contrast, which have been grouped here by broad types (Supplementary Material-S4.2 and Table S2). wm = working memory. Red indicates superior performance for geometric eigenmodes. Note that while there seems to be a performance advantage for connectome eigenmodes for reconstructions incorporating <10 modes relative to geometric eigenmodes, the reconstruction accuracy is generally low (average *r* = 0.42 across the different tasks) compared to the accuracy for 100 modes (average *r* = 0.71).

The first basis set is derived empirically from a connectome mapped with dMRI tractography at vertex resolution ^38^ (Supplementary Material-S6). The second basis set is derived from a connectome constructed synthetically according to a homogeneous stochastic wiring process governed by an exponential distance-dependent connection probability to mimic simple, EDR-like connectivity (Supplementary Material-S7). We threshold the empirical connectome to obtain a connection density of 0.10%, as done previously ^38^. The third basis set is derived from the empirical connectome thresholded at 1.55% to match the density of the EDR connectome (Supplementary Material-S7). The connectome, EDR, and density-matched connectome eigenmodes described above are derived from the graph Laplacian (a discrete counterpart of the LBO) of their respective connectivity matrices (Fig. 2B and Fig. S1; Supplementary Material-S6 and S7).

To summarize, geometric eigenmodes account for the intrinsic curvature of the cortical surface and local vertex-to-vertex relations in the surface mesh; connectome eigenmodes do not consider curvature but capture local spatial relations between points and short- and long-range connections measured with dMRI; and EDR eigenmodes account for the effect of a homogeneous, stochastic, distance-dependent connection rule without fully capturing the cortical geometry (Fig. 2A). Comparing these different basis sets thus allows us to disentangle the contributions to brain dynamics of cortical geometry from structural connectivity.

Direct comparison of the reconstruction accuracy of these different basis sets reveals that geometric eigenmodes consistently show the highest reconstruction accuracy across both spontaneous (Fig. 2C) and task-evoked (Fig. 2D) data. EDR eigenmodes perform nearly as well as the geometric eigenmodes, whereas connectome eigenmodes are the least accurate. This finding holds regardless of the parcellation used (Figs. S7 and S8), the specific connection density used to generate the connectome eigenmodes (Figs. S9 and S10), and whether we generate the connectome using a discrete regional parcellation rather than at vertex resolution (Fig. S11; Supplementary Material-S6). We additionally find that geometric eigenmodes show stronger out-of-sample generalization than principal components of the functional data itself (calculated via principal component analysis (PCA); Supplementary Material-S8 and Figs. S12 and S13) and better performance than Fourier spatial basis sets (Supplementary Material-S9 and Fig. S14). These results further demonstrate the robustness and generality of geometric eigenmodes as a basis set for brain function. Together, these findings support the prediction of NFT that brain activity is best represented in terms of eigenmodes derived directly from the shape of the cortex, and emphasize a fundamental role of geometry in constraining dynamics.

### Cortical activity is dominated by excitations of long-wavelength geometric modes

Reconstructions of both spontaneous and task-evoked data with geometric eigenmodes show that the spatial organization of brain activity is dominated by patterns with spatial wavelengths of ∼40 mm or longer (Figs. 1D–E). This result counters the assumptions of classical neuroimaging analyses, in which stimulus-evoked activations are mapped by thresholding statistical maps to identify focal, isolated clusters of heightened activity. This classical approach rests on the assumption that the focal clusters represent discrete brain regions putatively engaged by the stimulus and that subthreshold activity in other regions plays no role. The surprisingly long-wavelength content of task-activation data (Figs. 1D–E) suggests that classical procedures focus only on the tips of the iceberg and obscure the underlying spatially extended and structured patterns of activity evoked by the task (see Fig. S15 for an explanation as to why). These observations accord with the theoretical predictions of NFT and prior analyses of task-evoked electroencephalography (EEG) signals ^39–41^.

Here, we leverage the mode decomposition described in Fig. 1B to characterize the complete spatial pattern––the entire iceberg––of task-evoked activation (Supplementary Material-S10). To this end, we analyze the mean modal power spectrum obtained using a geometric mode decomposition of group-averaged unthresholded activation maps from the 47 task contrasts in HCP ^36,42^. As an independent replication, we also analyze 10,000 unthresholded activation maps from 1,178 independent experiments available in the NeuroVault repository ^43^ (Supplementary Material-S10), thus providing a comprehensive picture of the diversity of stimulus-evoked activation patterns mapped in the human brain.

Despite the diversity of stimuli, paradigms, and data-processing approaches used to acquire these activation maps, we observe that a large fraction of power in the maps is concentrated in the first 50 modes, corresponding to spatial wavelengths greater than ∼60 mm (Fig. 3A; similar results are found in each key HCP task-contrast map; Fig. S16). Using surrogate data, we confirm that these findings cannot be explained by the spatial smoothing induced by typical fMRI processing pipelines, which can filter out short-wavelength spatial patterns of activity (Fig. S17; Supplementary Material-S10). We further observe that incremental, sequential removal of long-wavelength modes has a much greater impact on reconstruction accuracy than removal of short-wavelength modes (Fig. 3B; Supplementary Material-S11). For instance, removing the top 25% long-wavelength modes (i.e., modes 1 to 50) yields a ∼40 to 60% drop in reconstruction accuracy, whereas removing the top 25% short-wavelength modes (i.e., modes 151 to 200) only yields a ∼2 to 4% drop in accuracy (Fig. 3B insets). These results indicate that, on temporal and spatial scales accessible with fMRI, evoked cortical activity comprises large-scale, nearly brain-wide spatial patterns, challenging classical views that such activity should be described in terms of discrete, isolated, and anatomically localized activation clusters.

**Fig. 3.**
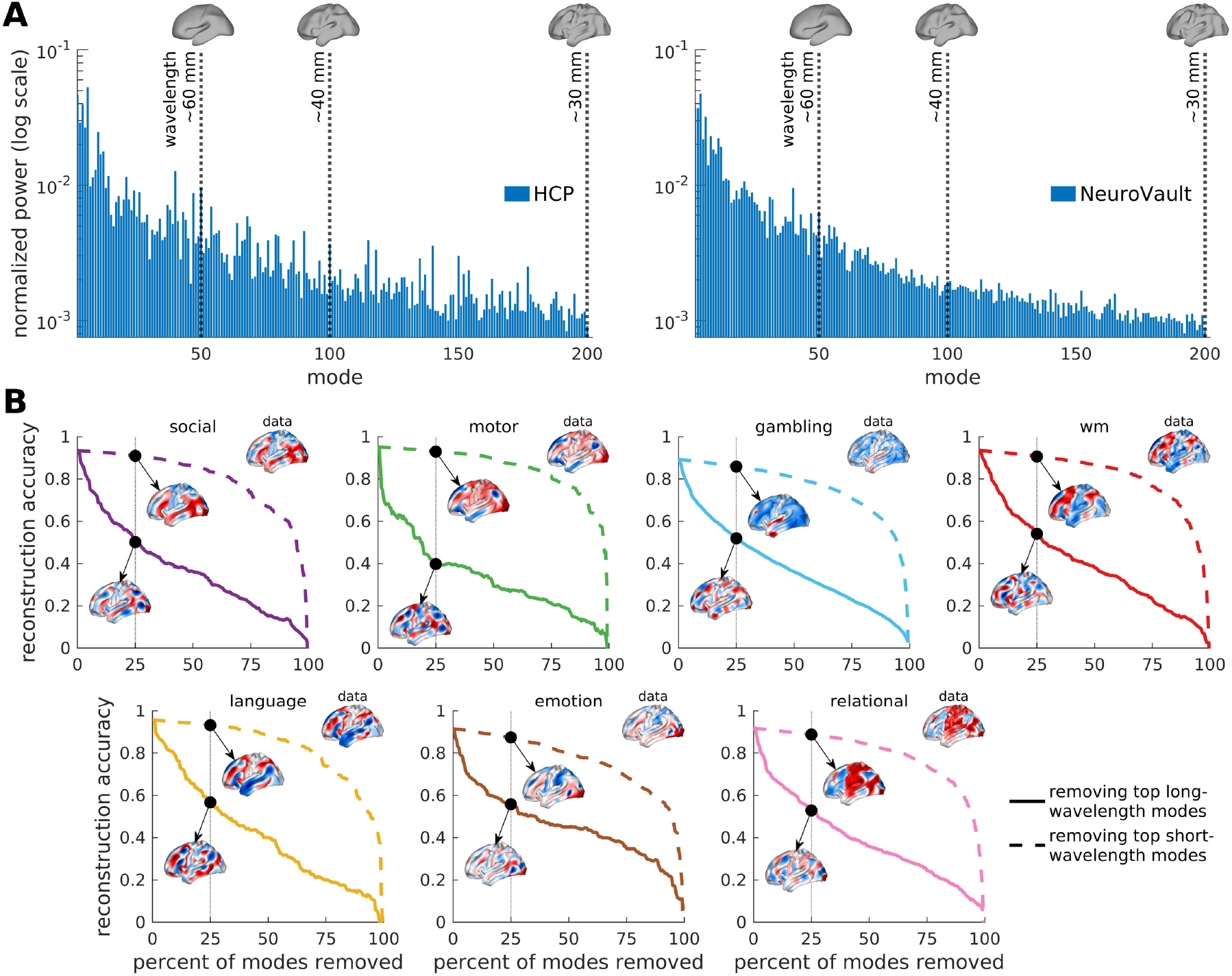
Task-evoked activity excites long-wavelength modes. (**A**) Normalized mean power spectra of 47 HCP task-contrast maps (left) and 10,000 contrast maps from the NeuroVault database (right). The insets show cortical-surface reconstructions demonstrating the spatial scales relevant to the first 50, 100, and 200 modes corresponding to spatial wavelengths of ∼60 mm, ∼40 mm, and ∼30 mm, respectively. Contrast-specific spectra for the 7 key HCP task contrasts are presented in Fig. S16. (**B**) Reconstruction accuracy of 7 key HCP task-contrast maps as a function of the percent of modes (out of 200 modes) removed in the reconstruction process. wm = working memory. The solid and dashed lines correspond to the removal of the top long-wavelength and short-wavelength modes, respectively. The insets show group-averaged empirical activation maps (‘data’) and their reconstructions after removing 25% of modes. Negative–zero–positive values are colored as blue–white– red.

### Traveling waves and geometry explain diverse neocortical dynamics

Geometric eigenmodes of the cortex are obtained by solving the eigenvalue problem of the LBO, which is also known as the Helmholtz equation (Eq. (1) and Supplementary Material-S2). In physically continuous systems, the solutions of the Helmholtz equation correspond to the spatial projections of the solutions of a more general wave equation, such that the resulting eigenmodes inherently represent the vibrational patterns, or standing waves, of the system’s dynamics ^44^. This equivalence implies that the superior efficacy of geometric eigenmodes in reconstructing diverse patterns of brain activity results from a fundamental role of wave dynamics in shaping these patterns, as predicted by NFT. This prediction has been confirmed through models of EEG recordings ^26,45^, but waves across the whole brain have only recently been observed in fMRI signals ^46,47^ and thus far lack a theoretical explanation. Here, we use NFT and geometric eigenmodes to show that wave dynamics can provide a unifying account of diverse empirical and physiological phenomena observed at scales accessible with fMRI.

We model the neural activity *ϕ* of a neocortical location ***r*** at time *t* using an isotropic damped NFT wave equation without regeneration ^25^,

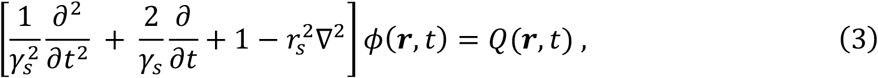

where γ_*S*_ is the damping rate, ***r***_*S*_ is the spatial length scale of local axonal projections, and *Q* is an external input (Supplementary Material-S12.1). Under this model, activity propagates between points on the neocortex through their white-matter connectivity, with a strength that decays approximately exponentially with distance (Supplementary Material-S1 and S12.1). To simulate resting-state neural activity, *Q* is a white noise input to mimic unstructured stochastic fluctuations ^26,48^. We compare the performance of this simple wave model to a biophysically-based neural mass model (balanced excitation-inhibition, or BEI, model) that has been used extensively to understand resting-state fMRI signals ^49,50^ (Fig. 4A). The neural mass model is closely aligned with the classical, connectome-centric view of brain function, representing dynamics as the result of interactions between neuronal populations in discrete anatomical regions, coupled according to an empirically measured connectome (Supplementary Material-S12.2).

**Fig. 4.**
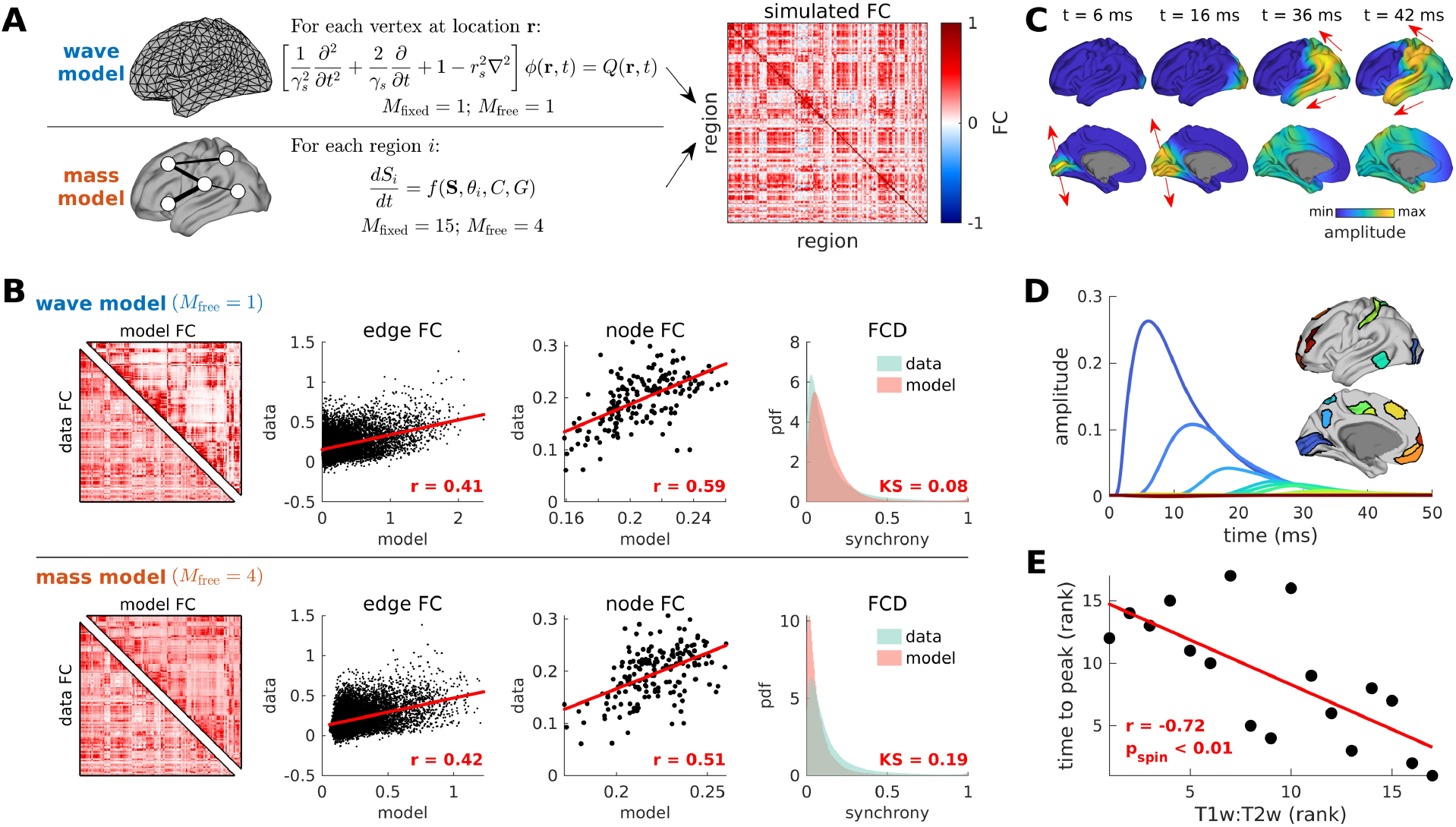
Traveling wave dynamics shape patterns of spontaneous and stimulus-evoked activity. (**A**) Simulation workflow using wave and neural mass models. For the wave model, the activity *ϕ*(***r*,** *t*) at location *r* and time *t* is governed by a wave equation with damping rate γ_*S*_, spatial length scale *r*_*S*_, and input *Q*(*r, t*). For the neural mass model, the activity *S*_*i*_(*t*) of region *i* is described by the function *f*, which depends on the activity of other regions ***S***, local population parameters *θ*_*j*_, and dMRI-derived structural connectivity between regions *C* and scaled by the global coupling parameter *G*. The model dynamics are used to calculate a simulated FC matrix (Supplementary Material-S12.4). *M*_fixed_ and *M*_free_ correspond to the number of fixed and free parameters of each model, respectively. (**B**) Comparison of data and model simulations based on various metrics, which from left to right are: FC matrix (for visual purposes), static pairwise FC (edge FC), static node-level average FC (node FC), and time-resolved FC dynamics (FCD). For edge FC and node FC, the red lines represent linear fits with Pearson correlation coefficient *r*. For FCD, the probability density function (pdf) of the similarity of global synchrony in data and model dynamics are compared using the Kolmogorov-Smirnov (KS) statistic. Note that the free parameters for each model are tuned to optimize data-model fitting using data that is independent of the data used to evaluate model performance (Supplementary Material-S12.5). (**C**) Wave propagation of activity after a 1 ms stimulation of the primary visual cortex (V1) from *t* = 1 to 2 ms. The arrows show the direction of propagation (Supplementary Video 1). (**D**) Activity profile of different regions in the visual cortical hierarchy. The insets show the spatial locations of the regions on the cortical surface colored following the activation profiles. (**E**) Relationship of the ranked activity profile time to peak and ranked T1w:T2w value of the regions in panel D. The red line represents a linear fit of the ranked variables with Spearman correlation coefficient *r* and spin-test *p*-value *p*_spin_ from 10,000 permutations (Supplementary Material-S12.7).

We first compare the efficacy of the two models in capturing distinct and commonly studied properties of spontaneous, task-free FC; namely, static pairwise FC (edge FC), node-level average FC (node FC), and time-resolved dynamic properties of FC (FCD) (Supplementary Material-S12.4). Across all FC-based benchmark measures, the wave model displays comparable or superior performance in reconstructing the empirical data relative to the neural mass model (Fig. 4B). Additionally, the wave model can better capture time-lagged properties ^46,47,51^ of empirical resting-state activity compared to the mass model (Supplementary Material-S12.6 and Fig. S19). This strong performance of the wave model is remarkable given its relative simplicity: the wave model only requires the geometry of the cortex (i.e., the surface mesh) as input and includes 1 fixed parameter and 1 free parameter (i.e., *r*_*S*_) for fitting to data (Supplementary Material-S12.1 and 11.5 and Fig. S20), whereas the neural mass model requires a dMRI-derived inter-regional anatomical connectivity and comprises 15 fixed parameters and 4 free parameters (Supplementary Material-S12.2 and S12.5). These considerations indicate that wave dynamics provide a more accurate and parsimonious mechanistic account of macroscale, spontaneous cortical dynamics captured by fMRI.

We next consider stimulus-evoked cortical activity in the wave model. We analyze cortical responses to sensory stimulation of primary visual cortex (V1), as it elicits a well-defined hierarchy of regional cortical responses ^52,53^ (Supplementary Material-S12.7). A 1 ms pulse input to V1 yields a propagating wave of activity that rapidly splits along the dorsal and ventral visual processing streams (Fig. 4C; see arrows and also Supplementary Video 1), consistent with the current mainstream understanding of hierarchical visual processing ^54^. Remarkably, this result indicates that geometric constraints on travelling waves of evoked activity are sufficient for the segregation of the dorsal and ventral processing streams, which have traditionally been thought to be mainly driven by complex patterns of layer-specific connectivity ^52,54,55^. Furthermore, the temporal profile of evoked responses across the visual system follows a well-defined temporal hierarchy, with higher-order association areas showing peak responses that are delayed and prolonged compared to visual lower areas (Fig. 4D). These findings thus indicate that this hierarchical ordering, which has previously been identified in experimental and modelling studies ^53,56,57^, emerges naturally from waves of excitation propagating through the cortical medium. Critically, this hierarchical temporal ordering of areal responses strongly correlates with an independent, anatomical measure of the cortical processing hierarchy based on non-invasive estimates of myeloarchitecture (T1w:T2w) ^58,59^ (Supplementary Material-S12.7). This correlation is particularly strong within the visual-processing hierarchy (*r* = −0.72, *p*_*Spin*_ < 0.01; Fig. 4E) but is also present when considering all cortical areas (*r* = −0.44, *p*_*Spin*_ = 0.037; Fig. S21). Together, our modelling results show how simple wave dynamics unfolding on the geometry of the cortex provide a unifying generative mechanism for capturing apparently complex properties of spatiotemporal brain activity.

### Geometry also constrains dynamics outside the neocortex

Our analyses have thus far focused on the strong coupling of geometry and dynamics in the neocortex. We next investigate this coupling in non-neocortical areas, focusing on the thalamus, striatum, and hippocampus because these structures have geometries that can be easily captured using MRI data and their functional organization has been extensively studied ^60^.

We first generalize our eigenmode analysis to 3D volumes using recently developed methods ^61^ (Supplementary Material-S13), yielding geometric eigenmodes that extend spatially through the 3D volume of each structure. Next, to fully capture the macroscale functional organization of these non-neocortical structures, we apply a widely used manifold-learning procedure to voxel-wise FC data to obtain the key *functional gradients* in each structure ^62^ (Supplementary Material-S14). These functional gradients describe the principal axes of spatial organization dictated by similarities in FC, thus representing the dominant modes of variation in functional organization, ordered according to the percentage of variance in FC similarity that they explain.

The spatial profiles of the first three functional gradients of the thalamus, striatum, and hippocampus (accounting for 24%, 50%, and 47% of the variance in FC similarity, respectively) show a near-perfect match to the first three geometric eigenmodes (Figs. 5A–C; spatial correlations *r* ≥ 0.93). This tight correspondence generalizes out to the first 20 gradients and first 20 modes of each structure (respectively accounting for 49%, 70%, and 68% of the total variance in FC similarity), with all absolute spatial correlations |*r*| ≥ 0.5, except for the 20th gradient and 20th mode in the striatum and hippocampus (Figs. 5D–F). This strong relationship is striking given that the functional gradients are generated via a complex processing pipeline applied to fMRI-derived FC measures, while the eigenmodes are derived simply from each structure’s geometry, independent of the functional data. These findings suggest that the functional organization of non-neocortical structures derives directly from their geometric eigenmodes.

**Fig. 5.**
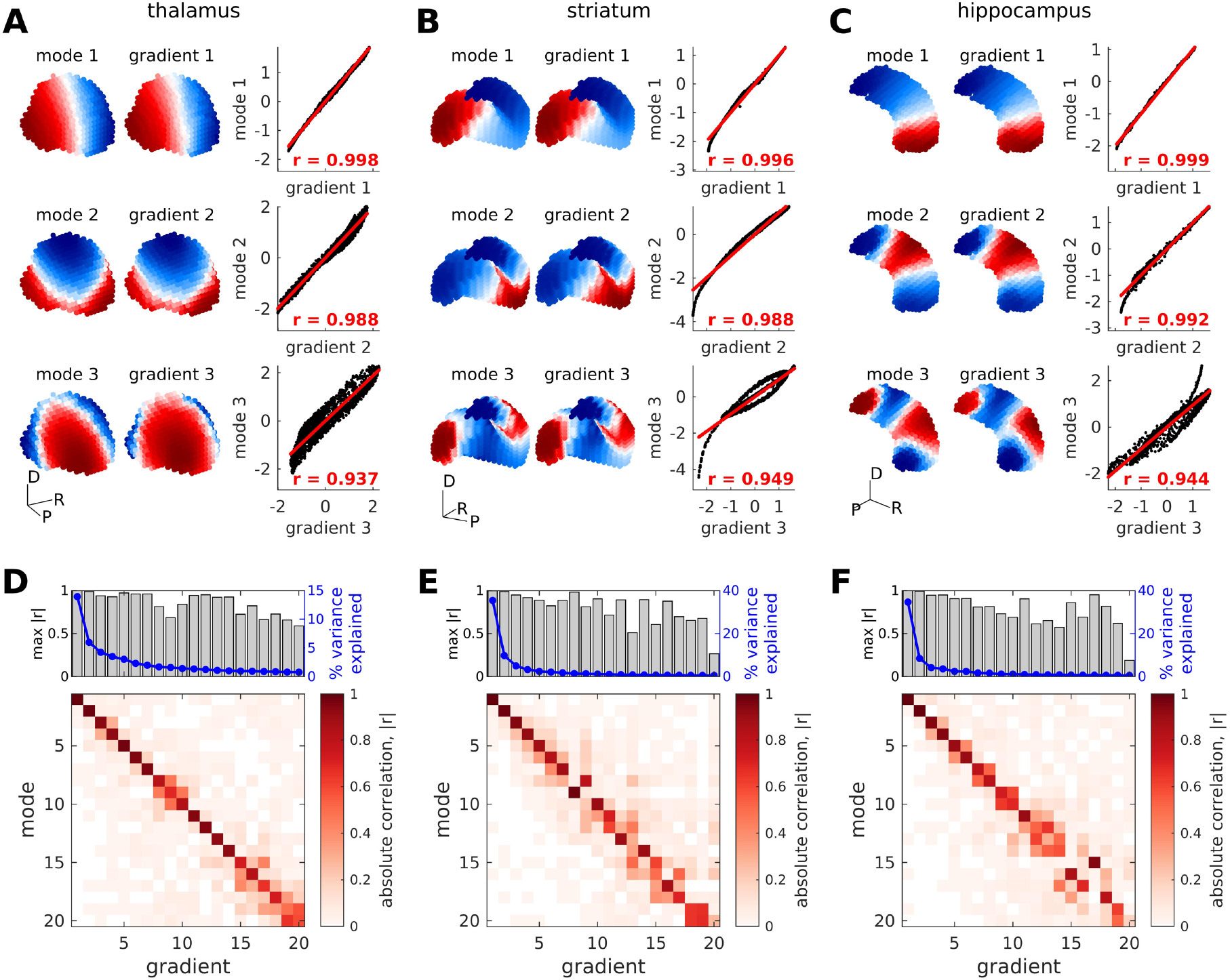
Geometric eigenmodes capture the functional organization of non-neocortical structures. (**A–C**) First three geometric eigenmodes and FC-based functional gradients in the thalamus, striatum, and hippocampus, respectively. The modes and gradients are shown in 3D coordinate space, with negative–zero–positive values colored as blue–white–red. The labels show dorsal (D), posterior (P), and rightward (R) directions. The scatter plots show the relationship between the modes and gradients, with the red lines representing linear fits with Pearson correlation coefficient *r*. (**D–F**) Absolute correlation (|*r*|) of the first 20 geometric eigenmodes and functional gradients in the thalamus, striatum, and hippocampus, respectively. The top panels show the highest |*r*| obtained by each functional gradient (gray bars), taking into account order flips in geometric eigenmodes (Supplementary Material-S14), and the percentage of variance explained by each functional gradient (blue lines).

## DISCUSSION

The dynamics of many physical systems are constrained by their geometry and can be understood as excitations of a relatively small number of excited structural modes ^6,7^. Here we show that structural eigenmodes derived solely from the geometry of the brain’s structure provide a more compact, accurate, and parsimonious representation of its macroscale activity than alternative connectome-based models. This mode-based view of the brain further reveals that spontaneous and evoked brain activity captured by fMRI is dominated by large-scale eigenmodes with relatively long wavelengths, whose dynamics are derived from a biophysically motivated wave equation. These findings challenge the classical neuroscientific paradigm, in which topologically complex patterns of inter-regional connectivity between discrete, specialized neuronal populations are viewed as a critical anatomical foundation for dynamics. Our findings further indicate that a physically grounded approach that treats the brain as a continuous, spatially embedded system offers a unifying framework for understanding structural constraints on diverse aspects of neuronal function.

The extensive comparisons of geometric eigenmodes with other anatomical (connectome and EDR eigenmodes) and mathematical (PCA and Fourier) basis sets show that their superior performance in capturing macroscale neocortical activity is not trivially driven by generic mathematical properties of basis set expansions. Rather, this result indicates that geometry represents a fundamental anatomical constraint on dynamics. Additionally, the strong performance of the EDR eigenmodes derived from a synthetic network suggests that a homogeneous, distance-dependent connectivity with near-exponential form represents another important anatomical constraint on activity. EDR-like connectivity is mathematically embedded in the Helmholtz equation in Eq. (1) ^25,30^, so the role of such connectivity is implicitly captured by the geometric eigenmodes. The comparatively poor performance of connectome eigenmodes indicates that topologically complex connections that exist beyond a simple EDR afford minimal further benefit in obtaining eigenmodes that can accurately explain spatiotemporal patterns of cortical activity, as measured with fMRI. Our findings thus counter traditional views that emphasize intricate patterns of anatomical connections as the primary driver of coordinated dynamics ^33,63,64^. Indeed, recent estimates indicate that long-range cortical connections are relatively rare ^65^; they may therefore represent a relatively minor perturbation of the dominant effect imposed by EDR-like connectivity. Nonetheless, the topological centrality, metabolic cost, and tight genetic control of such connections ^66–68^ suggest that they provide important functional and evolutionary advantages beyond wave-like dynamics ^69^. The limited resolution and sensitivity to preprocessing pipelines ^70,71^ of dMRI and fMRI data complicate attempts to uncover the role of long-range connections, but high-quality animal tract-tracing and electrophysiological data may be helpful in this regard.

The close coupling between geometry and dynamics is apparent in neocortical and non-neocortical structures alike, suggesting that the functional organization of regions outside the neocortex is also dominated by local anatomical connectivity and wave dynamics, as found in recent experiments ^46,72,73^. These observations indicate that geometric eigenmodes offer a simpler, more parsimonious, and mechanistically informative account of putative gradients of functional organization in non-neocortical structures than the complex manifold-learning procedures currently used in the literature ^74^. This is because such procedures are phenomenological, providing statistical descriptions of dominant sources of variances in the data, whereas the study of structural eigenmodes derives from a generative process. Notably, we do not observe the same one-to-one spatial correspondence between single geometric eigenmodes and previously described FC-derived functional gradients of the neocortex, the most dominant of which captures a hierarchical sensory-fugal axis of function (i.e., compare Fig. 1A with Fig. 1A in ^75^). Functional gradients of the neocortex may thus reflect a superposition of geometric modes ^76^, just as musical chords emerge from combinations of individual notes.

Geometric mode decomposition offers unique insights into the spatial properties of brain activation maps. Classical brain mapping analyses typically focus on responses in isolated clusters of spatial locations that exceed a statistical threshold ^77^. Our approach aligns with rigorously established results from physics and engineering, in which perturbations of spatially continuous systems elicit system-wide responses; for instance, the musical notes of a violin string result from oscillations across its entire length rather than the behaviour of an isolated string segment ^78^. Notably, the use of geometric eigenmodes indicates that, across >10,000 diverse maps from task-based fMRI studies, task engagement is associated predominantly with the excitation of modes with wavelengths of ∼60 mm and longer. This result coincides with similar observations of long-wavelength excitations in empirical EEG and evoked response potential (ERP) data ^39–41^ and suggests that classical analyses relying on thresholding of point-wise statistical maps obscure the spatially extended and complex patterns of activity actually evoked by a task.

Our modeling results offer insight into the physical processes underlying the close link observed between geometry and function. In particular, the relative simplicity and superior performance of the wave model in capturing diverse aspects of spontaneous fMRI dynamics indicates that it provides a more parsimonious account than a complex neural mass model that views the brain as a graph of discrete anatomical regions (nodes) coupled via the connectome (edges). This finding is consistent with experimental observations of wave dynamics in both human and animal fMRI data ^79,80^. Future work could explore whether introducing spatial heterogeneities ^81^ or complex structured input ^76^ into the wave model further improves its accuracy and explain more empirical phenomena.

Application of the wave model to mimic visual stimulation reveals that waves propagating from the stimulation site segregate along the classical dorsal and ventral visual pathways, and that regional responses to the perturbation conform to a well-described hierarchy of timescales ranging from rapidly responding unimodal areas to slower-responding transmodal regions ^53,56,57^. These canonical properties of hierarchical visual processing have been extensively studied for decades and are classically thought to be driven by complex patterns of layer-specific inter-regional connectivity ^52,54,55^, but our analysis shows that waves traveling through the cortical geometry are sufficient for the emergence of segregated, hierarchical processing streams. In other words, while our findings cannot rule out a role for complex inter-regional connectivity, but they do indicate that such connectivity is not necessary for the emergence of these macroscale dynamics.

The superior performance of geometric eigenmodes offers an immediate practical benefit, since the modes can be estimated using only a mesh representation of the structure of interest, which can easily be derived using well-established, automated processing pipelines for T1-weighted anatomical images ^82^. In contrast, connectome eigenmodes require a graph-based model of macroscopic inter-regional connectivity generated via complex data processing pipelines ^71,83^; the definition of graph nodes, which is a topic of contention ^84^; and the application of a thresholding procedure to remove putatively spurious connections, which our own analysis shows can affect the findings (Fig. S9). The fact that such choices are not required to obtain the geometric eigenmodes means that they can be applied robustly and flexibly across different experimental contexts in both humans and other species ^85,86^, opening new avenues of research. For example, one can investigate how geometric eigenmodes vary through neurodevelopment or are disrupted in clinical disorders. Indeed, the close link we identify between geometry and function implies that inter-species differences in spatiotemporal dynamics may largely be driven by differences in brain shape. Characterizing how variations in brain geometry, both within and between species, shape brain function will be essential for understanding physical and anatomical constraints on neuronal activity.

## METHODS

### Materials and methods

All details can be found in the Supplementary Materials.

### Data and code availability

Raw and preprocessed HCP data can be accessed at https://db.humanconnectome.org/. All source data and computer codes to calculate the eigenmodes, analyze results, and generate the main and supplementary figures of this study are openly available at https://github.com/BMHLab/BrainEigenmodes.

## ACKNOWLEDGMENTS

We thank Sina Mansour L for providing access to high-resolution connectome data and Koen Haak for assistance with connectopic mapping. HCP data were provided by the Human Connectome Project, Wu-Minn Consortium (Principal Investigators: David Van Essen and Kamil Ugurbil; 1U54MH091657) funded by the 16 NIH Institutes and Centers that support the NIH Blueprint for Neuroscience Research, and by the McDonnell Center for Systems Neuroscience at Washington University. This work was supported by the MASSIVE HPC facility (www.massive.og.au), Sylvia and Charles Viertel Foundation grant 2017042 to AF, National Health and Medical Research Council grants 1197431 and 1146292 to AF, Australian Research Council grant DP200103509 to AF, National Health and Medical Research Council grant 2008612 to MB, Australian Research Council Laureate Fellowship FL140100025 to PAR, and Australian Research Council Center of Excellence CE140100007 to PAR.

## AUTHOR CONTRIBUTIONS

KMA and AF conceptualized the study. JCP, KMA, AF, and MO designed the methodology. JCP, KMA, and AF performed the investigation and administered the project. JCP and KMA developed the visualizations. AF acquired funding. AF, PAR, BF, and MB supervised the project. JCP and AD wrote the original draft. All authors reviewed and edited the final manuscript.

## REFERENCES

1. Molkenthin, N. & Timme, M. Scaling Laws in Spatial Network Formation. Physical Review Letters 117, 168301 (2016).

2. Yang, Y., Wang, W., Lou, Y., Yin, J. & Gong, X. Geometric and amino acid type determinants for protein-protein interaction interfaces. Quantitative Biology 6, 163–174 (2018).

3. Nunez, P. L. Neocortical Dynamics and Human EEG Rhythms. (Oxford University Press, 1995).

4. Braitenberg, V. & Schüz, A. Cortex: Statistics and Geometry of Neuronal Connectivity. Cortex: Statistics and Geometry of Neuronal Connectivity (1998). doi:10.1007/978-3-662-03733-1.

5. Damoiseaux, J. S. & Greicius, M. D. Greater than the sum of its parts: A review of studies combining structural connectivity and resting-state functional connectivity. Brain Structure and Function 213, 525–533 (2009).

6. Melrose, D. B. & McPhedran, R. C. Electromagnetic Processes in Dispersive Media. (Cambridge University Press, 1991).

7. Goldstein, H. Classical Mechanics: Pearson New International Edition. (Pearson, 2013).

8. Nozari, E. et al. Is the brain macroscopically linear? A system identification of resting state dynamics. arXiv (2020).

9. Jones, E. G. Golgi, Cajal and the Neuron Doctrine. Journal of the History of the Neurosciences 8, 170–178 (1999).

10. Brodmann, K. Beiträge zur histologischen Lokalisation der Grosshirnrinde, VI: Mitteilung: Die Cortexgliederung des Menschen. J Psychologie Neurologie 10, 231–246 (1908).

11. Broca, P. Remarques sur le siège de la faculté du langage articulé, suivies d’une observation d’aphémie (perte de la parole). Bulletin et mémoires de la Société Anatomique de Paris 6, 330–357 (1861).

12. Vaidya, A. R., Pujara, M. S., Petrides, M., Murray, E. A. & Fellows, L. K. Lesion Studies in Contemporary Neuroscience. Trends Cogn Sci 23, 653–671 (2019).

13. Bullmore, E. & Sporns, O. Complex brain networks: Graph theoretical analysis of structural and functional systems. Nature Reviews Neuroscience 10, 186–198 (2009).

14. Yuste, R. From the neuron doctrine to neural networks. Nature Reviews Neuroscience 16, 487–497 (2015).

15. Fornito, A., Zalesky, A. & Bullmore, E. T. Fundamentals of Brain Network Analysis. (2016).

16. Breakspear, M. Dynamic models of large-scale brain activity. Nature Neuroscience 20, 340–352 (2017).

17. Atasoy, S., Donnelly, I. & Pearson, J. Human brain networks function in connectome-specific harmonic waves. Nature Communications 7, 10340 (2016).

18. Preti, M. G. & Van De Ville, D. Decoupling of brain function from structure reveals regional behavioral specialization in humans. Nature Communications 10, 4747 (2019).

19. Rué-Queralt, J. et al. The connectome spectrum as a canonical basis for a sparse representation of fast brain activity. NeuroImage 244, 118611 (2021).

20. Beurle, R. L. Properties of a mass of cells capable of regenerating pulses. Philosophical Transactions of the Royal Society of London. Series B, Biological Sciences 240, 55–94 (1956).

21. Lopes da Silva, F. H., van Rotterdam, A., Barts, P., van Heusden, E. & Burr, W. Models of Neuronal Populations: The Basic Mechanisms of Rhythmicity. Progress in Brain Research 45, 281–308 (1976).

22. Wright, J. J. & Liley, D. T. J. Simulation of electrocortical waves. Biological Cybernetics 72, 347–356 (1995).

23. Deco, G., Jirsa, V. K., Robinson, P. A., Breakspear, M. & Friston, K. The dynamic brain: From spiking neurons to neural masses and cortical fields. PLoS Computational Biology 4, (2008).

24. Jirsa, V. & Haken, H. Field Theory of Electromagnetic Brain Activity. Physical Review Letters 77, 960–963 (1996).

25. Robinson, P. A., Rennie, C. J. & Wright, J. J. Propagation and stability of waves of electrical activity in the cerebral cortex. Physical Review E 56, 826–840 (1997).

26. Robinson, P. A., Rennie, C. J., Rowe, D. L., O’Connor, S. C. & Gordon, E. Multiscale brain modelling. Philosophical Transactions of the Royal Society B: Biological Sciences 360, 1043–1050 (2005).

27. Robinson, P. A. Physical brain connectomics. Physical Review E 99, 012421 (2019).

28. Wang, X. J. & Kennedy, H. Brain structure and dynamics across scales: In search of rules. Current Opinion in Neurobiology 37, 92–98 (2016).

29. Roberts, J. A., Perry, A., Roberts, G., Mitchell, P. B. & Breakspear, M. Consistency-based thresholding of the human connectome. NeuroImage 145, 118–129 (2017).

30. Robinson, P. A. et al. Eigenmodes of brain activity: Neural field theory predictions and comparison with experiment. NeuroImage 142, 79–98 (2016).

31. Gabay, N. C. & Robinson, P. A. Cortical geometry as a determinant of brain activity eigenmodes: Neural field analysis. Physical Review E 96, (2017).

32. Gabay, N. C., Babaie-Janvier, T. & Robinson, P. A. Dynamics of cortical activity eigenmodes including standing, traveling, and rotating waves. Physical Review E 98, 042413 (2018).

33. Honey, C. J., Kötter, R., Breakspear, M. & Sporns, O. Network structure of cerebral cortex shapes functional connectivity on multiple time scales. Proceedings of the National Academy of Sciences of the United States of America 104, 10240–10245 (2007).

34. Fischl, B., Sereno, M. I., Tootell, R. B. H. & Dale, A. M. High-resolution intersubject averaging and a coordinate system for the cortical surface. Human Brain Mapping 8, 272–284 (1999).

35. Reuter, M., Wolter, F. E. & Peinecke, N. Laplace-Beltrami spectra as ‘Shape-DNA’ of surfaces and solids. CAD Computer Aided Design 38, 342–366 (2006).

36. van Essen, D. C. et al. The WU-Minn Human Connectome Project: An overview. NeuroImage 80, 62–79 (2013).

37. Glasser, M. F. et al. A multi-modal parcellation of human cerebral cortex. Nature 536, 171–178 (2016).

38. Naze, S., Proix, T., Atasoy, S. & Kozloski, J. R. Robustness of connectome harmonics to local gray matter and long-range white matter connectivity changes: Sensitivity analysis of Connectome Harmonics. NeuroImage 224, 117364 (2021).

39. Robinson, P. A., Loxley, P. N., O’Connor, S. C. & Rennie, C. J. Modal analysis of corticothalamic dynamics, electroencephalographic spectra, and evoked potentials. Physical Review E 63, (2001).

40. Wingeier, B. M., Nunez, P. L. & Silberstein, R. B. Spherical harmonic decomposition applied to spatial-temporal analysis of human high-density electroencephalogram. Physical Review E 64, (2001).

41. Mukta, K. N., Robinson, P. A., Pagès, J. C., Gabay, N. C. & Gao, X. Evoked response activity eigenmode analysis in a convoluted cortex via neural field theory. Physical Review E 102, (2020).

42. Barch, D. M. et al. Function in the human connectome: Task-fMRI and individual differences in behavior. NeuroImage 80, 169–189 (2013).

43. Gorgolewski, K. J. et al. NeuroVault.Org: A web-based repository for collecting and sharing unthresholded statistical maps of the human brain. Frontiers in Neuroinformatics 9, (2015).

44. Lévy, B. Laplace-beltrami eigenfunctions towards an algorithm that ‘understands’ geometry. in Proceedings - IEEE International Conference on Shape Modeling and Applications 2006, SMI 2006 vol. 2006 13 (2006).

45. Robinson, P. A. et al. Prediction of electroencephalographic spectra from neurophysiology. Physical Review E 63, 021903 (2001).

46. Raut, R. V. et al. Global waves synchronize the brain’s functional systems with fluctuating arousal. Science Advances 7, (2021).

47. Bolt, T. et al. A parsimonious description of global functional brain organization in three spatiotemporal patterns. Nature Neuroscience 25, 1093–1103 (2022).

48. Sanz-Leon, P. et al. NFTsim: Theory and Simulation of Multiscale Neural Field Dynamics. PLoS Computational Biology 14, e1006387 (2018).

49. Deco, G. et al. How local excitation-inhibition ratio impacts the whole brain dynamics. Journal of Neuroscience 34, 7886–7898 (2014).

50. Demirtaş, M. et al. Hierarchical Heterogeneity across Human Cortex Shapes Large-Scale Neural Dynamics. Neuron 101, 1181–1194 (2019).

51. Mitra, A., Snyder, A. Z., Blazey, T. & Raichle, M. E. Lag threads organize the brain’s intrinsic activity. Proceedings of the National Academy of Sciences 112, E2235–E2244 (2015).

52. Felleman, D. J. & Van Essen, D. C. Distributed hierarchical processing in the primate cerebral cortex. Cerebral Cortex 1, 1–47 (1991).

53. Chaudhuri, R., Knoblauch, K., Gariel, M. A., Kennedy, H. & Wang, X. J. A Large-Scale Circuit Mechanism for Hierarchical Dynamical Processing in the Primate Cortex. Neuron 88, 419–431 (2015).

54. Goodale, M. A. & Milner, A. D. Separate visual pathways for perception and action. Trends in Neurosciences 15, 20–25 (1992).

55. Markov, N. T. et al. Anatomy of hierarchy: Feedforward and feedback pathways in macaque visual cortex. Journal of Comparative Neurology 522, 225–259 (2014).

56. Hasson, U., Yang, E., Vallines, I., Heeger, D. J. & Rubin, N. A hierarchy of temporal receptive windows in human cortex. Journal of Neuroscience 28, 2539–2550 (2008).

57. Murray, J. D. et al. A hierarchy of intrinsic timescales across primate cortex. Nature Neuroscience 17, 1661–1663 (2014).

58. Glasser, M. F. & Van Essen, D. C. Mapping Human Cortical Areas In Vivo Based on Myelin Content as Revealed by T1-and T2-Weighted MRI. Journal of Neuroscience 31, 11597–11616 (2011).

59. Gao, R., Van den Brink, R. L., Pfeffer, T. & Voytek, B. Neuronal timescales are functionally dynamic and shaped by cortical microarchitecture. eLife 9, e61277 (2020).

60. Tian, Y., Margulies, D. S., Breakspear, M. & Zalesky, A. Topographic organization of the human subcortex unveiled with functional connectivity gradients. Nature Neuroscience 23, 1421–1432 (2020).

61. Wachinger, C., Golland, P., Kremen, W., Fischl, B. & Reuter, M. BrainPrint: A discriminative characterization of brain morphology. NeuroImage 109, 232–248 (2015).

62. Haak, K. V., Marquand, A. F. & Beckmann, C. F. Connectopic mapping with resting-state fMRI. NeuroImage 170, 83–94 (2018).

63. Deco, G., Jirsa, V. K. & McIntosh, A. R. Emerging concepts for the dynamical organization of resting-state activity in the brain. Nature Reviews Neuroscience 12, 43–56 (2011).

64. Pang, J. C., Gollo, L. L. & Roberts, J. A. Stochastic synchronization of dynamics on the human connectome. NeuroImage 229, 117738 (2021).

65. Rosen, B. Q. & Halgren, E. An estimation of the absolute number of axons indicates that human cortical areas are sparsely connected. PLoS Biology 20, e3001575 (2022).

66. van den Heuvel, M. P., Kahn, R. S., Goñi, J. & Sporns, O. High-cost, high-capacity backbone for global brain communication. Proceedings of the National Academy of Sciences of the United States of America 109, 11372–11377 (2012).

67. Arnatkeviciute, A. et al. Genetic influences on hub connectivity of the human connectome. Nature Communications 12, (2021).

68. Rosen, B. Q. & Halgren, E. An estimation of the absolute number of axons indicates that human cortical areas are sparsely connected. PLoS Biology 20, e3001575 (2022).

69. Pang, J. C., Rilling, J. K., Roberts, J. A., van den Heuvel, M. P. & Cocchi, L. Evolutionary shaping of human brain dynamics. eLife 11, e80627 (2022).

70. Aquino, K. M., Fulcher, B. D., Parkes, L., Sabaroedin, K. & Fornito, A. Identifying and removing widespread signal deflections from fMRI data: Rethinking the global signal regression problem. NeuroImage 212, 116614 (2020).

71. Gajwani, M. et al. Can hubs of the human connectome be identified consistently with diffusion MRI? 2022.12.21.521366 Preprint at https://doi.org/10.1101/2022.12.21.521366 (2022).

72. Hamid, A. A., Frank, M. J. & Moore, C. I. Wave-like dopamine dynamics as a mechanism for spatiotemporal credit assignment. Cell 184, 2733-2749.e16 (2021).

73. Yousefi, B. & Keilholz, S. Propagating patterns of intrinsic activity along macroscale gradients coordinate functional connections across the whole brain. NeuroImage 231, 117827 (2021).

74. Haak, K. V., Marquand, A. F. & Beckmann, C. F. Connectopic mapping with resting-state fMRI. NeuroImage 170, 83–94 (2018).

75. Margulies, D. S. et al. Situating the default-mode network along a principal gradient of macroscale cortical organization. Proceedings of the National Academy of Sciences of the United States of America 113, 12574–12579 (2016).

76. Henderson, J. A., Aquino, K. M. & Robinson, P. A. Empirical estimation of the eigenmodes of macroscale cortical dynamics: Reconciling neural field eigenmodes and resting-state networks. Neuroimage: Reports 2, 100103 (2022).

77. Coalson, T. S., Van Essen, D. C. & Glasser, M. F. The impact of traditional neuroimaging methods on the spatial localization of cortical areas. Proceedings of the National Academy of Sciences of the United States of America 115, E6356–E6365 (2018).

78. Robinson, P. A. et al. Determination of Dynamic Brain Connectivity via Spectral Analysis. Frontiers in Human Neuroscience 15, (2021).

79. Majeed, W. et al. Spatiotemporal dynamics of low frequency BOLD fluctuations in rats and humans. NeuroImage 54, 1140–1150 (2011).

80. Matsui, T., Murakami, T. & Ohki, K. Transient neuronal coactivations embedded in globally propagating waves underlie resting-state functional connectivity. Proceedings of the National Academy of Sciences 113, 6556–6561 (2016).

81. Deco, G. et al. Dynamical consequences of regional heterogeneity in the brain’s transcriptional landscape. Science Advances 7, eabf4752 (2021).

82. Fischl, B. FreeSurfer. NeuroImage 62, 774–781 (2012).

83. Oldham, S. et al. The efficacy of different preprocessing steps in reducing motion-related confounds in diffusion MRI connectomics. NeuroImage 222, 117252 (2020).

84. Arslan, S. et al. Human brain mapping: A systematic comparison of parcellation methods for the human cerebral cortex. NeuroImage 170, 5–30 (2018).

85. Tokariev, A. et al. Large-scale brain modes reorganize between infant sleep states and carry prognostic information for preterms. Nature Communications 10, 2619 (2019).

86. Chen, Y.-C. et al. The individuality of shape asymmetries of the human cerebral cortex. eLife 11, e75056 (2022).

