## Supplementary Materials for "Geometric constraints on human brain function"

### MATERIALS AND METHODS

#### S1. Neural field theory

Neural field theory (NFT) is a class of biophysical models that explain mesoscale to macroscale brain dynamics (from about 0.5 mm to whole brain) as the outcome of spatially extended, time-varying fields of neural activity [1–7]. General NFTs focus on local average dynamics of neural populations, such as mean firing rates and soma voltages. Important biophysical processes, such as dendritic and synaptic processes, axonal conduction delays, and summation of dendritic current in the cell soma, are incorporated as local population averages through a series of principled mathematical reductions.

NFTs treat neural tissue at scales above ~0.5 mm as a spatial continuum with local properties and point-to-point white-matter connectivity that decreases smoothly with distance. Although more general cases can be treated via NFT, including interactions with subcortical structures [7,8], the simplest versions include only the cortex and assume that its connectivity is homogeneous and isotropic, with the connection strength only depending on the distance between points and typically decreasing approximately exponentially with their separation [4,9–11]. This relation is especially relevant in the context of functional MRI (fMRI), where dynamics are relatively slow and the strength of active connections has a spatial dependence that is approximately exponential [9,11,12]. The resulting spatiotemporal evolution of the activity fields has been effectively described by a physiologically constrained variant of NFT developed by Robinson and colleagues [1,2,5,6,13] as comprising damped waves propagating across the cortical sheet after being excited by external inputs and local cortical or corticothalamic dynamics. Over the last two decades, this NFT formulation has successfully explained and unified a diverse range of experimental phenomena in a unified way, including but not limited to electroencephalography (EEG) spectra [14,15], evoked potentials [16,17], arousal states [18,19], whole-brain effective connectivity [12], cortical activity waves [20], and sleep state reorganizations [21]. Because mesoscale to macroscale brain dynamics are approximately operating in the linear regime under normal conditions (excluding seizure-like dynamics) [22], the Robinson et al. NFT has shown that spatial eigenmodes (modes) of activity naturally emerge in the brain [23,24], with their properties shaped by the brain's intrinsic geometry and topology [20,25]. These modes are the topic of interest in this study. We refer the readers to the extensive literature of NFTs for a more detailed discussion (see for example [4–8,23,26–29] and the references cited therein).

#### S2. Derivation of cortical geometric eigenmodes

If brain structure can be approximated as being constant in time, the resulting spatial and temporal dynamics can be treated separately via eigenmode decomposition [23,26], similar to the treatment of other physical systems [30]. In particular, the spatial aspect satisfies the Laplacian eigenvalue problem, which is also known as the Helmholtz equation,

$$\nabla^2\psi = \Delta\psi = -\lambda\psi, \qquad (S1)$$

where $\nabla$ is the gradient operator, $\Delta$ is the Laplace-Beltrami Operator (LBO), and $\psi = \{\psi_1(\boldsymbol{r}), \psi_2(\boldsymbol{r}), \ldots\}$ is the family of real-valued geometric eigenmodes with corresponding family of eigenvalues, $\lambda = \{\lambda_1, \lambda_2, \ldots\}$.

For the brain, the LBO captures intrinsic geometry, which includes the curvature of the cortical surface [31], and is defined generally as [32,33],

$$\Delta \coloneqq \frac{1}{W} \sum_{i,j} \frac{\partial}{\partial x_i} \left( g^{ij} W \frac{\partial}{\partial x_j} \right), \qquad (S2)$$

where $x_i, x_j$ are the local coordinates, $g^{ij}$ is the inverse of the inner product metric tensor $g_{ij} \coloneqq \langle \frac{\partial}{\partial x_i}, \frac{\partial}{\partial x_j} \rangle$, $W \coloneqq \sqrt{\det G}$, det denotes the determinant, and $G \coloneqq (g_{ij})$.

We employed the LaPy python library [31,34] installed on the MASSIVE High Performance Computing facility [35] to derive the geometric eigenmodes of the human cortex by using a triangular surface mesh representation of the midthickness human cortical surface, comprising 32,492 vertices in each hemisphere, obtained from a downsampled, left-right symmetric version of the FreeSurfer's fsaverage population-averaged template [36] (https://github.com/ThomasYeoLab/CBIG/tree/master/data/templates/surface/fs_LR_32k). This template is independent of the data sample used in all our analyses, thus obviating any concerns about circularity. Note that the continuous LBO operates on the underlying Riemannian manifold of the surface and not directly on mesh vertices. LaPy uses the cubic finite element method on the surface mesh to achieve numerically tractable solutions of Eq. (S1) on an interpolated smooth manifold. This distinguishes it from the discrete graph Laplacian [37], which does not encode spatial relations between points. All our analyses were focused on unihemispheric eigenmodes, but our approach can easily be extended to the whole brain because bihemispheric eigenmodes can be represented as symmetric or antisymmetric combinations of the eigenmodes derived from each hemisphere [23]; symmetric corresponds to mirror-symmetry across the sagittal midplane and asymmetric corresponds to hemispheres having the same spatial structure but with flipped signs.

The eigenvalue solutions of Eq. (S1) are ordered sequentially according to the spatial frequency or wavelength of the spatial patterns of each eigenmode; i.e., $0 \leq \lambda_1 \leq \lambda_2 \leq \cdots$. Note that the first eigenvalue $\lambda_1$ is approximately equal to zero (wavelength $\gg$ size of the brain) and the corresponding eigenmode $\psi_1$ is a constant function with no nodal lines (zero sets of the function). Throughout our study, we used the first 200 modes (including the constant mode $\psi_1$) in our analyses given the diminishing improvements in reconstruction accuracy observed when using an increasing number of modes (Fig. 1D).

Eq. (S1) can be used to solve the eigenmodes of any cortical surface mesh model, but changes in the geometry of the mesh can alter the resulting eigenvalues and eigenmodes. Hence, the eigenvalues and eigenmodes derived from individual subject surfaces, particularly at very short wavelengths, will differ and cannot be straightforwardly compared [38,39]. For simplicity, here we used the eigenmodes generated from a common template surface, as discussed above. While this allows for comparison of data reconstructions in different individuals, it obscures any potential effects associated with individual differences in cortical geometry. However, Fig. S4 shows that using geometric eigenmodes derived from individual cortical surfaces does not change the general results of the study, suggesting that, for present purposes, the eigenmodes derived from the population-averaged template surface represent a good approximation of fundamental geometric eigenmodes. Nonetheless, to highlight certain nuances of individual geometry, we show in Fig. S5 that 200 individual-specific eigenmodes perform slightly better than template-derived eigenmodes in some individuals, especially in reconstructing task-activation maps but not in reconstructing resting-state activity. However, Fig. S6 shows that the results for individual-specific and template-derived eigenmodes converge at very short wavelengths (~500$^{th}$ mode). Thus, the effects of individual differences in cortical geometry on brain function are captured by the first 200 modes, consistent with recent work [39], which corresponds to a very small fraction of the maximum possible number of eigenmodes (~0.6%).

Each eigenmode comprises spatial patterns with specific spatial wavelength. Following [23], we approximate the wavelengths using an idealized spherical case as it is topologically comparable to the human cortex. By solving Eq. (S1) on a sphere, degenerate solutions exist, such that certain eigenmodes have the same eigenvalue and spatial wavelength; this is analogous to spherical harmonics in quantum physics. In fact, the eigenmodes will approach the spherical harmonics in the limit of vanishing cortical folding [23]. Hence, the eigenmodes can be grouped together into an eigengroup with spatial wavelength [39],

$$wavelength = \frac{2\pi R_s}{\sqrt{l(l+1)}}, \qquad (S3)$$

where $R_s$ is the radius of the sphere (for the fsaverage population-averaged template used in this study, $R_s \approx 67.0$ mm) and $l$ is the eigengroup number (the angular momentum quantum number in atomic physics). The wavelengths of the first 15 eigengroups and the eigenmodes included in the eigengroup are shown in Table S1.

**Table S1. Spatial wavelengths of the eigenmodes of a sphere.**

| Eigengroup | Wavelength (mm) | Eigenmodes included in the eigengroup |
|---|---|---|
| 0 | – | 1 |
| 1 | 297.7 | 2–4 |
| 2 | 171.9 | 5–9 |
| 3 | 121.5 | 10–16 |
| 4 | 94.1 | 17–25 |
| 5 | 76.9 | 26–36 |
| 6 | 65.0 | 37–49 |
| 7 | 56.3 | 50–64 |
| 8 | 49.6 | 65–81 |
| 9 | 44.4 | 82–100 |
| 10 | 40.1 | 101–121 |
| 11 | 36.6 | 122–144 |
| 12 | 33.7 | 145–169 |
| 13 | 31.2 | 170–196 |
| 14 | 29.1 | 197–225 |

<u>S3. Mode decomposition of brain activity</u>

We used the geometric eigenmodes to decompose spatiotemporal fMRI data, measured at spatial location $\boldsymbol{r}$ and time $t$, for each individual as a weighted sum of modes,

$$y(\boldsymbol{r},t)=\sum_{j=1}^{N}a_j(t)\psi_j(\boldsymbol{r}), \qquad (S4)$$

where $a_j$ is the amplitude of mode $j$ in explaining the data, $\psi_j$ is the $j$th mode, and $N$ is the number of modes used. As outlined in Section S2, $N = 200$ for our analyses. For spatiotemporal data—i.e., recordings of spontaneous dynamics from task-free fMRI—each time frame of the data was substituted into Eq. (S4), resulting in a time-dependent amplitude $a_j(t)$ for each mode $\psi_j$. For purely spatial data—i.e., task-evoked activation maps—the amplitudes are independent of time such that $a_j(t) \to a_j$. In both cases, the amplitudes can be obtained by integrating over the cortical surface,

$$a_j(t)=\int y(\boldsymbol{r},t)\psi_j(\boldsymbol{r})d\boldsymbol{r}, \qquad (S5)$$

which can be derived from Eq. (S4) by using the orthogonal property of the eigenmodes [24,40]. If there are insufficient measurements to evaluate the integral, the amplitudes can also be estimated via a statistical general linear model.

After obtaining the amplitudes, Eq. (S4) is used to calculate the reconstructed data. We then quantified the accuracy of this reconstruction by calculating the correlation between the empirical and reconstructed data. For the spatiotemporal task-free data, we first parcellated the empirical and reconstructed data by taking the average of the data within discrete parcels/regions (Section S5) as per standard practice in the field [41], and then constructed a matrix of inter-regional functional coupling (FC) by calculating the Pearson correlation coefficient of pairs of parcel time series. For the task-evoked data, we applied the same parcellation on the activation maps to allow direct comparison. Finally, the reconstruction accuracy for the task-free data was calculated by taking the correlation of the upper triangular elements of the empirical and reconstructed FCs. For the task-evoked data, the reconstruction accuracy was calculated from the spatial correlation of parcellated empirical and reconstructed maps. We then took the average reconstruction accuracy across all participants.

### S4. Human Connectome Project data

#### *S4.1 – Participants*

We used preprocessed functional magnetic resonance imaging (fMRI) data from the Human Connectome Project (HCP) [42]. We did not perform any additional preprocessing steps, such as global signal removal (GSR), because the first eigenmode (considered as the global, constant mode) already explicitly captures global deviations in the data, allowing the other modes to capture functionally relevant non-global activity. We analyzed data from 255 unrelated healthy individuals (ages 22–35; 132 females), which is the largest HCP sample excluding twins or siblings and with all participants having completed task-evoked and task-free resting-state data. Procedures were carried out in accordance with protocols set by the HCP's data use terms. For a detailed account of the image acquisition protocol and preprocessing pipelines, see [42,43].

*S4.2 – Task-evoked data*

We analyzed task-evoked fMRI data measured in 7 task domains that have been shown to reliably recruit a wide array of neural systems [43]. The 7 tasks were: social, motor, gambling, working memory (wm), language, emotion, and relational. Table S2 shows the specific contrasts involved in each task domain and the key contrast investigated in this study. In total, we analyzed 47 contrasts, which include the 7 key contrasts. The key contrasts represent those used commonly in the literature to map the major activation pattern elicited by the task. See [43] for details about each task and contrast. The analysis was performed on individual task-activation maps computed via FSL's (https://fsl.fmrib.ox.ac.uk/) cross-run (Level 2) FEAT analysis [44]. We used the task maps as provided by HCP with minimal smoothing (2 mm) mapped onto the fsLR-32k CIFTI space, with 32,492 vertices in each hemisphere, via multimodal surface matching [45].

**Table S2. HCP task contrasts.**

| Task type | Number of contrasts | Contrasts | Key contrast |
|---|---|---|---|
| social | 3 | random; tom; tom_random | tom_random |
| motor | 13 | cue; lf; lh; rf; rh; t; avg; lf_avg; lh_avg; rf_avg; rh_avg; t_avg; cue_avg | cue_avg |
| gambling | 3 | punish; reward; punish_reward | punish_reward |
| working memory (wm) | 19 | 2bk_body; 2bk_face; 2bk_place; 2bk_tool; 0bk_body; 0bk_face; 0bk_place; 0bk_tool; 2bk; 0bk; body; face; place; tool; body_avg; face_avg; place_avg; tool_avg; 2bk_0bk | 2bk_0bk |
| language | 3 | math; story; math_story | math_story |
| emotion | 3 | faces; shapes; faces_shapes | faces_shapes |
| relational | 3 | match; rel; match_rel | match_rel |

*S4.3 – Task-free resting-state data*

We analyzed task-free resting-state fMRI acquired in one scanning session using a left to right (LR) encoding direction. The scan lasted for 14.4 min with a total of 1200 time frames. In brief, the resting-state fMRI acquisition parameters were: isotropic voxel size of 2 mm, repetition time (TR) of 720 ms, and echo time (TE) of 33.1 ms. All other acquisition parameters can be found in [42]. Each individual's data were preprocessed by HCP via their minimal preprocessing pipeline [46] and were also subjected to ICA-FIX to correct for structured noise and residual confounds [47]. No additional smoothing was performed. Similar to the task-evoked data, the resting-state data were mapped onto the fsLR-32k CIFTI space. Hence, each individual's data were represented as a matrix of size 32,492 vertices × 1200 time frames on each hemisphere.

*S4.4 – Connectome data*

To derive the connectome eigenmodes, we used individual connectomes derived from diffusion MRI (dMRI) data via probabilistic tractography, as provided in [48]. In brief, the dMRI acquisition parameters were: isotropic voxel size of 1.25 mm, TR of 5520 ms, TE of 89.5 ms, b-weightings of 1000, 2000, 3000 s/mm$^2$, and 6 b0-scans. All other acquisition parameters can be found in [42]. Each individual's data were preprocessed by HCP via their diffusion preprocessing pipeline (v3.19.0) [46]. To generate the connectomes, tractograms were generated using MRtrix with probabilistic tractography, 5 million streamlines that connect anatomically distinct brain regions, multi-shell multi-tissue (MSMT) constrained spherical deconvolution (CSD) anatomically constrained tractography (ACT), and 2nd-order Integration over Fiber Orientation Distributions algorithm (iFOD2). See [48] for further details. The fsLR-32k cortical surface mesh in standard MNI space was used to define the gray-matter–white-matter interface. For each individual, streamlines generated in each hemisphere were mapped to the closest vertices on the surface mesh to construct a high-resolution weighted connectome (32,492 × 32,492 matrix size and weights representing the total number of streamlines). See [48] for further details.

S5. Cortical parcellations

The main results of the study analyzed task-evoked activation maps and task-free FC data parcellated into discrete regions. In the main text, we present results using the HCP-MMP1 parcellation with 180 regions per hemisphere (we term this as the Glasser360 parcellation in the paper figures), which reflects sharp areal boundaries based on the combination of cortical architecture, function, connectivity, and topography [49]. The parcellation was on the fsLR-32k space, as provided by HCP. To test the robustness of our results (Figs. S3, S7, and S8), we also performed our analysis using parcellations provided by Schaefer et al. [50] on the fsLR-32k space with varying resolutions (100, 200, 400, 600, 800, and 1000 parcels across both hemispheres; we refer to these as the Schaefer100, Schaefer200, Schaefer400, Schaefer600, Schaefer800, and Schaefer1000 parcellations in the paper figures).

S6. Derivation of connectome eigenmodes

Connectome eigenmodes were derived as per prior methods [51] to enable comparison with previous findings. Note that in previous work [51–54], connectome eigenmodes have been referred to as connectome harmonics, but we use eigenmodes here since the term harmonics implies integer frequency ratios, which is not necessarily guaranteed.

We obtained high-resolution maps of connectivity measured with dMRI tractography as described in [48], in which the connectivity of each of the 32,492 vertices in the cortical surface mesh was estimated by tracing streamlines from each point until they terminated at some other point. Connection weights between vertices (considered as nodes) were estimated as the number of interconnecting streamlines without the need for normalization [55]. The tractography was performed on individuals from HCP (see Sections S4.1 and S4.4 for further details about the data and tractography method). From the tractography data, we combined the individual weighted connectivity matrices of size 32,492 × 32,492 to generate a group-averaged connectome $W_{\text{connectome}}$, with weights representing the average number of streamlines. We then generated a binary adjacency matrix, $A_{\text{local}}$, that captures the discrete representation of local spatial relations between points in the cortical surface mesh model constructed by connecting two vertices that are direct neighbors in the mesh. These links are intended to capture local, very short-range connectivity that cannot be resolved by traditional dMRI tractography [51].

Following [51], the group-averaged weighted connectome, $W_{\text{connectome}}$, was thresholded to remove the smallest weights, such that the number of connections was four times greater than $A_{\text{local}}$. The resulting thresholded matrix was binarized to obtain the group adjacency matrix, $A_{\text{connectome}}$. Finally, we generated a merged adjacency matrix $A_C = A_{\text{local}} \,||\, A_{\text{connectome}}$ (with matrix of size 32,492 × 32,492; $||$ is the OR operator), which captures both local, vertex-to-vertex connectivity and complex short- and long-range connections as measured empirically.

The connectome eigenmodes were obtained by solving the eigenvalue problem,

$$L'\psi = -\lambda\psi, \qquad (S6)$$

where $L'$ is the normalized graph Laplacian, a discrete counterpart of the LBO. The normalized graph Laplacian is related to the unnormalized graph Laplacian, $L$, as $L' = D^{-1/2} L D^{-1/2}$, with $L$ defined as per prior work [56],

$$L = \frac{1}{2}[(D - A_C) + (D - A_C)^T], \qquad (S7)$$

where $D$ is the diagonal degree matrix and the superscript $T$ denotes the matrix transpose. As with geometric eigenmodes, the eigenvalue solutions of Eq. (S6) form the sequence $0 \leq \lambda_1 \leq \lambda_2 \leq \cdots$. Note also that the use of a high-resolution, vertex-level connectome results in connectome eigenmodes spanning a space with dimensions (number of modes) equal to the number of vertices, allowing fair comparison with geometric eigenmodes.

Note that the connection density of the adjacency matrix, $A_C$, that resulted from the thresholding process described above was 0.10%. Since many network properties depend on network connection densities, and our specific thresholding procedure is somewhat arbitrary, we also derived connectome eigenmodes for variants of $A_C$ mapped across a range of thresholds, from 1% to 13% (i.e., the density of the unthresholded connectome) in increments of 2%. The results in Fig. S9 show that connection density did indeed affect the reconstruction accuracy of connectome eigenmodes, such that sparser matrices were associated with improved reconstructions. Given that long-range connections tend to have lower weights and are thus more likely to be thresholded away first, the application of more conservative thresholds emphasizes local, exponential distance rule (EDR)-like connectivity until finally converging on the surface mesh alone. These results thus align with the view that reconstruction performance is generally improved when focusing mostly on local connectivity. Notably, finely sampling connection densities from 0.01% to 5% reveals a putatively optimal threshold between 0.4% and 0.7% that results in peak reconstruction accuracy for 200 connectome eigenmodes. This optimal threshold may represent a physiologically plausible trade-off between sensitivity and specificity in tract reconstruction with diffusion MRI. However, the reconstruction accuracy never surpasses the accuracy of the geometric eigenmodes. The sensitivity of the reconstruction accuracies obtained with connectome eigenmodes to the density threshold further underscores the complexity of this approach in comparison to geometric eigenmodes, which require no choices regarding a specific threshold.

Our focus on high-resolution, vertex-level connectomes is intended to allow fair comparison with previous work examining the efficacy of connectome eigenmodes in reconstructing functional data [51,52]. However, many other studies on connectomes first apply a discrete parcellation to the data and investigate the properties of connectomes generally mapped at the level of ~$10^2$ to ~$10^3$ parcellated brain regions [57,58]. For completeness, we also derived eigenmodes using connectomes parcellated at different resolutions, generated using the Schaefer400, Schaefer600, Scahefer800, and Schaefer1000 parcellations. These parcellations were chosen because they have at least 200 parcels per hemisphere, allowing us to calculate at least 200 connectome eigenmodes for direct comparison with the 200 geometric eigenmodes used in the study. Note, however, that this approach ignores the spatial embedding of the brain, and resulting in inferior reconstruction accuracy when compared to geometric eigenmodes (Fig. S11). This result again confirms the importance of capturing local connectivity and regional geometry when deriving an appropriate anatomical basis set for brain function.

S7. Derivation of EDR eigenmodes

EDR eigenmodes were derived also by solving Eq. (S6). However, in this case the unnormalized graph Laplacian, $L$, was defined using a synthetically constructed 32,492 × 32,492 adjacency matrix, $A_E$, that follows a stochastic EDR. To construct $A_E$, we used the group-averaged, unthresholded weighted connectome, $W_{\text{connectome}}$, from Section S6. We then fitted the variation of weights as a function of the Euclidean distance, $d$, between vertices in the cortical surface by an exponential function of the form $e^{-\alpha \cdot d}$, where $\alpha$ is a scale exponent parameter. The fitting was performed in MATLAB 2019b using a nonlinear least-squares method, resulting in an optimal empirical parameter value of $\alpha_{\text{empirical}} = 0.12$, consistent with prior estimates based on the connection probability versus distance function [59]. We then generated a random, binary adjacency matrix following a stochastic EDR wiring process, where two vertices were connected with probability $e^{-\alpha \cdot d}$ with $\alpha = \alpha_{\text{empirical}} = 0.12$.

The adjacency matrix $A_E$ generated by an instance of the EDR model had a connection density of 1.55%, which was substantially higher than the 0.10% density of $A_C$. We therefore constructed another version of connectome eigenmodes that thresholded the group-averaged empirical connectome to achieve a final $A_C$ with a density that matches $A_E$ to allow fair comparison (we termed this the density-matched connectome eigenmodes). As outlined above, Fig. S9 presents a more thorough evaluation of the performance of connectome eigenmodes as a function of network connection density.

S8. Comparison between geometric eigenmodes and a functionally derived basis set

To further evaluate the efficacy of geometric eigenmodes in representing brain activity, we compared their performance to a basis set derived via a principal component analysis (PCA) of functional data itself. Since PCA defines a linearly optimal decomposition of the functional data, we sought to evaluate the out-of-sample generalizability of this approach by applying the PCA to a training set of 200 individuals and validating its reconstruction accuracy on a held-out test set of 55 individuals. The number of individuals for the training set was chosen to produce 200 principal components (PCs), matching the number of geometric eigenmodes analyzed in the study. Note that the PCs were arranged in order of the variance in the data that they explain.

For the task-evoked data, we constructed 7 training datasets, one for each of the 7 key task contrasts in Table S2. Each training dataset was constructed by forming a matrix of size 32,492 vertices × 200 individuals with each task's activation data. We then applied a spatial PCA to obtain 200 spatial PCs, describing common modes of variance in spatial activation patterns across individuals (Fig. S12).

For the task-free resting-state data, a similar approach can be implemented, where a training dataset is constructed by temporally concatenating the data of individuals in the training set, producing a matrix of size 32,492 × 240,000 (i.e., 240,000 = 1200 time frames × 200 individuals). However, performing a standard PCA on this very large data matrix carries a high computational burden. We therefore used the MELODIC Incremental Group-PCA (MIGP) method, which has been shown to produce outputs that closely approximate those of a standard PCA in a fully-concatenated dataset using a much more computationally efficient method [60]. In summary, MIGP temporally concatenates the data of a small number of individuals (here we used two individuals) to construct the matrix $W$. It then uses $W$ to construct a time frame × time frame covariance matrix and applies an eigenvalue decomposition to extract the top $m$ temporal eigenvectors (here we used $m$ = 1200), and multiplies the eigenvectors with $W$ to obtain $m$ weighted spatial eigenvectors, yielding a new $W$. Then, $W$ is incrementally updated by concatenating it with the next individual's data and the entire process above is repeated to obtain the new $W$ representing $m$ spatial eigenvectors. This iterative method was performed until all the individuals in the training set were considered. We retained the top 200 spatial PCs from $W$.

The PCs obtained from the training sets in the task-evoked and task-free resting-state data were then used to decompose the data of the individuals in the test set and calculate the corresponding out-of-sample reconstruction accuracy, following Section S3. Note that the performance of the geometric eigenmodes is inherently out-of-sample, since the modes were derived from a cortical surface mesh representation that is independent of the fMRI datasets considered here.

We emphasize that components derived from PCA only capture statistical properties of the data (i.e., maximal variance) and do not provide any mechanistic insights; i.e., they are statistical or phenomenological descriptions of the data and are ordered according to variance explained without regard for spatial wavelength. They are thus fundamentally different from geometric eigenmodes, which are directly linked to the structural properties of the system, are informed by a generative model of how brain structure gives rise to function [23], and are arranged according to their spatial frequencies or wavelengths. As shown in Fig. S13, this distinction leads to an early performance advantage for PCA over geometric eigenmodes, in which the first few PCs can more accurately capture the data than the first few geometric eigenmodes. This is because the PCs do not have the same ordering constraint as the geometric eigenmodes; i.e., early PCs are free to span any spatial wavelength, as dictated by the optimal decomposition of the data, whereas geometric eigenmodes are ordered from long to short wavelengths, by construction. However, Fig. S13 also shows that there is a limit on the accuracy with which the PCs can reconstruct out-of-sample data, likely because higher-order PCs capture idiosyncratic properties of the training data. In contrast, adding more geometric eigenmodes improves the model's capacity to resolve reproducible shorter-wavelength features of both the training and test data, such that the out-of-sample reconstruction surpasses the accuracy of the PCs across most comparisons by about 40–50 modes and beyond.

S9. Comparison between geometric eigenmodes and Fourier basis sets

To further confirm that the performance of geometric eigenmodes is not trivially driven by how the mathematics of any basis set expansion, we compared the geometric eigenmodes to six simple spatial Fourier basis sets based on combinations of sines and/or cosines. The Fourier basis sets were constructed using the following real-valued functions:

$$\mathrm{F}_1 := \cos\left(\frac{2\pi(j_x-1)x}{L_x} + \frac{2\pi(j_y-1)y}{L_y} + \frac{2\pi(j_z-1)z}{L_z}\right), \qquad (S8)$$

$$\mathrm{F}_2 := \mathrm{C}_1\cos\left(\frac{2\pi(j_x-1)x}{L_x}\right) + \mathrm{C}_2\cos\left(\frac{2\pi(j_y-1)y}{L_y}\right) + \mathrm{C}_3\cos\left(\frac{2\pi(j_z-1)z}{L_z}\right), \qquad (S9)$$

$$F_3 \coloneqq C_1 \cos\left(\frac{2\pi(j_x-1)x}{L_x}+\frac{2\pi(j_y-1)y}{L_y}+\frac{2\pi(j_z-1)z}{L_z}\right)+ C_2 \sin\left(\frac{2\pi(j_x-1)x}{L_x}+\frac{2\pi(j_y-1)y}{L_y}+\frac{2\pi(j_z-1)z}{L_z}\right), \quad (S10)$$

where $j_x$, $j_y$, $j_z$ are integer constants, $x$, $y$, $z$ are the spatial positions of each point on the spherical surface mesh representation of the cortex, $L_x$, $L_y$, $L_z$ are the periods in each direction, and $C_1$, $C_2$, $C_3$ are fitting constants. For each function, we constructed a regular and an irregular version, resulting in six basis sets. The regular version corresponds to when $j_x = j_y = j_z = j$ such that the spatial wavelengths of mode $j$ in the x-, y-, and z-directions are regularly spaced and increase by $\frac{2\pi}{L_x}, \frac{2\pi}{L_y}, \frac{2\pi}{L_z}$ as the mode number increases, which is a standard implementation in Fourier analysis. The irregular version corresponds to when $j_x$, $j_y$, $j_z$ are integer combinations and not necessarily equal; hence, the spatial wavelengths in the x-, y-, and z-directions are irregularly spaced. We implemented this version because it affords the Fourier basis sets greater freedom and thus the best possible chance of performing well. To associate a $(j_x, j_y, j_z)$ combination to a single mode $j$, we arranged the combinations in order of increasing $j_x + j_y + j_z$. For example, the $(j_x, j_y, j_z)$ combination for the first ten modes follows the set $\{(1,1,1), (1,1,2), (1,2,1), (2,1,1), (1,2,2), (2,1,2), (2,2,1), (1,1,3), (1,3,1), (3,1,1)\}$. These two versions allow us to explore how the spatial wavelengths can affect the decompositions of the Fourier basis sets, given that the spatial wavelengths of the geometric eigenmodes are irregularly spaced (see Table S1).

Next, we defined the periods to be $L_i \coloneqq [\max(i) - \min(i)]$ for $i = x, y, z$, to have a heuristic that respects the shape of the brain. In addition, the $-1$ in $(j_i - 1)$ ensures that the first mode is constant to make it comparable to the first geometric mode. The forms in Eqs. (S9) and (S10) have more degrees of freedom, with the fitting constants, $C_1$, $C_2$, $C_3$, estimated separately, allowing us to weight the cosine and/or sine functions differently. Therefore, mode *j* now has multiple amplitudes to be estimated during the decomposition in Eq. (S4) instead of just one. These added degrees of freedom result in an increased model complexity relative to geometric eigenmodes; more specifically, the Fourier basis sets formed by Eqs. (S9) and (S10) involve estimating 2 and 3 coefficients per mode, respectively, whereas geometric eigenmodes require fitting just one coefficient per mode.

Although other forms and complicated choices can be made to construct a Fourier basis set, the above choices are well motivated by their simplicity, yielding unique, real-valued spatial modes, and cover the key implementation choices one could make. Figure S14A shows the spatial profiles of the resulting modes with unit coefficients, which we used to reconstruct task-activation maps and resting-state data (similar to Fig. 1D). Figure S14B shows that geometric eigenmodes significantly outperform the Fourier basis sets in reconstructing both task-evoked and resting-state data, further emphasizing that the accurate representation provided by geometric eigenmodes is not a trivial result of any basis set expansion. Moreover, this conclusion remains regardless of how the spatial wavelengths of the modes in the x-, y-, and z-directions are defined. To reiterate, even though the Fourier basis sets based on Eqs. (S9) and (S10) have more degrees of freedom to fit the data when compared to geometric eigenmodes, geometric eigenmodes still show superior performance, underscoring their parsimony in accounting for brain dynamics.

We present the above analysis of Fourier basis sets for completeness and to demonstrate that such functions cannot accommodate the boundary conditions of non-regularly shaped objects, such as the cortical surface (e.g., Dirichlet or Neumann conditions for the medial wall) [61]. This is because Fourier basis sets can only be properly constructed for objects with regular shapes (e.g., rectangle) and are more suitable to be used for analyzing functions defined on such shapes (e.g., 2D images). Hence, for the cortical surface defined on a Riemannian manifold, using a Fourier basis set is generally not well motivated and using the eigenmodes of the Laplace-Beltrami operator is more appropriate [62,63]. In fact, mathematically speaking, LBO eigenmode analysis on the Riemannian manifold is considered a generalization of Fourier analysis [64]. The same applies to discrete networks, such as a connectome; for this reason, the eigendecomposition of the graph Laplacian (as done in the work) is commonly used in the field of spectral graph theory as an alternative to a classical Fourier transform [65]. Thus, whilst it is possible to decompose spatial maps of brain activity using Fourier basis sets, they are highly inefficient and poorly suited to the current problem. We therefore do not advocate their use. We also note that Fourier basis sets offer no insights into the generative processes underlying brain activity. Our primary focus is to compare physiologically principled and anatomically constrained basis sets (i.e., geometric and connectome eigenmodes) to uncover the critical constraints on brain dynamics, and not to identify statistically optimal basis sets.

S10. Modal power spectra of task-evoked activation maps

To investigate the spectral content of task-evoked activation maps, we calculated their modal power spectra by using the mode decomposition in Eq. (S4) and taking the absolute square of the amplitudes, $a$. This is analogous to calculating the temporal power spectral density from Fourier analysis. We then normalized the power in mode $j$ with respect to the total power in all modes, such that

$$P_j = \frac{|a_j|^2}{\sum_{j=1}^{N} |a_j|^2}. \qquad (S11)$$

We calculated the modal power spectra of two sets of task-evoked data. The first set comprises unthresholded activation maps from the HCP task-evoked data described in Section S4.2. Before doing the spectral analysis, we took the group average of the activation maps across 255 individuals. Then, we analyzed the power spectrum of the group-averaged activation map of each task contrast. The results in Figs. 3A and S13A show the mean power spectrum across the 47 HCP task-contrast maps. Contrast-specific power spectra for the 7 key HCP task contrasts are shown in Fig. S16.

The second set comprises 10,000 activation maps from 1,178 independent experiments from the NeuroVault repository (https://neurovault.org/) [66]. We used the python module Nilearn [67] to retrieve activation maps from NeuroVault that were unthresholded and with a modality tag of fMRI-BOLD. We also projected the activation maps from volume onto the fsLR-32k CIFTI space to match the HCP data using Nilearn (via the function nilearn.surface.vol_to_surf). We then analyzed the power spectrum of each activation map. The results in Figs. 3A and S17A show the mean power spectrum across the 10,000 NeuroVault maps.

Typical image processing pipelines induce some degree of spatial smoothing, which serves to filter out high-frequency (short-wavelength) spatial patterns of activity. To ensure that the dominant long-wavelength power we identified in our analyses was not merely the product of spatial smoothing induced by fMRI preprocessing, we analyzed the power spectra of surrogate random data with varying levels of smoothing. We first generated 10,000 random maps in volume space taken from a zero-mean Gaussian distribution. We then smoothed the surrogate maps at kernel sizes with full-width at half-maximum (FWHM) ranging from 0 to 50 mm. Finally, we projected the surrogate maps onto the fsLR-32k CIFTI space (via Nilearn as discussed above) and analyzed their modal power spectra. Note that we performed the smoothing in volumetric instead of surface space to mimic prevailing neuroimaging processing practices, in which data processing is commonly applied to volumetric data before results are resampled to the cortical surface [68]. This procedure thus also incorporates any smoothing induced by the resampling procedure.

We compared the modal power spectra of the empirical activation maps and the surrogate maps by calculating the mean square logarithmic error (MSLE),

$$\mathrm{MSLE} = \sqrt{\frac{1}{N}\sum_{j=1}^{N=200}\left[\log_{10}\left(P_j^{\mathrm{empirical}}\right) - \log_{10}\left(P_j^{\mathrm{surrogate}}\right)\right]^2}, \qquad (S12)$$

where $P_j^{\mathrm{empirical}}$ and $P_j^{\mathrm{surrogate}}$ is the power in mode $j$ in the empirical and surrogate maps, respectively. The MSLE is more effective as a distance measure than the typical mean square error because the power of long- to short-wavelength modes have scales that vary across several orders of magnitude.

The colored lines in Fig. S17A show the modal power spectra of the surrogate data, Fig. S17B shows the average distance (i.e., MSLE) between empirical and surrogate spectra averaged across contrasts, and Fig. S17C shows MSLE separately obtained for each task contrast (instead of averaged across contrasts). We found that smoothing kernels smaller than 10 mm do not adequately capture the concentration of spectral power in long spatial wavelengths of empirical maps (Figs. 3A and S17A). In fact, the surrogate data needed an average kernel size of 20 mm (best fits obtained from the minimums in Fig. S17B) to achieve a spectrum comparable to the empirical data, which is considerably larger than the size of the kernels used in neuroimaging processing (~8 mm in volume space [68] and ~15 mm in surface space [69]). Similar results were obtained when task contrasts were separately analyzed instead of their average, with minimum MSLE showing minor variations around the mean (solid lines in Fig. S17C). These findings thus confirm that the long-wavelength content of empirical activation maps cannot be explained by preprocessing alone. Moreover, whilst spatially extended patterns of task activations can also potentially be recovered using simple smoothing functions and avoiding statistical thresholding, the geometric eigenmode approach provides much deeper understanding of the mechanisms underlying the activations because the modes can be directly linked to rigorous biophysical models of brain dynamics as established by NFT (see for example [6,7,14,23]). They also offer a natural multiscale characterization of the data.

S11. Mode removal to isolate contributions from long- and short-wavelength modes

To understand the contributions of long- and short-wavelength geometric eigenmodes in the reconstruction of task-evoked activation maps, we sequentially removed modes prior to performing the reconstruction process described in Section S3. Specifically, we started by reconstructing the 7 key HCP contrast maps using 200 geometric eigenmodes and calculated the reconstruction accuracy (i.e., correlation between empirical and reconstructed maps), serving as our baseline. We then performed an incremental, sequential removal of modes starting from long-wavelength modes (i.e., removing mode 1, modes 1–2, modes 1–3, modes 1–4, …, modes 1–200) and calculated the reconstruction accuracy at every increment. We repeated the same procedure but starting from short-wavelength modes (i.e., removing mode 200, modes 199–200, modes 198–200, modes 197–200, …, modes 1–200).

S12. Models of brain dynamics

*S12.1 NFT wave model*

As stated in the main text, an implication of the superior performance of geometric eigenmodes is that neuronal activity is dominated by wave dynamics, as predicted by NFT. To investigate whether wave dynamics can explain complex spatiotemporal patterns of neuronal activity, we implemented a simple NFT wave model in which dynamics are described by an isotropic damped wave equation without regeneration [5,6,12],

$$\left[\frac{1}{\gamma_s^2}\frac{\partial^2}{\partial t^2} + \frac{2}{\gamma_s}\frac{\partial}{\partial t} + 1 - r_s^2\nabla^2\right]\phi(\boldsymbol{r},t) = Q(\boldsymbol{r},t), \qquad (S13)$$

where $\phi(\boldsymbol{r},t)$ is the neural activity at location $\boldsymbol{r}$ and time $t$, $Q$ is an external input, $\gamma_s$ is the damping rate, and $r_s$ is the spatial length scale of the wave propagation (conceptually related to the $\alpha$ length scale of the stochastic EDR wiring process in Section S7). This form tells us that an impulse input will produce an activity that dissipates at a rate of $\gamma_s$ and propagating at a velocity of $\gamma_s r_s$. Here, we treated $\gamma_s$ as a fixed parameter with value of 116 $s^{-1}$ taken from electrophysiological estimates [7] and $r_s$ as a free parameter. We applied the model on the cortical surface mesh with 32,492 vertices per hemisphere described in Section S2 to solve the activity at each vertex. Note that the propagation of activity between points is governed by their white-matter connectivity, with strength that decays approximately exponentially with distance. This distance-dependence is more apparent when Eq. (S13) is converted into its equivalent integral form as follows. Consider two points on the neocortex at locations $\boldsymbol{r}'$ and $\boldsymbol{r}$ connected by a white-matter tract (see visual schematic in Fig. S18). The activity $\phi(\boldsymbol{r},t)$ at location $\boldsymbol{r}$ and time $t$ can be considered as a spatiotemporal convolution of the source $Q(\boldsymbol{r}',t')$ at location $\boldsymbol{r}'$ and time $t'$ and the white-matter-based connectivity kernel $W(\boldsymbol{r},t;\boldsymbol{r}',t')$; i.e.,

$$\phi(\boldsymbol{r},t) = \int W(\boldsymbol{r},t;\boldsymbol{r}',t')Q(\boldsymbol{r}',t')d^2\boldsymbol{r}'dt'\,. \qquad (S14)$$

In the isotropic case, $W$ depends only on the spatial separation between points and the time difference such that $W(\boldsymbol{r},t;\boldsymbol{r}',t') \coloneqq W(\boldsymbol{r}-\boldsymbol{r}',t-t')$. The kernel $W$ is also known as the Green's function, which is the activity $\phi$ due to a point source. Using the Green's function method

[70], previous work has shown that an appropriate expression for $W$, such that it becomes a solution of the damped wave equation in Eq. (S13) with an intense point source $Q(\boldsymbol{r},t;\boldsymbol{r}',t') = \delta(\boldsymbol{r}-\boldsymbol{r}')\delta(t-t')$ and for $|\boldsymbol{r}-\boldsymbol{r}'| \leq \gamma_s r_s(t-t')$, is [4,6,71,72]

$$W(\boldsymbol{r}-\boldsymbol{r}',t-t') = \frac{\gamma_s}{r_s}\frac{\exp\left(\frac{-|\boldsymbol{r}-\boldsymbol{r}'|}{r_s}\right)}{\sqrt{\gamma_s^2 r_s^2 (t-t')^2 - |\boldsymbol{r}-\boldsymbol{r}'|^2}}\Theta[\gamma_s r_s(t-t') - |\boldsymbol{r}-\boldsymbol{r}'|], \qquad (S15)$$

where $|\boldsymbol{r}-\boldsymbol{r}'|$ is the white-matter tract distance between points, $\Theta$ is the Heaviside step function, and $\gamma_s$ and $r_s$ are the damping rate and spatial length scale, respectively, as defined in Eq. (S13). See [6] for a detailed derivation of Eq. (S15). Equation (S15) thus demonstrates how the wave dynamics defined by Eq. (S13) can be directly related to an underlying isotropic anatomical connectivity that decays exponentially with distance. In particular, solving the damped wave equation in differential form in Eq. (S13) is equivalent to solving the integral equation,

$$\phi(\boldsymbol{r},t) = \frac{\gamma_s}{r_s}\int \frac{\exp\left(\frac{-|\boldsymbol{r}-\boldsymbol{r}'|}{r_s}\right)}{\sqrt{\gamma_s^2 r_s^2 (t-t')^2 - |\boldsymbol{r}-\boldsymbol{r}'|^2}}\Theta[\gamma_s r_s(t-t') - |\boldsymbol{r}-\boldsymbol{r}'|]Q(\boldsymbol{r}',t')d^2\boldsymbol{r}'dt', \qquad (S16)$$

with the latter explicitly showing the spatiotemporal effect invoked by white-matter connectivity of a characteristic range of $r_s$ (~84 mm [6]). There are other more complex variations of the wave equation in Eq. (S13), but this simple version of the model evaluates the basic physical process that could account for empirical data. Incorporation of spatial heterogeneities or structured input into the wave equation in Eq. (S13) is a topic for future investigation.

Despite the simplicity of the model, the full spatiotemporal solution of Eq. (S13) at 32,492 vertices is computationally expensive. We therefore used the following approach to efficiently solve Eq. (S13) on the cortex. First, we assumed that $\phi$ is time-space separable because the structure of the system is constant in time, such that

$$\phi(\boldsymbol{r},t) = \phi(t)\phi(\boldsymbol{r}). \qquad (S17)$$

By substituting Eq. (S17) into Eq. (S13), the term $\nabla^2\phi(\boldsymbol{r},t)$ becomes $\phi(t)\nabla^2\phi(\boldsymbol{r})$. Note that $\nabla^2\phi(\boldsymbol{r})$ is the same as the left-hand side of the Helmholtz equation in Eq. (S1); hence, the spatial component of the traveling wave solution, $\phi(\boldsymbol{r})$, on the cortical surface can be written as a combination of geometric eigenmodes $\psi_j(\boldsymbol{r})$. Therefore, Eq. (S17) can be written as

$$\phi(\boldsymbol{r},t) = \sum_{j=1}^{N}\phi_j(t)\psi_j(\boldsymbol{r}), \qquad (S18)$$

where $\psi_j(\boldsymbol{r})$ is mode $j$, $\phi_j(t)$ is the time-varying component of mode $j$, and $N$ is the number of modes. Second, we employed mode decomposition on the input $Q(\boldsymbol{r},t)$ such that

$$Q(\boldsymbol{r},t) = \sum_{j=1}^{N} q_j(t)\psi_j(\boldsymbol{r}), \tag{S19}$$

where $q_j(t)$ is the time-varying amplitude obtained as in Section S3. Important consequences of the above separation into temporal and spatial factors are that the spatial modes are not sensitive to the local temporal dynamics, nor to the value of $r_s$ so long as it is small compared to the radius of curvature of the surface.

Substituting Eqs. (S18), (S19), and (S1) into Eq. (S13), we obtain

$$\sum_{j=1}^{N}\left[\frac{1}{\gamma_s^2}\frac{\partial^2}{\partial t^2} + \frac{2}{\gamma_s}\frac{\partial}{\partial t} + 1 + r_s^2\lambda_j\right]\phi_j(t)\psi_j(\boldsymbol{r}) = \sum_{j=1}^{N} q_j(t)\psi_j(\boldsymbol{r}), \tag{S20}$$

where $\lambda_j$ is the eigenvalue corresponding to mode $j$. Therefore, the ordinary differential equation for solving the time-varying component of mode $j$, $\phi_j(t)$, is

$$\left[\frac{1}{\gamma_s^2}\frac{d^2}{dt^2} + \frac{2}{\gamma_s}\frac{d}{dt} + 1 + r_s^2\lambda_j\right]\phi_j(t) = q_j(t). \tag{S21}$$

One can solve Eq. (S21) via numerical methods, but it can be solved more simply – and exactly – by taking its Fourier transform vs. time, thereby yielding the algebraic equation,

$$\left[-\frac{\omega^2}{\gamma_s^2} - \frac{2i\omega}{\gamma_s} + 1 + r_s^2\lambda_j\right]\phi_j(\omega) = q_j(\omega), \tag{S22}$$

where $\omega$ is the temporal angular frequency, $\phi_j(\omega)$ is the Fourier transform of $\phi_j(t)$, and $q_j(\omega)$ is the Fourier transform of $q_j(t)$. Therefore, the solution $\phi_j(t)$ has the form,

$$\phi_j(t) = \mathcal{F}^{-1}\{\phi_j(\omega)\} = \mathcal{F}^{-1}\left\{\frac{\gamma_s^2 q_j(\omega)}{-\omega^2 - 2i\omega\gamma_s + \gamma_s^2\left(1 + r_s^2\lambda_j\right)}\right\}, \tag{S23}$$

where $\mathcal{F}^{-1}$ is the inverse Fourier transform operator. Finally, Eq. (S23) can be substituted into Eq. (S18) to obtain the neural activity $\phi(\boldsymbol{r},t)$ at every vertex.

As noted above, the spatial part of Eq. (S17) satisfies the Helmholtz equation in Eq. (S1). This differential equation is equivalent to an integral form (Eq. S16), where the activity at a given point gives rise to activity elsewhere with a weight that represents the connectivity that is implicit in NFT. In the cortical case, this corresponds to an exponential decrease with distance—i.e., an EDR-like spatial dependence—consistent with experimental evidence [6]. Hence, the use of geometric eigenmodes implicitly incorporates an EDR-like connectivity, which also includes long-range connections, but does not directly account for topologically complex connections not conforming to a simple exponential rule.

*S12.2 Neural mass model*

We compared the performance of our simple NFT wave model to a biophysical large-scale neural mass model, where mesoscopic dynamics emerge from the interactions of neural populations (i.e., neural masses) coupled via an empirical anatomical connectivity [73]. In a typical neural mass model, each brain region $i$ has its own mean-field population dynamics and its temporal activity, $S_i$, is defined by the general equation,

$$\frac{dS_i}{dt} = f(\boldsymbol{S}, \theta_i, C, G), \tag{S24}$$

where $f$ is a function describing the evolution of the region's activity. The function depends on the activity of other regions $\boldsymbol{S}$, local population parameters $\theta_i$, anatomical connectivity between regions $C$ (a parcellated version of the vertex-resolution $W_{\text{connectome}}$ described in Section S6), and global coupling parameter $G$ that scales the connectivity between regions.

There are several whole-brain neural mass models available in the literature that we can use [74]; from simple phase oscillator models (e.g., Kuramoto model [75]) to more complex biophysical population models (e.g., Wilson-Cowan model [76]). All these models follow the form of Eq. (S24), especially their reliance on an anatomical interregional connectivity matrix, $C$. Here, we focus on one widely used neural mass model, i.e., the balanced excitation-inhibition (BEI) model [77–80]. The BEI model uses a mean-field approach to approximate local population dynamics in each brain region, which are coupled via an anatomical connectivity matrix derived from dMRI. Each brain region $i$ comprises interacting populations of excitatory ($E$) and inhibitory ($I$) neurons governed by the following nonlinear stochastic differential equations,

$$\frac{dS_i^{(E)}}{dt} = -\frac{S_i^{(E)}}{\tau_E} + \left(1 - S_i^{(E)}\right)\gamma r_i^{(E)} + \sigma \nu_i(t), \tag{S25}$$

$$\frac{dS_i^{(I)}}{dt} = -\frac{S_i^{(I)}}{\tau_I} + r_i^{(I)} + \sigma \nu_i(t), \tag{S26}$$

$$r_i^{(E)} = H^{(E)}\left(I_i^{(E)}\right) = \frac{a_E I_i^{(E)} - b_E}{1 - \exp\left[-d_E\left(a_E I_i^{(E)} - b_E\right)\right]}, \tag{S27}$$

$$r_i^{(I)} = H^{(I)}\left(I_i^{(I)}\right) = \frac{a_I I_i^{(I)} - b_I}{1 - \exp\left[-d_I\left(a_I I_i^{(I)} - b_I\right)\right]}, \tag{S28}$$

$$I_i^{(E)}(t) = I^{\text{ext}} + W_E I_0 + w_{EE} S_i^E(t) + GJ \sum_j C_{ij} S_j^{(E)}(t) - w_{IE} S_i^{(I)}(t), \tag{S29}$$

$$I_i^{(I)}(t) = W_I I_0 + w_{EI} S_i^{(E)}(t) - S_i^{(I)}(t), \tag{S30}$$

where $S_i^{(E,I)}$, $r_i^{(E,I)}$, and $I_i^{(E,I)}$ represent the synaptic gating variable, firing rate, and total input current, respectively, for $E$ and $I$ populations. The parameters $\tau_{E,I}$ are the time constants, $\gamma$ is a kinetic rate constant, and $\nu_i(t)$ is a time-varying random Gaussian input with standard deviation $\sigma$. The functions $H^{(E,I)}$ are sigmoidal neuronal response functions transforming the total input currents $I_i^{(E,I)}$ into firing rates $r_i^{(E,I)}$, which are parametrized by the gain factors $a_{E,I}$, threshold currents $b_{E,I}$, and curvature parameters $d_{E,I}$. In Eqs. (S29) and (S30), $I_i^{\text{ext}}$ is the external input, $I_0$ is the local input current scaled by $W_E$ and $W_I$ for the excitatory and inhibitory populations, respectively, $w_{EE}$ is the excitatory-excitatory strength, $w_{EI}$ is the excitatory-inhibitory strength, $w_{IE}$ is the inhibitory-excitatory strength, $G$ is the global coupling parameter, $J$ is the effective NMDA conductance, and $C_{ij}$ is the structural connectivity strength between region $i$ and region $j$ estimated from dMRI.

Overall, the BEI model has 15 fixed parameters and 4 free parameters, as detailed in Table S3. The values of the fixed parameters were taken from [78]. To allow direct comparison, we also used the discretized connectome data provided by [78] to define the inter-regional structural connectivity matrix, $C$. These connectome data were derived from minimally preprocessed dMRI data of 334 unrelated HCP subjects and constructed via FSL's probabilistic tractography [81]. We refer the readers to the article for further details of data processing and connectome construction [78]. Our results did not change when the parcellated version of the connectome data described in Sections S4.4 and S6 was used. We note that numerical solutions of NFT equations (such as in Eq. S10) also spatially discretize the cortex, but into a very fine array of points, before integrating whatever temporal dynamics have been chosen at each point with connectivity that is correct in the continuum limit. So neural mass models are approximations to the more general NFT approach [4,82]. We also reiterate that the local dynamics do not affect the spatial eigenfunctions of the NFT wave model.

**Table S3. Fixed and free parameters of the neural mass model.**

| **Fixed parameters** | | **Free parameters** |
|---|---|---|
| **Symbol** | **Value** | **Symbol** |
| $\tau_E$ | 0.1 s | $w_{EE}$ |
| $\tau_I$ | 0.01 s | $w_{IE}$ |
| $\gamma$ | 0.641 | $w_{EI}$ |
| $\sigma$ | 0.01 nA | $G$ |
| $a_E$ | 310 nC$^{-1}$ | |
| $b_E$ | 125 s$^{-1}$ | |
| $d_E$ | 0.16 s | |
| $a_I$ | 615 nC$^{-1}$ | |
| $b_I$ | 177 s$^{-1}$ | |
| $d_I$ | 0.087 s | |
| $I^{ext}$ | 0 nA | |
| $I_0$ | 0.382 nA | |
| $W_E$ | 1.0 | |
| $W_I$ | 0.7 | |
| $J$ | 0.15 nA | |

*S12.3 Hemodynamic model*

To simulate fMRI data, we transformed the neural activity generated by the NFT wave and BEI models to a blood oxygen-level dependent (BOLD) signal using the well-established Balloon-Windkessel hemodynamic model [83]. Note that this model is a simple approximation to more detailed models of the physiological hemodynamic processes underlying the BOLD signal [84–86], but we use this approximation here to allow direct comparison with the vast majority of modeling studies in the literature [78–80,87,88]. The BOLD-fMRI signal in each vertex or brain region $i$ is governed by the following differential equations,

$$\frac{dz_i}{dt} = N_i(t) - \kappa z_i(t) - \gamma[f_i(t) - 1], \tag{S31}$$

$$\frac{df_i}{dt} = z_i(t), \tag{S32}$$

$$\frac{dv_i}{dt} = \frac{1}{\tau}\left[f_i(t) - v_i^{1/\alpha}(t)\right], \tag{S33}$$

$$\frac{dq_i}{dt} = \frac{1}{\tau}\left\{\frac{f_i(t)}{\rho}\left[1 - (1-\rho)^{1/f_i(t)}\right] - v_i^{1/\alpha - 1}(t)\right\}, \tag{S34}$$

$$\frac{dy_i}{dt} = V_0\left\{k_1[1 - q_i(t)] + k_2\left[1 - \frac{q_i(t)}{v_i(t)}\right] + k_3[1 - v_i(t)]\right\}, \tag{S35}$$

where $z_i$, $f_i$, $v_i$, $q_i$, and $y_i$ are the vasodilatory signal, blood inflow, blood volume, deoxyhemoglobin content, and BOLD signal variables, respectively. The variable $N$ represents the neural activity generated by the wave and BEI models. For the wave model, $N(t)$ is $\phi(t)$, while for the BEI model, $N_i(t)$ is $S_i^{(E)}(t)$. The model parameters and their values taken from previous works [83,89] were as follows: $\kappa = 0.65$ s$^{-1}$ is the signal decay rate, $\gamma = 0.41$ s$^{-1}$ is the flow-dependent elimination rate, $\tau = 0.98$ s is the hemodynamic transit time, $\alpha = 0.32$ is the Grubb's exponent, $\rho = 0.34$ is the resting oxygen extraction fraction, $V_0 = 0.02$ is the resting blood volume fraction, and $k_1 = 3.72$, $k_2 = 0.53$, and $k_3 = 0.53$ are 3T fMRI parameters.

*S12.4 Modeling resting-state dynamics*

We used the NFT wave and BEI models, together with the hemodynamic model, to estimate various spontaneous FC properties. For the wave model, we solved Eq. (S23) with a white noise input to mimic the absence of any structured stimulus, following previous studies [7,8]. We combined the resulting solution with the hemodynamic model in Eqs. (S31) to (S35) to simulate the BOLD-fMRI signal. The simulated BOLD signal was downsampled to a sampling interval of 0.72 s with 1200 time frames to match the resting-state HCP data described in Section S4.3. Finally, we parcellated the simulated BOLD signal using the HCP-MMP1 parcellation (Section S5) and calculated the correlation matrix representing the model FC.

For the BEI model, we can use the same method described above by calculating the model FC starting from the numerical solutions of Eq. (S25) for each brain region. However, [78] has shown that one can linearize the equations of the BEI and hemodynamic models to obtain an analytic approximation of the model FC. Due to the large number of model parameters in the BEI model, we used this analytic approximation in this study as it allows more comprehensive and computationally efficient model fitting. See [78] for further details.

We fitted the model FCs to empirical FCs (also parcellated using the HCP-MMP1 parcellation) of the same HCP participants described in Section S4.1 by fitting each model's free parameters. We divided the participant sample into a training set with 125 individuals and test set with 125 individuals to enable out-of-sample evaluation of model performance and avoid overfitting. In particular, we fitted the model parameters on the training set's data and used the fitted model parameters to predict the test set's data. Model fitting and performance evaluation were based on three widely used FC-related metrics [79,80]: static pairwise FC (edge FC), node-level average FC (node FC), and dynamic properties of FC (FCD).

The edge FC metric was calculated by taking the Pearson correlation of the upper triangular elements (i.e., the strength of FC edges) of the $z$-transformed model and empirical FCs. We only took the upper triangular elements because the FC values are symmetric with respect to the diagonal. Higher correlations represent a better fit between model and data.

The node FC metric was calculated by taking the Pearson correlation of the average FC strength of each brain region in the model and empirical FCs. The average FC strength of a brain region $i$ was defined as $\sum_{j=1}^{N} FC_{ij}$. Once again, higher correlations indicate better model fits.

The FCD metric captures the spatiotemporal statistics of resting-state activity and was calculated as follows [80]. The time series at each region $i$ was filtered between 0.04 and 0.07 Hz using a 2nd-order Butterworth filter; this band was based on [90] and was motivated by its functional relevance to the brain [91,92]. Then, the time series was Hilbert transformed to calculate the quantity $y_i(t) = x_i(t) + jH_i(t)$, where $j$ is the imaginary number. The instantaneous complex argument $\theta_i(t) = \tan^{-1}[H_i(t)/x_i(t)]$ was then computed. The level of synchrony between regions $i$ and $j$ at time $t$, $\Delta(i,j,t)$, was calculated as,

$$\Delta(i,j,t) = \cos\left[\theta_i(t) - \theta_j(t)\right]. \quad (S36)$$

Note that we do not term $\theta_i(t)$ a phase, as is sometimes done in the literature, because our signals are broadband, whereas this interpretation is only applicable to signals that are nearly monochromatic. We compute this quantity for comparison with prior work [79,80]. Then, we calculated the similarity of the global synchrony, $\varphi_{uv}$, between two time instances, $\tau_u$ and $\tau_v$, with $\varphi_{uv}$ defined as

$$\varphi_{uv} = \frac{1}{d_u d_v} \sum_{i>j} \Delta(i,j,\tau_u)\Delta(i,j,\tau_v), \quad (S37)$$

where

$$d_x = \sqrt{\sum_{i>j} [\Delta(i,j,\tau_x)]^2}. \quad (S38)$$

Here, $\varphi_{uv}$ is the FCD, which is a symmetric time × time matrix, interpreted as the similarity of the global synchrony between time $\tau_u$ and $\tau_v$. We then compared the distributions of the upper triangular elements of the model and empirical FCD estimates, concatenated across all individuals or model realizations, as per [80], to better capture the general fluctuations of the data. The FCD distributions of the model and the data were compared using the Kolmogorov-Smirnov (KS) statistic, with lower KS statistic indicating better model fit.

*S12.5 Model optimization*

The wave model has one free parameter—the spatial length scale of wave propagation, $r_s$—that was optimized to fit the empirical fMRI data. Specifically, we constructed a vector of 20 values of $r_s$ evenly spaced between 10 and 100 mm. For each $r_s$ value, the FC metrics described in Section S12.4 were calculated. The optimization landscapes are shown in Fig. S20. We took the value of $r_s$ that minimized the FCD KS statistic because this metric has been found to be the most stringent benchmark for model-data comparisons among the three metrics discussed above [80]. This procedure resulted in an optimized value of $r_s = 28.9$ mm. This estimate is smaller than those obtained previously in analyses of EEG data [7], possibly due to hemodynamic processes limiting neural activity propagation to neighbouring regions or our focus here on cortico-cortical dynamics [84,85].

The BEI neural mass model has four free parameters, i.e., $w_{EE}$, $w_{EI}$, $w_{IE}$, and $G$, that were optimized to fit the data, following [78]. Briefly, feedback inhibition control [77] was implemented to adjust $w_{IE}$ to set the firing rate $r_i^{(E)}$ of all regions to approximately 3 Hz. Hence, the analytic expression for $w_{IE}$ at the steady-state conditions $\langle S^{(E)}\rangle \approx 0.17$ nA and $\langle I^{(E)}\rangle \approx 0.38$ nA is

$$w_{IE} = \frac{W_E I_0 + w_{EE}\langle S^{(E)}\rangle + GJ\langle S^{(E)}\rangle - \langle I^{(E)}\rangle}{\langle S^{(I)}\rangle}, \qquad (S39)$$

where the steady-state value $\langle S^{(I)}\rangle = H^{(I)}\big(\langle I^{(I)}\rangle\big)\tau_I$ was numerically solved using the expression

$$W_I I_0 + w_{EI}\langle S^{(E)}\rangle - H^{(I)}\big(\langle I^{(I)}\rangle\big)\tau_I - \langle I^{(I)}\rangle = 0. \qquad (S40)$$

Hence, the value of $w_{IE}$ dynamically changes as a function of the other free parameters: $w_{EE}$, $w_{EI}$, and $G$. Because finding the optimal parameter set on a multidimensional parameter space is computationally expensive, the analytic FC, as described in Section S12.4, was used. Model fitting was done using the methods of Approximate Bayesian Computation and hierarchical Population Monte Carlo [93,94] that minimizes the distance between empirical and model FCs. See [78] for further details about the model optimization process. This procedure resulted in optimized parameters: $w_{EE} = 9.80$, $w_{EI} = 1.48$, $w_{IE} = 7.13$, and $G = 6.87$.

*S12.6 Measuring time-lagged properties of resting-state dynamics*

In addition to the FC-based metrics used for model fitting and evaluation of model performance (Section S12.4), we investigated whether the wave and mass models can also capture the temporal properties of propagated activity. In particular, we analyzed the lag structure (or lag threads) of resting-state BOLD-fMRI time courses, as proposed by [95,96]. We briefly discuss the algorithm for calculating the lag structure of both empirical and simulated fMRI data below and refer the readers to previous articles [95,96] for further details. Note that there are several other ways of characterizing the spatiotemporal properties of resting-state activity [97–99], but the currently chosen method is sufficient for our purposes.

The algorithm starts by calculating the lagged cross-covariance function of the time series between brain regions. Assuming that BOLD-fMRI time series are aperiodic [100], the time lag (or delay) between regions where the cross-covariance function exhibits an extremum (typically between 0 to 2 s) was obtained [95]. This resulted in an anti-symmetric time-delay matrix $TD$, with elements $\tau_{ij}$ corresponding to the time delay between regions $i$ and $j$, with column $i$ representing the lag map of the system with reference to region $i$, and $\tau_{ij} = -\tau_{ji}$. Therefore, $\tau_{ij} > 0$ means that region $j$ lags behind region $i$. The underlying lag structure was quantified in two ways: (i) taking the mean time lag of each region (mean of each column of $TD$); and (ii) applying a PCA on $TD_z$, which is $TD$ with each column being zero-meaned. The first method obtains the average temporal ordering of brain regions, assuming that a single lag process governs the brain, which has been used in several past studies [95,99,101]. However, for systems like the brain with multiple lag processes, the mean time lag cannot capture all fundamental lag patterns, which can be recovered via PCA [96]. Here, we used the first two dominant PCs, explaining 74% of the variance of the empirical data. Thus, we calculated three lag projections: (i) mean lag; (ii) first PC (PC1 lag); and (iii) second PC (PC2 lag).

The $TD$ matrix was calculated on empirical and simulated resting-state BOLD-fMRI time courses parcellated using the HCP-MMP1 parcellation (Section S5). For the empirical data, a $TD$ matrix was calculated for each of the 255 HCP individuals (Section S4.1 and S4.3), and then an average $TD$ matrix was calculated across individuals. For the simulated data, we generated time series for 255 trials (to match the 255 HCP individuals of the empirical data) of the wave and mass models using their respective original optimized parameters (Section S12.5). A $TD$ matrix was calculated for each trial. Then, an average $TD$ matrix was calculated across trials. The average $TD$ matrices are shown in Fig. S19A. Finally, the three lag projections were obtained from the empirical and simulated average $TD$ matrices (Figs. S19B to S19D).

The results in Fig. S19B show that the mean lag pattern of the wave model is significantly correlated with the empirical pattern, whereas the mass model's mean lag pattern is not. For PC1 lag, the performance of the wave model slightly decreased but was still superior to the mass model (Fig. S19C). Correlations between model and empirical patterns for PC2 lag (Fig. S19D) were not significant, but the correlation was still higher for the wave model. These results show that, regardless of the method for calculating lag projections, the wave model captures the time-lagged properties of empirical fMRI data better than the mass model. This further highlights that wave dynamics can provide an accurate and physically mechanistic account of macroscale, resting-state dynamics, consistent with previous studies [102–104]. We also emphasize that the performance of these models in capturing lag structure is likely to be a conservative estimate, as the models rely on a simple and spatially uniform hemodynamic forward model that does not account for regional variations in neurovascular coupling [86,105,106]. Such variations are likely to strongly affect the empirically observed lags.

*S12.7 Modeling stimulus-evoked dynamics*

We also evaluated the degree to which the wave model could capture classical properties of evoked neural responses. Specifically, we used the optimized wave model (with $r_s = 28.9$ mm) to investigate neural dynamics in response to a stimulus applied to the left primary visual cortex (V1). In particular, the stimulus $Q(\boldsymbol{r}, t)$ was a 1 ms pulse (from $t = 1$ to 2 ms) with amplitude of 20 s$^{-1}$ (the results are robust to changes in the amplitude) restricted to vertices in the cortical surface that fall inside the V1 region defined by the HCP-MMP1 parcellation. We first decomposed $Q(\boldsymbol{r}, t)$ to calculate the time-varying amplitude $q_j(t)$ in Eq. (S19). We then calculated $\phi_j(t)$ by solving either Eq. (S21) or Eq. (S23) and subsequently combined $\phi_j(t)$ with the geometric eigenmodes via Eq. (S18) to obtain the spatiotemporal response $\phi(\boldsymbol{r}, t)$ at every location $\boldsymbol{r}$ and time $t$. We did the simulation over a 100 ms time period with 0.1 ms resolution. Finally, we parcellated $\phi(\boldsymbol{r}, t)$ into 180 regions using the HCP-MMP1 parcellation.

We compared the activity profile (amplitude vs time) of each brain region, focusing on the time for the activity to reach the peak amplitude (i.e., time to peak). Specifically, we investigated whether the temporal precedence of activity follows the human visual cortical hierarchy from visual to frontal cortices, which included the following brain regions: V1, V4, 7m, 7Am, TE1p, 7AL, 24dd, 2, 24dv, 8BM, 10r, 10v, 8BL, 10pp, 10d, a9-46v, and 9-46d. These regions closely resemble the areas previously identified in the macaque neocortex visual hierarchy using tract-tracing data and nonlinear network modeling [107].

We also compared the regional estimates of peak response times to T1w:T2w values, which is a noninvasive measure sensitive to intracortical myelin content [108] and a good proxy for cortical hierarchy rank [109]. The myelin map in fsLR-32k space was obtained from the HCP dataset [42,49] (https://balsa.wustl.edu/study/show/RVVG) and then parcellated using the HCP-MMP1 parcellation. We quantified the relationship between the regional values of time to peak and myelin content via Spearman rank correlation. The statistical significance of the correlation was assessed by comparing it with a null distribution of 10,000 correlation values obtained using a spatially constrained spin-test approach [110,111]. The approach calculates the correlation between one map and random spatially rotated versions of the other map, thereby preserving the spatial relationship of the parcels in the map. The resulting $p$-value, $p_{\text{spin}}$, is the fraction of null correlation values greater than the empirical correlation value. Finally, we repeated the above statistical test including all brain regions (i.e., not restricted to the visual hierarchy brain regions) to ensure that our findings were not driven by our particular selection of regions-of-interest. This is a more conservative test since not all brain regions show strong evoked responses to the visual stimulation.

S13. Estimating geometric eigenmodes of the thalamus, striatum, and hippocampus

We extended our eigenmode analysis to regions outside the neocortex focusing in particular on the subcortex (thalamus and striatum) and archicortex (hippocampus). Unlike the cortical ribbon, which can be modelled as a 2D sheet, these structures are solid three-dimensional (3D) objects. We therefore calculated the geometric eigenmodes using a tetrahedral mesh instead of the surface-based triangular mesh to account for the full 3D geometry of the non-neocortical structures, as outlined below.

We first used the probabilistic Harvard–Oxford subcortical atlas (https://fsl.fmrib.ox.ac.uk/fsl/fslwiki/Atlases) to generate a volumetric binary mask in each hemisphere for the thalamus, striatum, and hippocampus. Voxels with a probability of 25% or more of belonging to these structures were included in the masks. We used FreeSurfer's mri_mc function, which implements a marching cubes algorithm to construct a 2D surface by tessellating the volumetric masks, followed by the Gmsh software (https://gmsh.info/) to convert the 2D surface into a 3D tetrahedral mesh. We then used the LaPy python library, as in Section S2, to solve Eq. (S1) on the tetrahedral mesh of each non-neocortical structure to obtain the eigenmodes and their corresponding eigenvalues. Finally, we projected the eigenmodes in tetrahedral space back into the natural volumetric space via interpolation. Hence, the resulting eigenmodes spatially vary through the 3D voxels comprising each non-neocortical structure's volume. For this part of the study, we discarded the first constant mode and used the next 20 modes in our analyses.

S14. Mapping the functional organization of the thalamus, striatum, and hippocampus

The signal-to-noise ratio of fMRI is generally weaker in non-neocortical structures than in the neocortex [112]. Moreover, fine-grained task activations are often hard to resolve in these smaller structures due to the limited resolution of fMRI and the spatial smoothing induced by common fMRI processing methods. Thus, to efficiently map the functional organization of the thalamus, striatum, and hippocampus, we used connectopic mapping [113] of the resting-state fMRI signals in each structure to obtain their dominant functional modes (often called gradients although the patterns simply capture spatial variations in point-wise similarities of FC profiles and are not gradients themselves). This technique, and related procedures, has been extensively used in past work to study the functional organization of these structures [114–116].

We applied connectopic mapping to the volumetric voxel-wise resting-state fMRI data of the HCP individuals in Section S4.3, following the procedure described by [113]. Specifically, for each individual and non-neocortical ROI (i.e., thalamus, striatum, and hippocampus), we constructed a ROI time series data matrix, $A$, of size $T \times N$, where $T$ is the number of time frames and $N$ is the number of voxels in the ROI. Similarly, for each individual, we constructed the gray matter time-series data matrix, $B$, of size $T \times M$, where $M$ is the number of gray matter voxels outside the ROI. Since $M$ is generally large (i.e., $\geq 10^3$), we reduced the dimensionality of $B$ using singular value decomposition (SVD) to construct the matrix $\tilde{B}$ of size $T \times (T-1)$. The connectivity fingerprint of every voxel within the ROI was then calculated using the Pearson correlation of $A$ and $\tilde{B}$ to obtain the matrix, $C$, of size $N \times (T-1)$. Similarity in the connectivity fingerprints between each pair of ROI voxels was calculated using the $\eta^2$ coefficient [117], resulting in the matrix, $S$, of size $N \times N$. The $\eta^2$ coefficient represents the fraction of the variance in one connectivity profile accounted for by the variance in another. We then took the average $S$ across all individuals. A nonlinear manifold learning procedure using the Laplacian Eigenmaps algorithm [118] was applied to the $S$ matrix of each ROI to calculate its eigenvectors and eigenvalues. The eigenvectors represent the functional patterns, termed functional gradients, of the ROI, ordered according to the variance in FC similarity they explain. We only analyzed the first 20 non-constant gradients to enable direct comparisons with the geometric eigenmodes described in Section S13. Typical applications of FC-based gradient analysis rarely consider more than the first five gradients [113,119].

We analyzed the correspondence between the geometric eigenmodes and functional gradients in each non-neocortical structure by taking their absolute spatial correlations, given that the signs of the modes and gradients are arbitrary. Moreover, the ordering of the geometric eigenmodes within the same eigengroup can change (Section S2 and Table S1; e.g., the first eigengroup can have order flips among modes 2 to 4), hence one-to-one correspondence between the indices of geometric eigenmodes and functional gradients is not guaranteed. We therefore examined all possible pairwise correlations between the modes and gradients, and evaluated correspondence with respect to the mode that maximally correlated with each gradient. The maximal order difference observed between geometric modes and functional gradients were 1, 8, and 5 out of 20 for the thalamus, striatum, and hippocampus, respectively.

S15. Data and code availability

Raw and preprocessed HCP data can be accessed at https://db.humanconnectome.org/. All source data and computer codes to calculate the eigenmodes, analyze results, and generate the main and supplementary figures of this study are openly available at https://github.com/BMHLab/BrainEigenmodes.

**REFERENCES**


1. Beurle, R. L. Properties of a mass of cells capable of regenerating pulses. *Philosophical Transactions of the Royal Society of London. Series B, Biological Sciences* **240**, 55–94 (1956).
2. Lopes da Silva, F. H., van Rotterdam, A., Barts, P., van Heusden, E. & Burr, W. Models of Neuronal Populations: The Basic Mechanisms of Rhythmicity. *Progress in Brain Research* **45**, 281–308 (1976).
3. Wright, J. J. & Liley, D. T. J. Simulation of electrocortical waves. *Biological Cybernetics* **72**, 347–356 (1995).
4. Deco, G., Jirsa, V. K., Robinson, P. A., Breakspear, M. & Friston, K. The dynamic brain: From spiking neurons to neural masses and cortical fields. *PLoS Computational Biology* **4**, (2008).
5. Jirsa, V. & Haken, H. Field Theory of Electromagnetic Brain Activity. *Physical Review Letters* **77**, 960–963 (1996).
6. Robinson, P. A., Rennie, C. J. & Wright, J. J. Propagation and stability of waves of electrical activity in the cerebral cortex. *Physical Review E* **56**, 826–840 (1997).
7. Robinson, P. A., Rennie, C. J., Rowe, D. L., O'Connor, S. C. & Gordon, E. Multiscale brain modelling. *Philosophical Transactions of the Royal Society B: Biological Sciences* **360**, 1043–1050 (2005).
8. Sanz-Leon, P. *et al.* NFTsim: Theory and Simulation of Multiscale Neural Field Dynamics. *PLoS Computational Biology* **14**, e1006387 (2018).
9. Braitenberg, V. & Schüz, A. *Cortex: Statistics and Geometry of Neuronal Connectivity*. *Cortex: Statistics and Geometry of Neuronal Connectivity* (1998). doi:10.1007/978-3-662-03733-1.
10. Henderson, J. A. & Robinson, P. A. Relations between the geometry of cortical gyrification and white-matter network architecture. *Brain Connectivity* **4**, 112–130 (2014).
11. Robinson, P. A. Physical brain connectomics. *Physical Review E* **99**, 012421 (2019).
12. Robinson, P. A. Interrelating anatomical, effective, and functional brain connectivity using propagators and neural field theory. *Physical Review E* **85**, (2012).
13. Nunez, P. L. The brain wave equation: a model for the EEG. *Mathematical Biosciences* **21**, 279–297 (1974).
14. Robinson, P. A. *et al.* Prediction of electroencephalographic spectra from neurophysiology. *Physical Review E* **63**, 021903 (2001).

15. Pang, J. C. & Robinson, P. A. Neural mechanisms of the EEG alpha-BOLD anticorrelation. *NeuroImage* **181**, 461–470 (2018).
16. Rennie, C. J., Robinson, P. A. & Wright, J. J. Unified neurophysical model of EEG spectra and evoked potentials. *Biological Cybernetics* **86**, 457–471 (2002).
17. Mukta, K. N., Robinson, P. A., Pagès, J. C., Gabay, N. C. & Gao, X. Evoked response activity eigenmode analysis in a convoluted cortex via neural field theory. *Physical Review E* **102**, (2020).
18. Abeysuriya, R. G., Rennie, C. J. & Robinson, P. A. Physiologically based arousal state estimation and dynamics. *Journal of Neuroscience Methods* **253**, 55–69 (2015).
19. Assadzadeh, S. & Robinson, P. A. Necessity of the sleep-wake cycle for synaptic homeostasis: System-level analysis of plasticity in the corticothalamic system. *Royal Society Open Science* **5**, (2018).
20. Gabay, N. C., Babaie-Janvier, T. & Robinson, P. A. Dynamics of cortical activity eigenmodes including standing, traveling, and rotating waves. *Physical Review E* **98**, 042413 (2018).
21. Tokariev, A. *et al.* Large-scale brain modes reorganize between infant sleep states and carry prognostic information for preterms. *Nature Communications* **10**, 2619 (2019).
22. Nozari, E. *et al.* Is the brain macroscopically linear? A system identification of resting state dynamics. *arXiv* (2020).
23. Robinson, P. A. *et al.* Eigenmodes of brain activity: Neural field theory predictions and comparison with experiment. *NeuroImage* **142**, 79–98 (2016).
24. Robinson, P. A. *et al.* Determination of Dynamic Brain Connectivity via Spectral Analysis. *Frontiers in Human Neuroscience* **15**, (2021).
25. Gabay, N. C. & Robinson, P. A. Cortical geometry as a determinant of brain activity eigenmodes: Neural field analysis. *Physical Review E* **96**, (2017).
26. Nunez, P. L. *Neocortical Dynamics and Human EEG Rhythms*. (Oxford University Press, 1995).
27. Bressloff, P. C. Spatiotemporal dynamics of continuum neural fields. *Journal of Physics A: Mathematical and Theoretical* **45**, 033001 (2012).
28. Coombes, S., Beim Graben, P., Potthast, R. & Wright, J. J. *Neural fields: Theory and applications*. vol. 9783642545 (Springer, 2014).
29. Bick, C., Goodfellow, M., Laing, C. R. & Martens, E. A. Understanding the dynamics of biological and neural oscillator networks through exact mean-field reductions: a review. *Journal of Mathematical Neuroscience* **10**, (2020).
30. Melrose, D. B. & McPhedran, R. C. *Electromagnetic Processes in Dispersive Media*. (Cambridge University Press, 1991).
31. Wachinger, C., Golland, P., Kremen, W., Fischl, B. & Reuter, M. BrainPrint: A discriminative characterization of brain morphology. *NeuroImage* **109**, 232–248 (2015).
32. Chavel, I. *Eigenvalues in Riemannian geometry*. vol. 115 (1984).
33. Seo, S. & Chung, M. K. Laplace-Beltrami eigenfunction expansion of cortical manifolds. in *Proceedings - International Symposium on Biomedical Imaging* 372–375 (2011). doi:10.1109/ISBI.2011.5872426.
34. Reuter, M., Wolter, F. E. & Peinecke, N. Laplace-Beltrami spectra as 'Shape-DNA' of surfaces and solids. *CAD Computer Aided Design* **38**, 342–366 (2006).

35. Goscinski, W. J. *et al.* The multi-modal Australian ScienceS Imaging and Visualization Environment (MASSIVE) high performance computing infrastructure: applications in neuroscience and neuroinformatics research. *Frontiers in Neuroinformatics* **8**, (2014).
36. Fischl, B., Sereno, M. I., Tootell, R. B. H. & Dale, A. M. High-resolution intersubject averaging and a coordinate system for the cortical surface. *Human Brain Mapping* **8**, 272–284 (1999).
37. Shuman, D. I., Narang, S. K., Frossard, P., Ortega, A. & Vandergheynst, P. The emerging field of signal processing on graphs. *IEEE Signal Processing Magazine* **30**, 83–98 (2013).
38. Henderson, J. A., Aquino, K. M. & Robinson, P. A. Empirical estimation of the eigenmodes of macroscale cortical dynamics: Reconciling neural field eigenmodes and resting-state networks. *Neuroimage: Reports* **2**, 100103 (2022).
39. Chen, Y.-C. *et al.* The individuality of shape asymmetries of the human cerebral cortex. *eLife* **11**, e75056 (2022).
40. Courant, R. & Hilbert, D. Methods of Mathematical Physics. *Methods of Mathematical Physics* **1**, 1–560 (2007).
41. Eickhoff, S. B., Yeo, B. T. T. & Genon, S. Imaging-based parcellations of the human brain. *Nature Reviews Neuroscience* **19**, 672–686 (2018).
42. van Essen, D. C. *et al.* The WU-Minn Human Connectome Project: An overview. *NeuroImage* **80**, 62–79 (2013).
43. Barch, D. M. *et al.* Function in the human connectome: Task-fMRI and individual differences in behavior. *NeuroImage* **80**, 169–189 (2013).
44. Woolrich, M. W., Behrens, T. E. J., Beckmann, C. F., Jenkinson, M. & Smith, S. M. Multilevel linear modelling for FMRI group analysis using Bayesian inference. *NeuroImage* **21**, 1732–1747 (2004).
45. Robinson, E. C. *et al.* Multimodal surface matching with higher-order smoothness constraints. *NeuroImage* **167**, 453–465 (2018).
46. Smith, S. *et al.* The minimal preprocessing pipelines for the Human Connectome Project. *NeuroImage* **80**, 105–124 (2013).
47. Salimi-Khorshidi, G. *et al.* Automatic denoising of functional MRI data: combining independent component analysis and hierarchical fusion of classifiers. *Neuroimage* **15**, 449–468 (2013).
48. Mansour L, S., Tian, Y., Yeo, B. T. T., Cropley, V. & Zalesky, A. High-resolution connectomic fingerprints: Mapping neural identity and behavior. *NeuroImage* **229**, 117695 (2021).
49. Glasser, M. F. *et al.* A multi-modal parcellation of human cerebral cortex. *Nature* **536**, 171–178 (2016).
50. Schaefer, A. *et al.* Local-global parcellation of the human cerebral cortex from intrinsic functional connectivity mri. *Cerebral Cortex* **28**, 3095–3114 (2018).
51. Naze, S., Proix, T., Atasoy, S. & Kozloski, J. R. Robustness of connectome harmonics to local gray matter and long-range white matter connectivity changes: Sensitivity analysis of Connectome Harmonics. *NeuroImage* **224**, 117364 (2021).
52. Atasoy, S., Donnelly, I. & Pearson, J. Human brain networks function in connectome-specific harmonic waves. *Nature Communications* **7**, 10340 (2016).
53. Preti, M. G. & Van De Ville, D. Decoupling of brain function from structure reveals regional behavioral specialization in humans. *Nature Communications* **10**, 4747 (2019).

54. Rué-Queralt, J. *et al.* The connectome spectrum as a canonical basis for a sparse representation of fast brain activity. *NeuroImage* **244**, 118611 (2021).
55. Tournier, J.-D. *et al.* MRtrix3: A fast, flexible and open software framework for medical image processing and visualisation. *NeuroImage* **202**, 116137 (2019).
56. Lévy, B. Laplace-beltrami eigenfunctions towards an algorithm that 'understands' geometry. in *Proceedings - IEEE International Conference on Shape Modeling and Applications 2006, SMI 2006* vol. 2006 13 (2006).
57. Zalesky, A. *et al.* Whole-brain anatomical networks: Does the choice of nodes matter? *NeuroImage* **50**, 970–983 (2010).
58. Fornito, A., Zalesky, A. & Bullmore, E. T. *Fundamentals of Brain Network Analysis*. (2016).
59. Theodoni, P. *et al.* Structural Attributes and Principles of the Neocortical Connectome in the Marmoset Monkey. *Cerebral Cortex* **32**, 15–28 (2022).
60. Smith, S. M., Hyvärinen, A., Varoquaux, G., Miller, K. L. & Beckmann, C. F. Group-PCA for very large fMRI datasets. *NeuroImage* **101**, 738–749 (2014).
61. Do Carmo, M. P. *Differential Geometry of Curves and Surfaces: Revised and Updated Second Edition*. (Dover Publications, 2016).
62. Klingenberg, W. P. A. *Riemannian Geometry*. (de Gruyter, 1995).
63. Reuter, M. *Laplace Spectra for Shape Recognition*. (Books On Demand, 2006).
64. Strichartz, R. S. Harmonic analysis as spectral theory of Laplacians. *Journal of Functional Analysis* **87**, 51–148 (1989).
65. Chung, F. R. K. *Spectral Graph Theory*. (American Mathematical Society, 1996).
66. Gorgolewski, K. J. *et al.* NeuroVault.Org: A web-based repository for collecting and sharing unthresholded statistical maps of the human brain. *Frontiers in Neuroinformatics* **9**, (2015).
67. Abraham, A. *et al.* Machine learning for neuroimaging with scikit-learn. *Frontiers in Neuroinformatics* **8**, (2014).
68. Carp, J. The secret lives of experiments: Methods reporting in the fMRI literature. *NeuroImage* **63**, 289–300 (2012).
69. Coalson, T. S., Van Essen, D. C. & Glasser, M. F. The impact of traditional neuroimaging methods on the spatial localization of cortical areas. *Proceedings of the National Academy of Sciences of the United States of America* **115**, E6356–E6365 (2018).
70. Arfken, G. *Mathematical Methods for Physicists*. (Academic Press Inc, 1985).
71. Nunez, P. L. The brain wave equation: a model for the EEG. *Mathematical Biosciences* **21**, 279–297 (1974).
72. Jirsa, V. & Haken, H. Field Theory of Electromagnetic Brain Activity. *Physical Review Letters* **77**, 960–963 (1996).
73. Breakspear, M. Dynamic models of large-scale brain activity. *Nature Neuroscience* **20**, 340–352 (2017).
74. Sanz-Leon, P. *et al.* The virtual brain: A simulator of primate brain network dynamics. *Frontiers in Neuroinformatics* **7**, 10 (2013).
75. Breakspear, M., Heitmann, S. & Daffertshofer, A. Generative models of cortical oscillations: Neurobiological implications of the Kuramoto model. *Frontiers in Human Neuroscience* **4**, 190 (2010).
76. Wilson, H. R. & Cowan, J. D. Excitatory and Inhibitory Interactions in Localized Populations of Model Neurons. *Biophysical Journal* **12**, 1–24 (1972).
77. Deco, G. *et al.* How local excitation-inhibition ratio impacts the whole brain dynamics. *Journal of Neuroscience* **34**, 7886–7898 (2014).

78. Demirtaş, M. *et al.* Hierarchical Heterogeneity across Human Cortex Shapes Large-Scale Neural Dynamics. *Neuron* **101**, 1181–1194 (2019).
79. Deco, G. *et al.* Dynamical consequences of regional heterogeneity in the brain's transcriptional landscape. *Science Advances* **7**, eabf4752 (2021).
80. Aquino, K. M. *et al.* On the intersection between data quality and dynamical modelling of large-scale fMRI signals. *NeuroImage* **256**, 119051 (2022).
81. Behrens, T. E. J. *et al.* Characterization and Propagation of Uncertainty in Diffusion-Weighted MR Imaging. *Magnetic Resonance in Medicine* **50**, 1077–1088 (2003).
82. Spiegler, A. & Jirsa, V. Systematic approximations of neural fields through networks of neural masses in the virtual brain. *NeuroImage* **83**, 704–725 (2013).
83. Stephan, K. E., Weiskopf, N., Drysdale, P. M., Robinson, P. A. & Friston, K. J. Comparing hemodynamic models with DCM. *NeuroImage* **38**, 387–401 (2007).
84. Aquino, K. M., Schira, M. M., Robinson, P. A., Drysdale, P. M. & Breakspear, M. Hemodynamic traveling waves in human visual cortex. *PLoS Computational Biology* **8**, (2012).
85. Pang, J. C., Robinson, P. A., Aquino, K. M. & Vasan, N. Effects of astrocytic dynamics on spatiotemporal hemodynamics: Modeling and enhanced data analysis. *NeuroImage* **147**, 994–1005 (2017).
86. Pang, J. C., Aquino, K. M., Robinson, P. A., Lacy, T. C. & Schira, M. M. Biophysically based method to deconvolve spatiotemporal neurovascular signals from fMRI data. *Journal of Neuroscience Methods* **308**, 6–20 (2018).
87. Deco, G., Jirsa, V., McIntosh, A. R., Sporns, O. & Kötter, R. Key role of coupling, delay, and noise in resting brain fluctuations. *Proceedings of the National Academy of Sciences of the United States of America* **106**, 10302–10307 (2009).
88. Cabral, J., Kringelbach, M. L. & Deco, G. Exploring the network dynamics underlying brain activity during rest. *Progress in Neurobiology* **114**, 102–131 (2014).
89. Heinzle, J., Koopmans, P. J., den Ouden, H. E. M., Raman, S. & Stephan, K. E. A hemodynamic model for layered BOLD signals. *NeuroImage* **125**, 556–570 (2016).
90. Deco, G., Kringelbach, M. L., Jirsa, V. K. & Ritter, P. The dynamics of resting fluctuations in the brain: metastability and its dynamical cortical core. *Sci Rep* **7**, 3095 (2017).
91. Glerean, E., Salmi, J., Lahnakoski, J. M., Jääskeläinen, I. P. & Sams, M. Functional Magnetic Resonance Imaging Phase Synchronization as a Measure of Dynamic Functional Connectivity. *Brain Connectivity* **2**, 91–101 (2012).
92. Pang, J. C. & Robinson, P. A. Power spectrum of resting-state blood-oxygen-level-dependent signal. *Physical Review E* **100**, 022418 (2019).
93. Beaumont, M. A., Cornuet, J. M., Marin, J. M. & Robert, C. P. Adaptive approximate Bayesian computation. *Biometrika* **96**, 983–990 (2009).
94. Turner, B. M. & Van Zandt, T. Hierarchical Approximate Bayesian Computation. *Psychometrika* **79**, 185–209 (2014).
95. Mitra, A., Snyder, A. Z., Hacker, C. D. & Raichle, M. E. Lag structure in resting-state fMRI. *Journal of Neurophysiology* **111**, 2374–2391 (2014).
96. Mitra, A., Snyder, A. Z., Blazey, T. & Raichle, M. E. Lag threads organize the brain's intrinsic activity. *Proceedings of the National Academy of Sciences* **112**, E2235–E2244 (2015).

97. Cabral, J., Kringelbach, M. L. & Deco, G. Functional connectivity dynamically evolves on multiple time-scales over a static structural connectome: Models and mechanisms. *NeuroImage* **160**, 84–96 (2017).
98. Kashyap, A. & Keilholz, S. Dynamic properties of simulated brain network models and empirical resting-state data. *Network Neuroscience* **3**, 405–426 (2019).
99. Bolt, T. *et al.* A parsimonious description of global functional brain organization in three spatiotemporal patterns. *Nature Neuroscience* **25**, 1093–1103 (2022).
100. He, B. J., Zempel, J. M., Snyder, A. Z. & Raichle, M. E. The Temporal Structures and Functional Significance of Scale-free Brain Activity. *Neuron* **66**, 353–369 (2010).
101. Raut, R. V. *et al.* Global waves synchronize the brain's functional systems with fluctuating arousal. *Science Advances* **7**, (2021).
102. Majeed, W. *et al.* Spatiotemporal dynamics of low frequency BOLD fluctuations in rats and humans. *NeuroImage* **54**, 1140–1150 (2011).
103. Matsui, T., Murakami, T. & Ohki, K. Transient neuronal coactivations embedded in globally propagating waves underlie resting-state functional connectivity. *Proceedings of the National Academy of Sciences* **113**, 6556–6561 (2016).
104. Chan, A. W., Mohajerani, M. H., LeDue, J. M., Wang, Y. T. & Murphy, T. H. Mesoscale infraslow spontaneous membrane potential fluctuations recapitulate high-frequency activity cortical motifs. *Nature Communications* **6**, 7738 (2015).
105. Aquino, K. M., Schira, M. M., Robinson, P. A., Drysdale, P. M. & Breakspear, M. Hemodynamic traveling waves in human visual cortex. *PLoS Computational Biology* **8**, (2012).
106. Pang, J. C., Robinson, P. A., Aquino, K. M. & Vasan, N. Effects of astrocytic dynamics on spatiotemporal hemodynamics: Modeling and enhanced data analysis. *NeuroImage* **147**, 994–1005 (2017).
107. Chaudhuri, R., Knoblauch, K., Gariel, M. A., Kennedy, H. & Wang, X. J. A Large-Scale Circuit Mechanism for Hierarchical Dynamical Processing in the Primate Cortex. *Neuron* **88**, 419–431 (2015).
108. Glasser, M. F. & Van Essen, D. C. Mapping Human Cortical Areas In Vivo Based on Myelin Content as Revealed by T1- and T2-Weighted MRI. *Journal of Neuroscience* **31**, 11597–11616 (2011).
109. Burt, J. B. *et al.* Hierarchy of transcriptomic specialization across human cortex captured by structural neuroimaging topography. *Nature Neuroscience* **21**, 1251–1259 (2018).
110. Váša, F. *et al.* Adolescent tuning of association cortex in human structural brain networks. *Cerebral Cortex* **28**, 281–294 (2018).
111. Alexander-Bloch, A. F. *et al.* On testing for spatial correspondence between maps of human brain structure and function. *NeuroImage* **178**, 540–551 (2018).
112. Uğurbil, K. *et al.* Pushing spatial and temporal resolution for functional and diffusion MRI in the Human Connectome Project. *NeuroImage* **80**, 80–104 (2013).
113. Haak, K. V., Marquand, A. F. & Beckmann, C. F. Connectopic mapping with resting-state fMRI. *NeuroImage* **170**, 83–94 (2018).
114. Vos de Wael, R. *et al.* Anatomical and microstructural determinants of hippocampal subfield functional connectome embedding. *Proceedings of the National Academy of Sciences* **115**, 10154–10159 (2018).
115. Yang, S. *et al.* The thalamic functional gradient and its relationship to structural basis and cognitive relevance. *NeuroImage* **218**, 116960 (2020).

116. Oldehinkel, M. *et al.* Mapping dopaminergic projections in the human brain with resting-state fMRI. *eLife* **11**, e71846 (2022).
117. Cohen, A. L. *et al.* Defining functional areas in individual human brains using resting functional connectivity MRI. *NeuroImage* **41**, 45–57 (2008).
118. Belkin, M. & Niyogi, P. Laplacian eigenmaps for dimensionality reduction and data representation. *Neural computation* **15**, 1373–1396 (2003).
119. Margulies, D. S. *et al.* Situating the default-mode network along a principal gradient of macroscale cortical organization. *Proceedings of the National Academy of Sciences of the United States of America* **113**, 12574–12579 (2016).

### SUPPLEMENTARY FIGURES

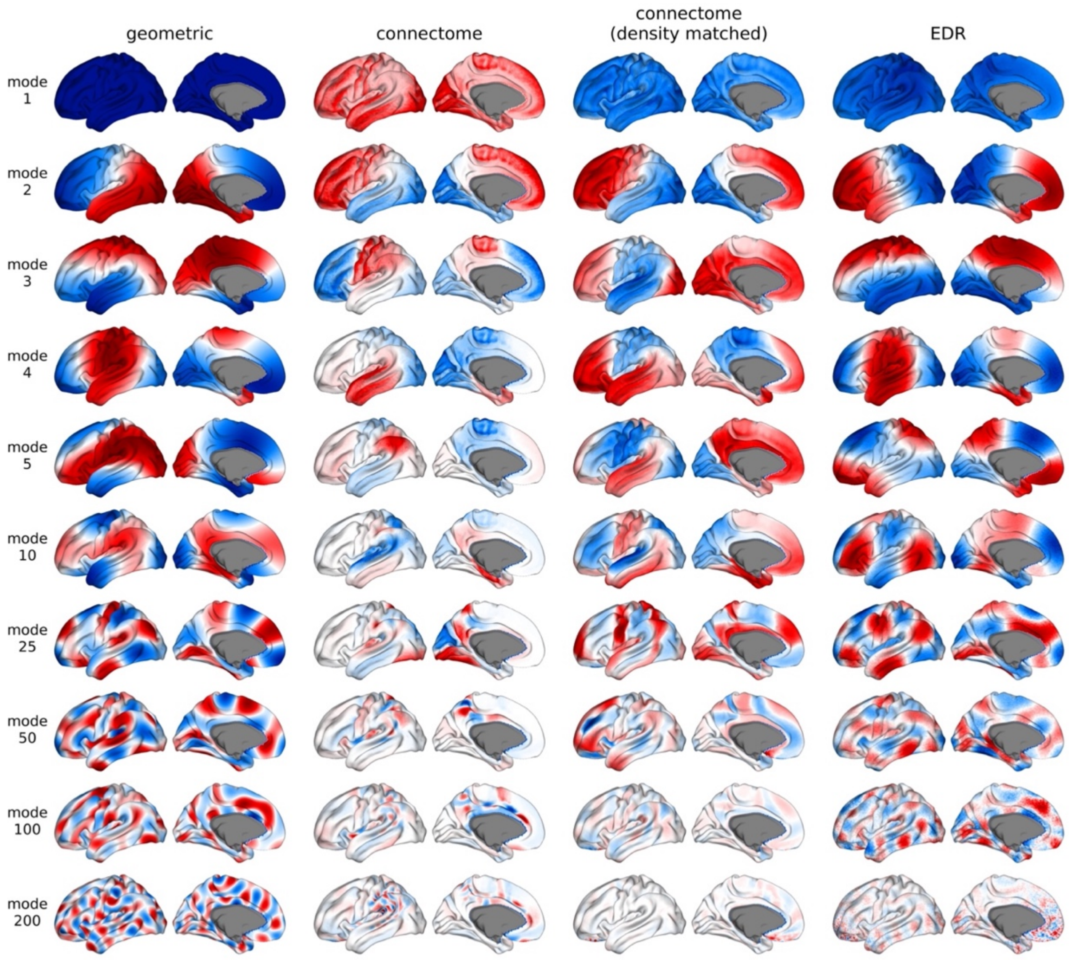


**Fig. S1. Eigenmode basis sets.** The basis sets from left to right are geometric eigenmodes, connectome eigenmodes, connectome eigenmodes using a connectivity matrix matching the density used by the exponential distance rule (EDR) eigenmodes, and EDR eigenmodes. Negative–zero–positive values are colored as blue–white–red.

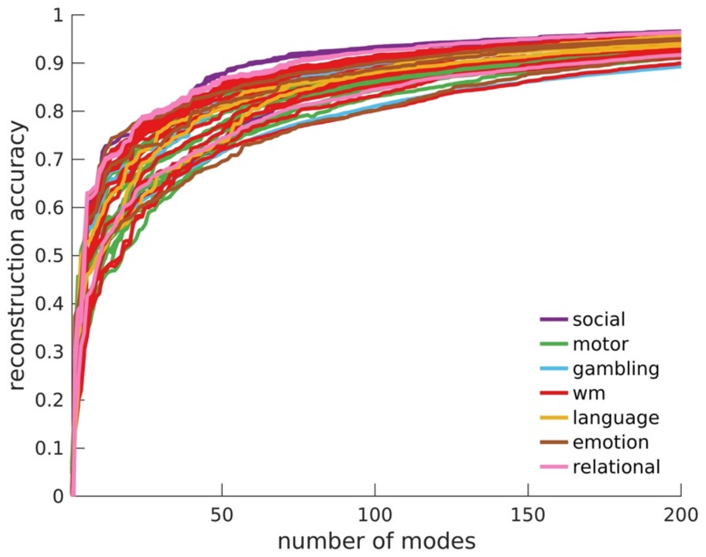


**Fig. S2. Reconstruction accuracy of 47 HCP task-contrast maps obtained with geometric eigenmodes.** The lines are colored according to the groups defined by the 7 broad HCP task types (Section S4.2 and Table S2). wm = working memory.

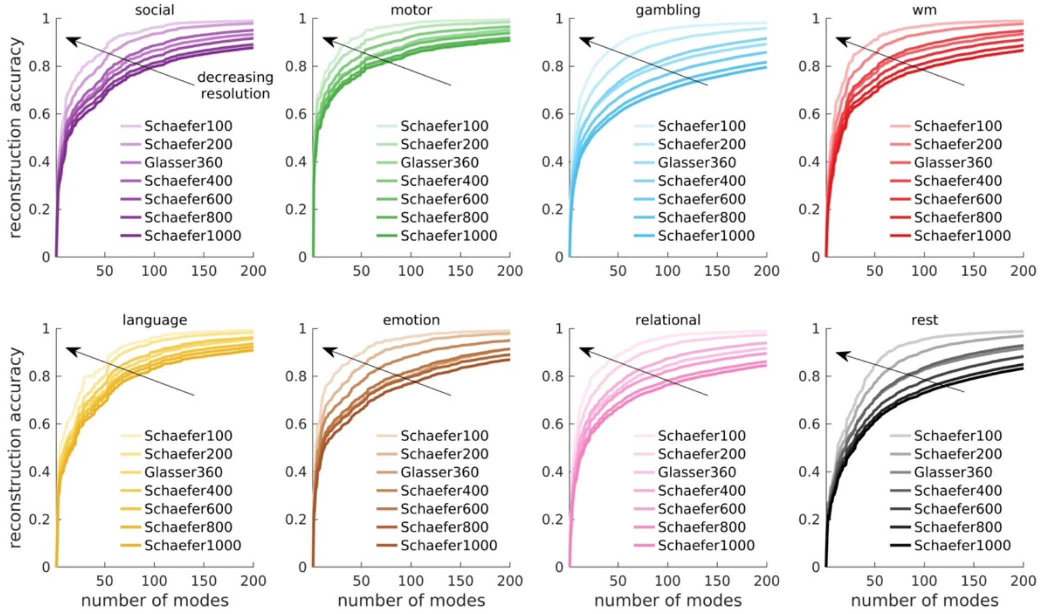


**Fig. S3. Reconstruction accuracy of 7 key HCP task-contrast maps and resting-state FC for different parcellation resolutions.** See Section S4.2 and Table S2 for details about the contrast maps. wm = working memory. The lines with dark to light colors represent decreasing parcellation resolutions (direction of arrows); i.e., Schaefer100, Schaefer200, Glasser360, Schaefer400, Schaefer600, Schaefer800, and Schaefer1000 has 100, 200, 360, 400, 600, 800, and 1000 parcels, respectively, across both hemispheres.

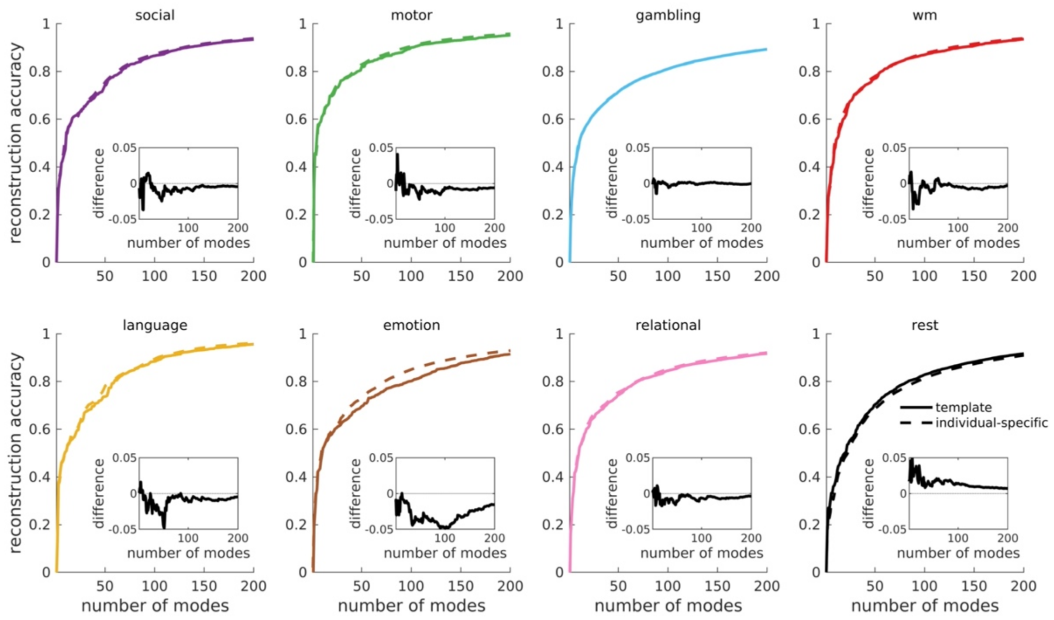


**Fig. S4. Reconstruction accuracy of 7 key HCP task-contrast maps and resting-state FC using template-derived and individual-specific geometric eigenmodes.** See Section S4.2 and Table S2 for details about the contrast maps. wm = working memory. The solid lines represent results achieved by template eigenmodes derived from a template surface (Fig. 1A). The dashed lines represent results achieved by individual-specific eigenmodes derived from individual-specific surfaces. The insets show the difference between the two results (i.e., template minus individual-specific).

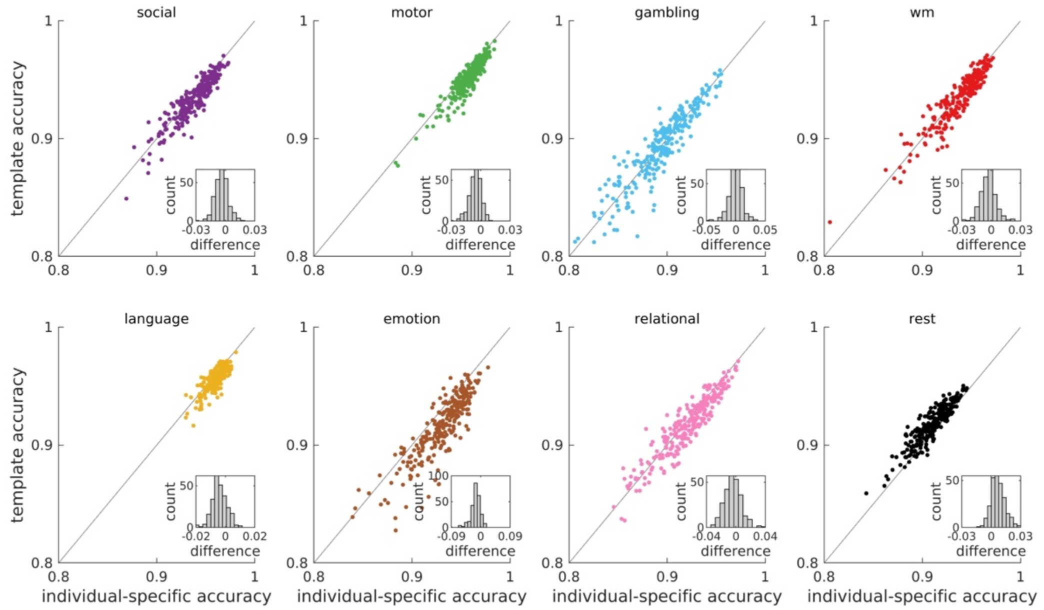


**Fig. S5. Individual-based reconstruction accuracy of 7 key HCP task-contrast maps and resting-state FC using 200 template-derived and individual-specific geometric eigenmodes.** See Section S4.2 and Table S2 for details about the contrast maps. wm = working memory. Template eigenmodes were derived from a template surface, while individual-specific eigenmodes were derived from individual-specific surfaces. Each point corresponds to an individual. The dotted lines represent the template accuracy = individual-specific accuracy lines. Points above the dotted lines mean that template accuracy > individual-specific accuracy. The insets show the histogram of the difference between the reconstruction accuracy achieved by template eigenmodes and individual-specific eigenmodes (i.e., template minus individual-specific).

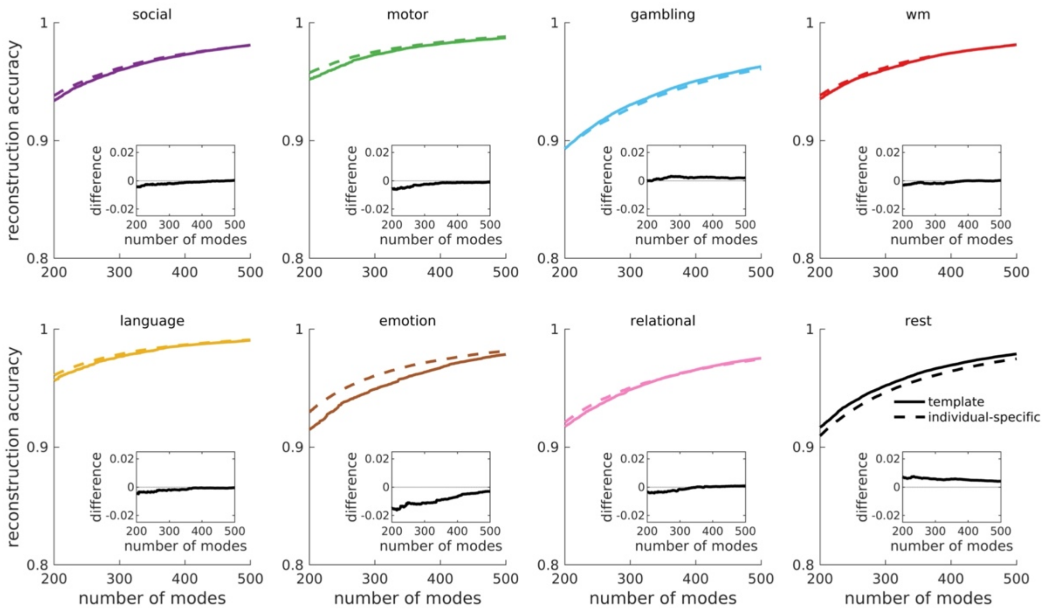


**Fig. S6. Reconstruction accuracy of 7 key HCP task-contrast maps and resting-state FC using 200 to 500 template-derived and individual-specific geometric eigenmodes.** See Section S4.2 and Table S2 for details about the contrast maps. wm = working memory. The solid lines represent results achieved by template eigenmodes derived from a template surface (Fig. 1A). The dashed lines represent results achieved by individual-specific eigenmodes derived from individual-specific surfaces. The insets show the difference between the two results (i.e., template minus individual-specific).

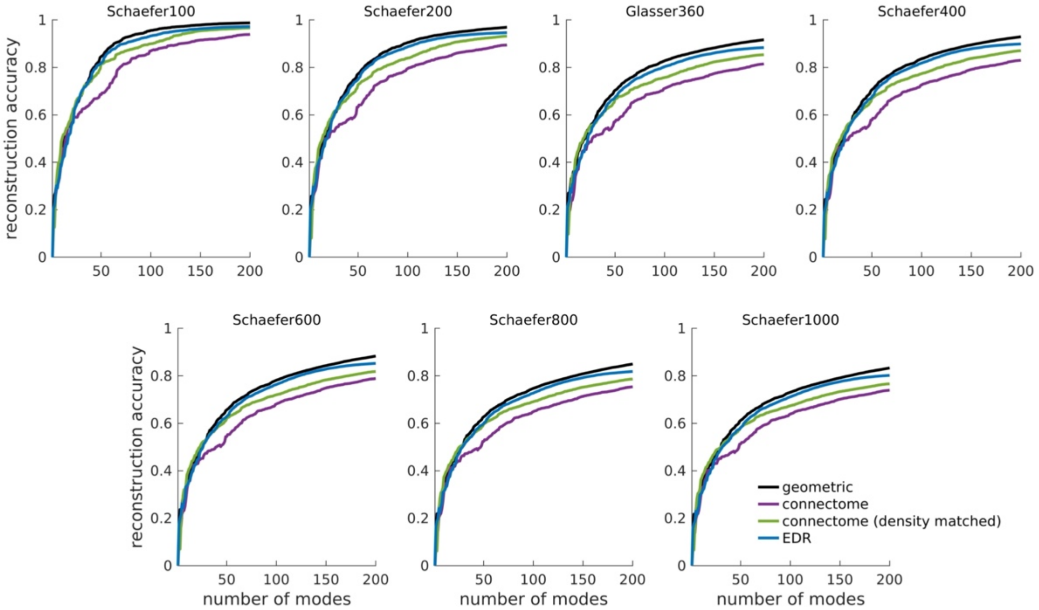


**Fig. S7. Reconstruction accuracy of resting-state FC achieved by different basis sets for different parcellation resolutions.** The basis sets are geometric eigenmodes, connectome eigenmodes, connectome eigenmodes using a connectivity matrix with density matched to that used by the EDR eigenmodes, and EDR eigenmodes. Schaefer100, Schaefer200, Glasser360, Schaefer400, Schaefer600, Schaefer800, and Schaefer1000 has 100, 200, 360, 400, 600, 800, and 1000 parcels, respectively, across both hemispheres.

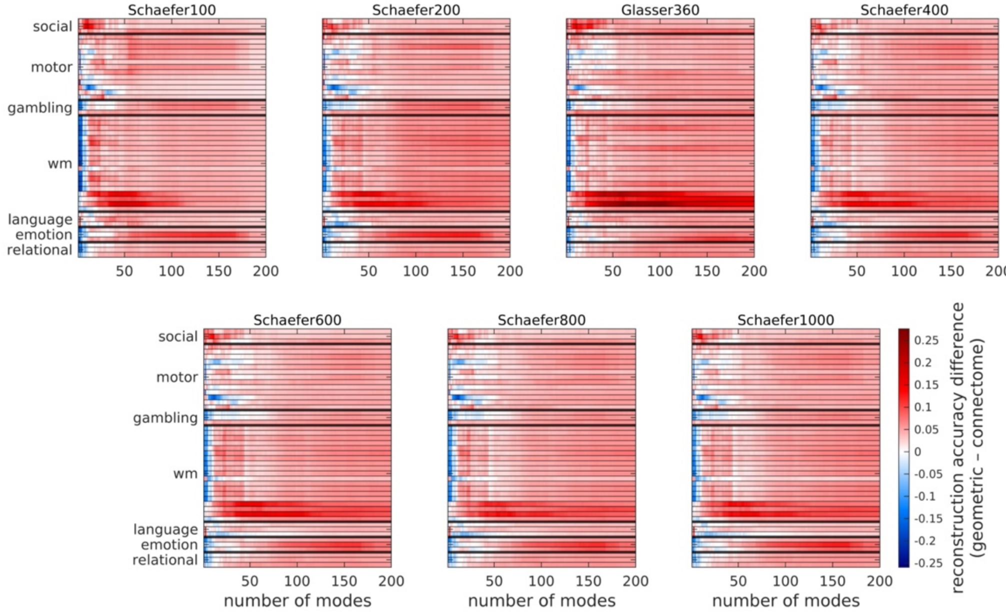


**Fig. S8. Difference in reconstruction accuracy of all 47 HCP task-contrast maps achieved by geometric eigenmodes and connectome eigenmodes for different parcellation resolutions.** Each row represents a different task contrast, which have been grouped here by broad types (Section S4.2 and Table S2). wm = working memory. Red indicates superior performance for geometric eigenmodes. Schaefer100, Schaefer200, Glasser360,

Schaefer400, Schaefer600, Schaefer800, and Schaefer1000 has 100, 200, 360, 400, 600, 800, and 1000 parcels, respectively, across both hemispheres.

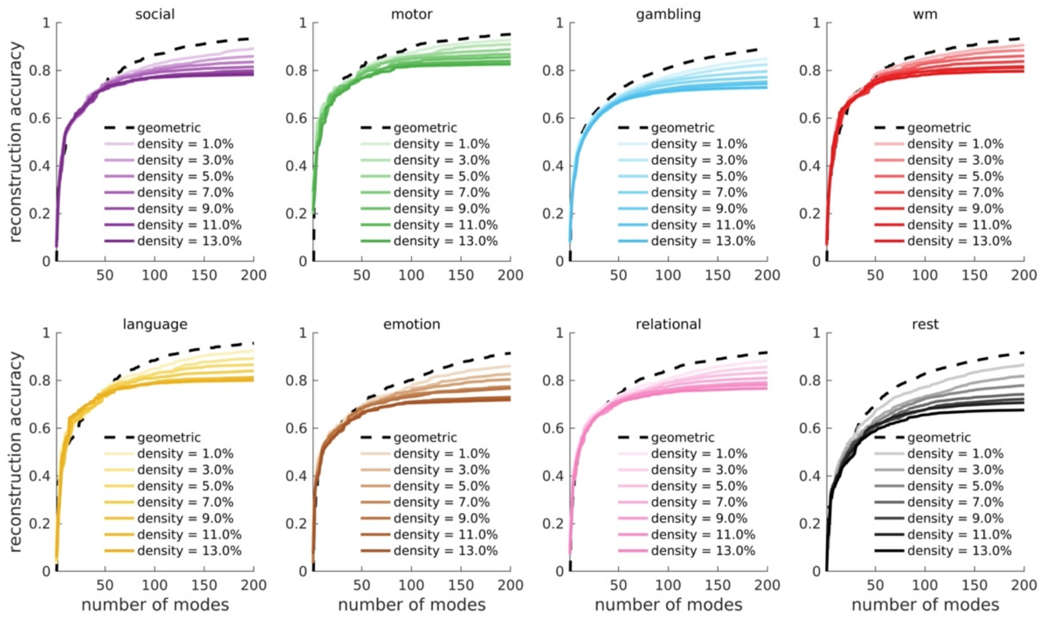


**Fig. S9. Reconstruction accuracy of 7 key HCP task-contrast maps and resting-state FC achieved by geometric eigenmodes and connectome eigenmodes of varying connectome densities.** See Section S4.2 and Table S2 for details about the contrast maps. wm = working memory. The dashed lines represent results achieved by geometric eigenmodes (Fig. 2A). The lines with light to dark colors represent results achieved by connectome eigenmodes using a connectivity matrix of increasing densities (from 1.0% to 13.0%).

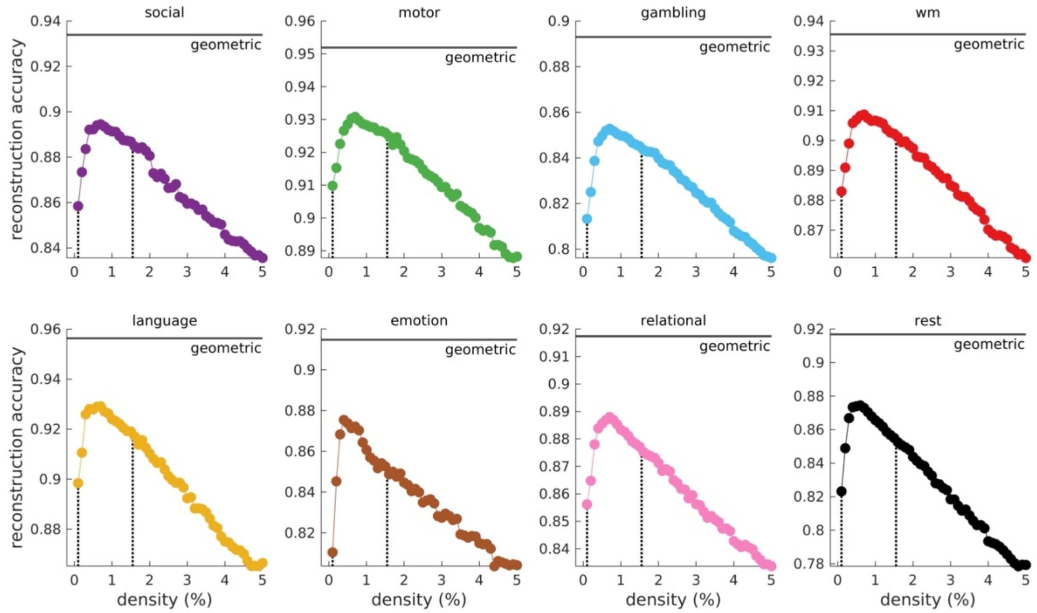

**Fig. S10. Reconstruction accuracy of 7 key HCP task-contrast maps and resting-state FC using 200 connectome eigenmodes of varying connectome densities.** See Section S4.2 and Table S2 for details about the contrast maps. wm = working memory. The solid line corresponds to the reconstruction accuracy achieved by 200 geometric eigenmodes. The dotted lines correspond to connectome densities of 0.1% and 1.55% used to generate the connectome and density-matched connectome eigenmodes in Fig. 2, respectively.

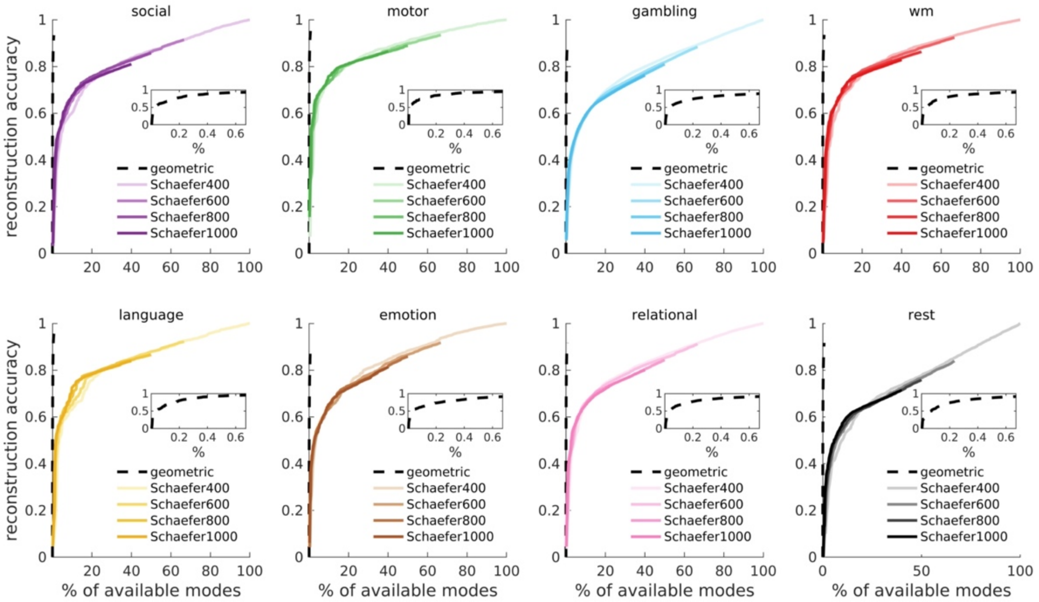


**Fig. S11. Reconstruction accuracy of 7 key HCP task-contrast maps and resting-state FC achieved by geometric eigenmodes and discrete connectome eigenmodes for different parcellation resolutions.** See Section S4.2 and Table S2 for details about the contrast maps. wm = working memory. For all cases, we use a maximum of 200 modes to directly compare the results to the rest of the study. The dashed lines represent results achieved by geometric eigenmodes (Fig. 2A); see magnified version in the insets. The lines with light to dark colors represent results achieved by discrete connectome eigenmodes using a connectivity matrix parcellated at increasing resolutions. The reconstruction accuracies are plotted versus the percentage of modes used with respect to the dimension of each full basis set (i.e., total number of available), which corresponds to the number of vertices of the cortical surface (for geometric eigenmodes) or number of parcels in each hemisphere (for discrete connectome eigenmodes). Hence, for the geometric eigenmodes, we use the first 200 modes out of 32,492 available modes per hemisphere, which is why the dotted lines terminate at 0.6% of available modes. For the discrete connectome eigenmodes using parcellations of Schaefer400, Schaefer600, Schaefer800, and Schaefer1000, we use the first 200 modes out of 200, 300, 400, and 500 available modes per hemisphere, respectively. Hence, the solid lines terminate at 100%, 66.7%, 0.50%, and 0.40% of available modes, respectively. The results emphasize that geometric eigenmodes provide the most parsimonious and compact representation of brain activity.

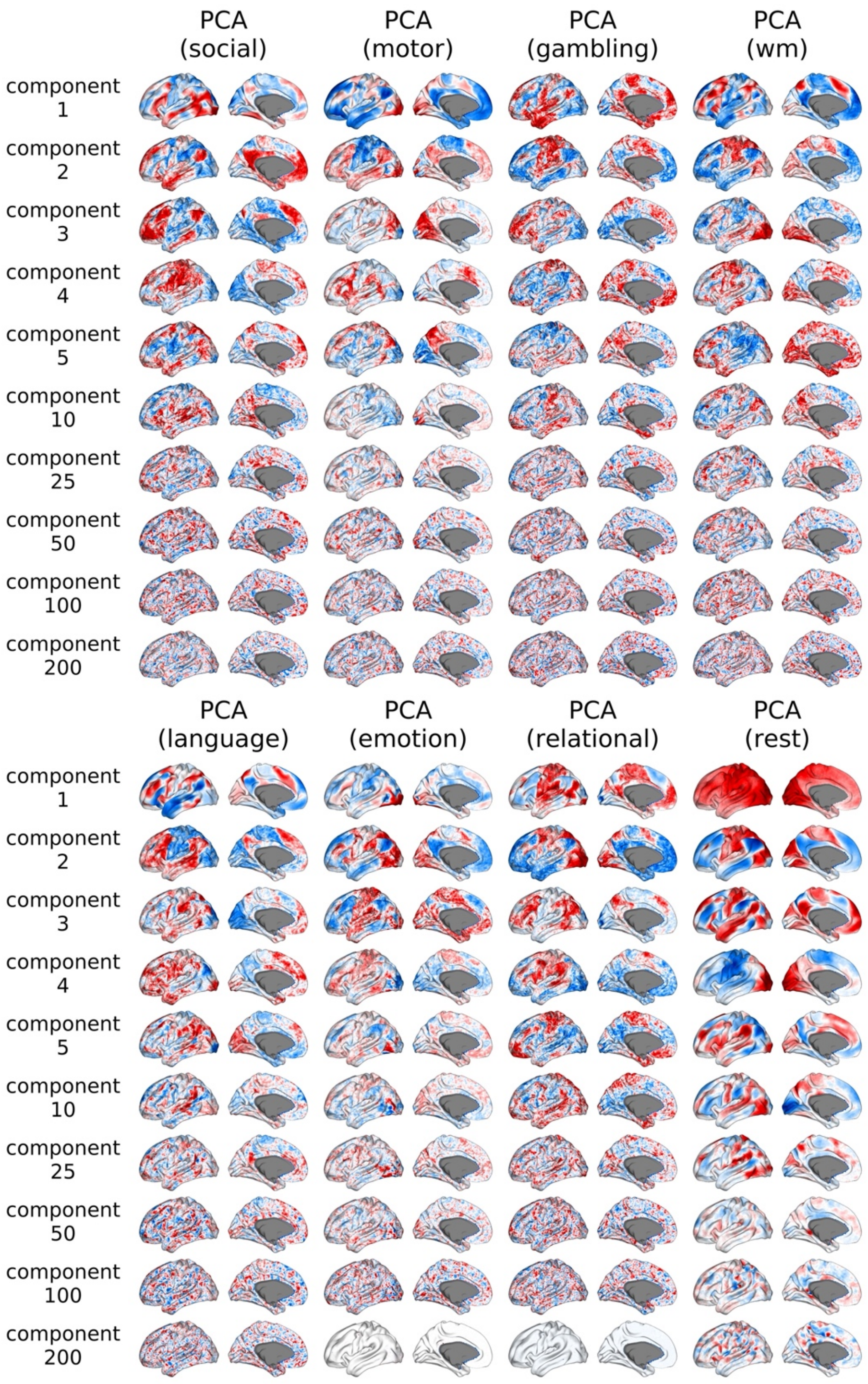


**Fig. S12. Principal components obtained via principal component analysis (PCA) of fMRI data.** The PCAs are trained on each of the 7 key HCP task-contrast maps and resting-state time series of 200 individuals. See Section S4.2 and Table S2 for details about the contrast maps. wm = working memory. Principal components 1–5, 10, 25, 50, 100, and 200 are shown from top to bottom. Negative–zero–positive values are colored as blue–white–red.

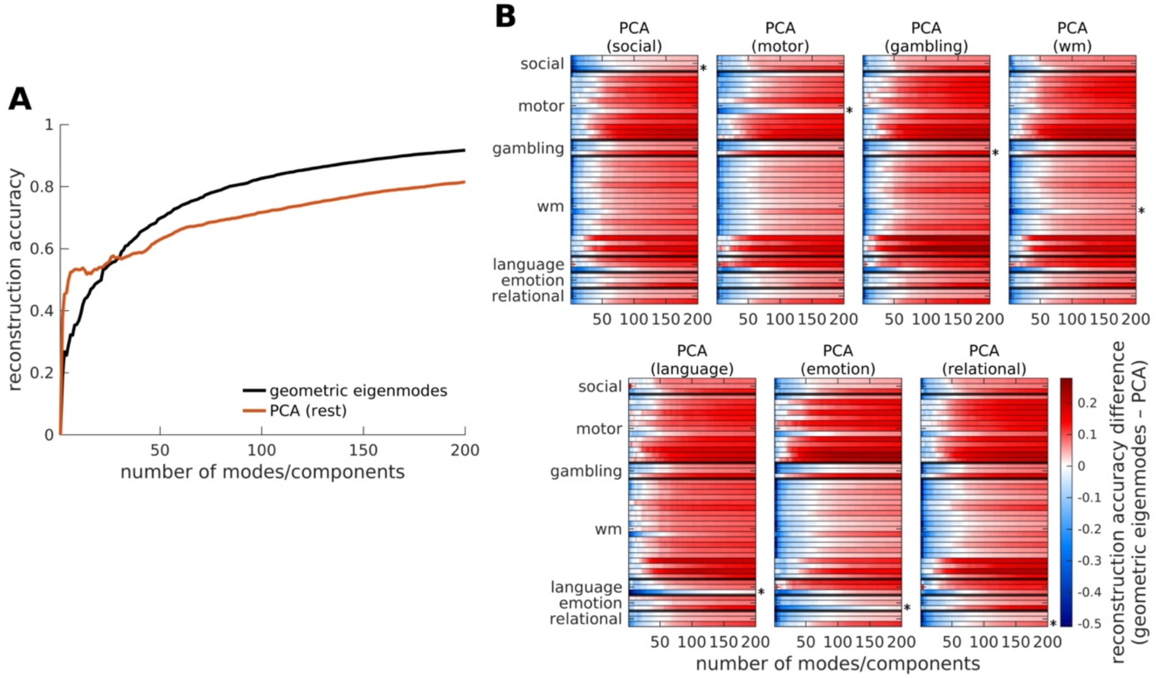


**Fig. S13. Reconstruction accuracy achieved by geometric eigenmodes and PCA.** (**A**) Reconstruction accuracy of resting-state FC. (**B**) Comparison of the reconstruction accuracy of all 47 HCP task-contrast maps, which have been grouped here by broad types (Section S4.2 and Table S2). wm = working memory. Each row represents a different task contrast. Red indicates superior performance for geometric eigenmodes. The asterisk denotes the contrast (i.e., the 7 key HCP task contrasts) within the relevant task used to train the PCA.

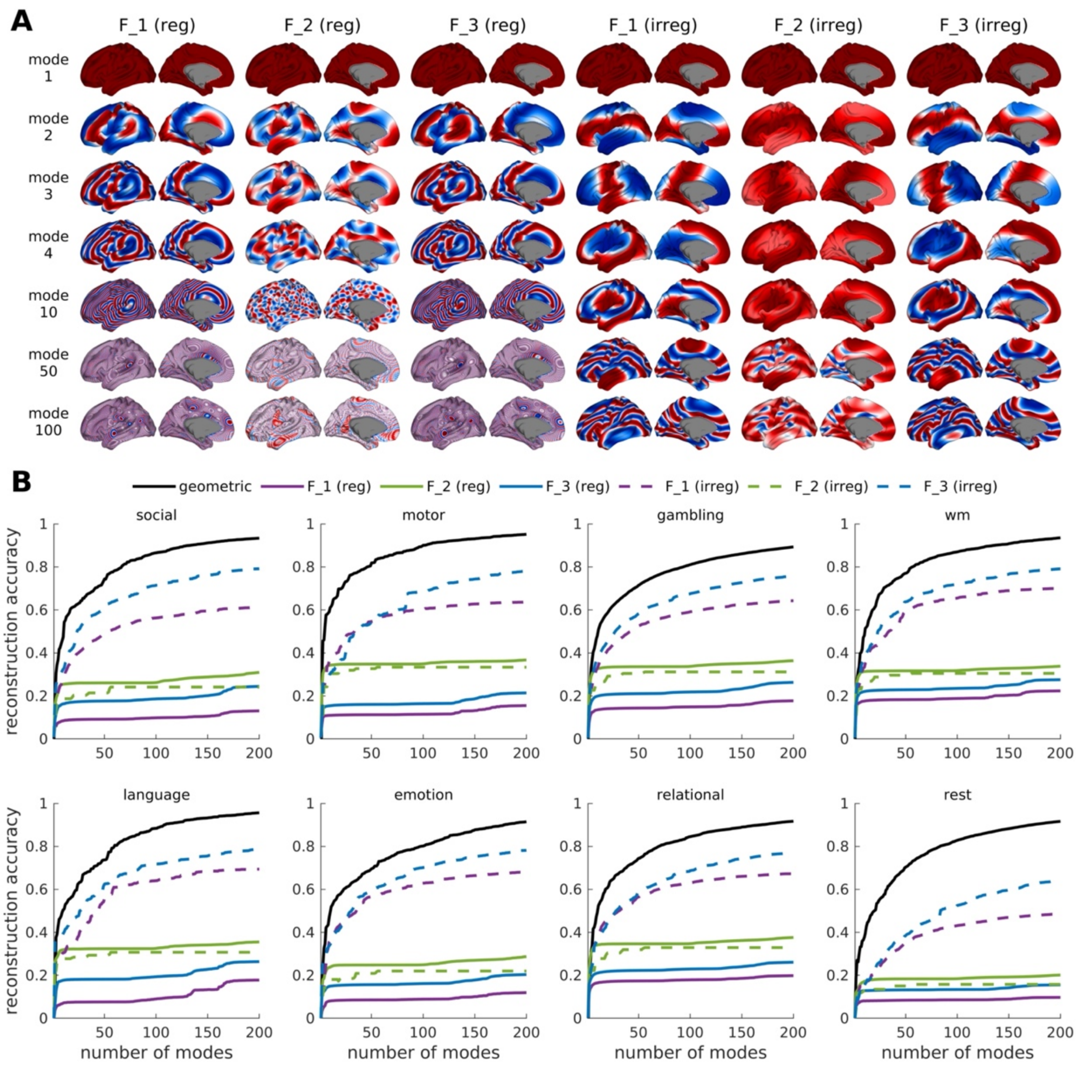


**Fig. S14. Comparison of geometric eigenmodes and Fourier basis sets.** (**A**) Spatial maps of modes 1, 2, 3, 4, 10, 50, and 100 of six different Fourier basis sets with unit coefficients. The terms reg and irreg mean that the spatial wavelengths of the modes in the x-, y-, and z-directions are spaced in regular and irregular increments, respectively. See Supplementary Material-S9 for details. (**B**) Reconstruction accuracy of 7 key HCP task-contrast maps and resting-state FC. See Section S4.2 and Table S2 for details about the contrast maps. wm = working memory.

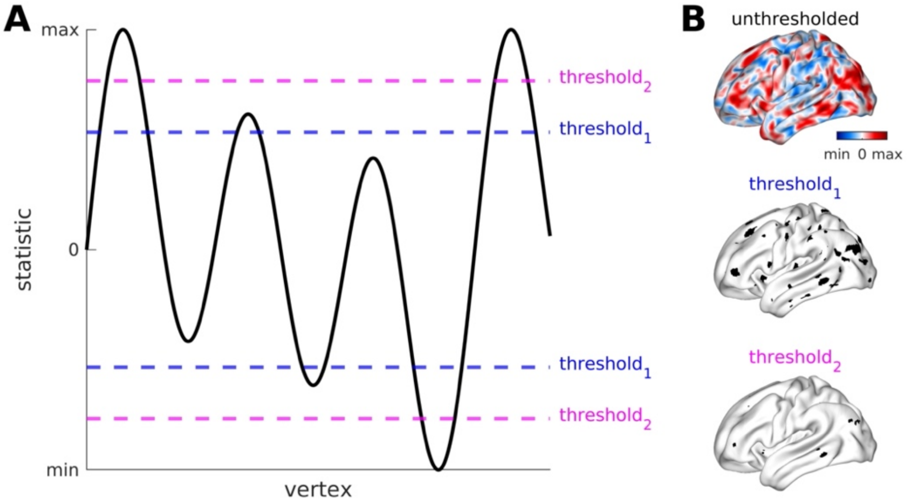


**Fig. S15. Classical neuroimaging approach of thresholding statistical maps.** (**A**) Simple one-dimensional example of how different thresholds only capture focal clusters of activations and ignore the underlying structured pattern of activations. (**B**) Spatially embedded demonstration of the concept depicted in panel A using unthresholded and binarized thresholded maps.

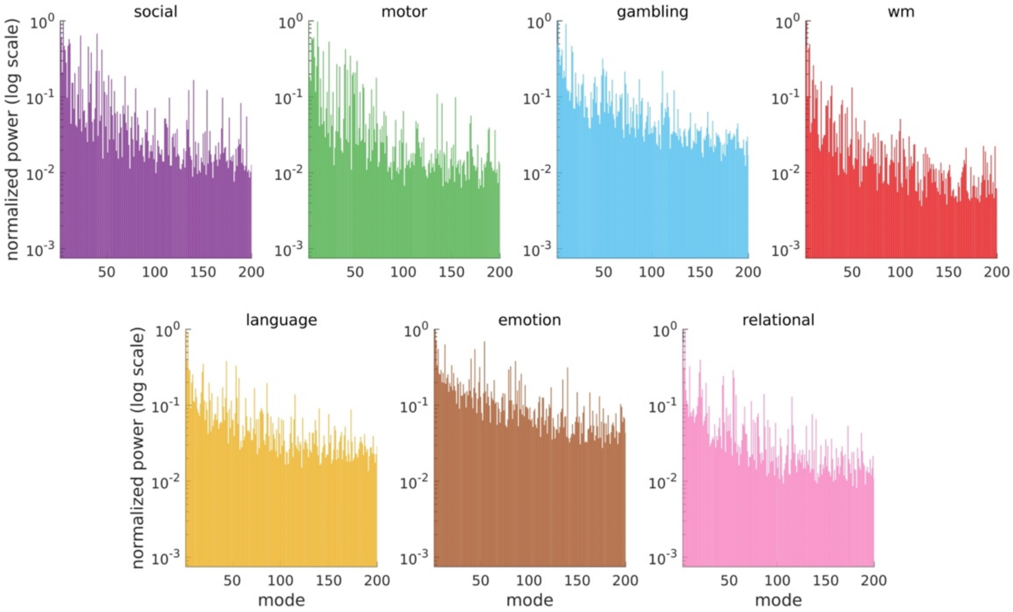


**Fig. S16. Normalized modal power spectrum of each of the 7 key HCP task-contrast maps.** See Section S4.2 and Table S2 for details about the contrast maps and Section S10 for details about calculation of the power spectrum. wm = working memory.

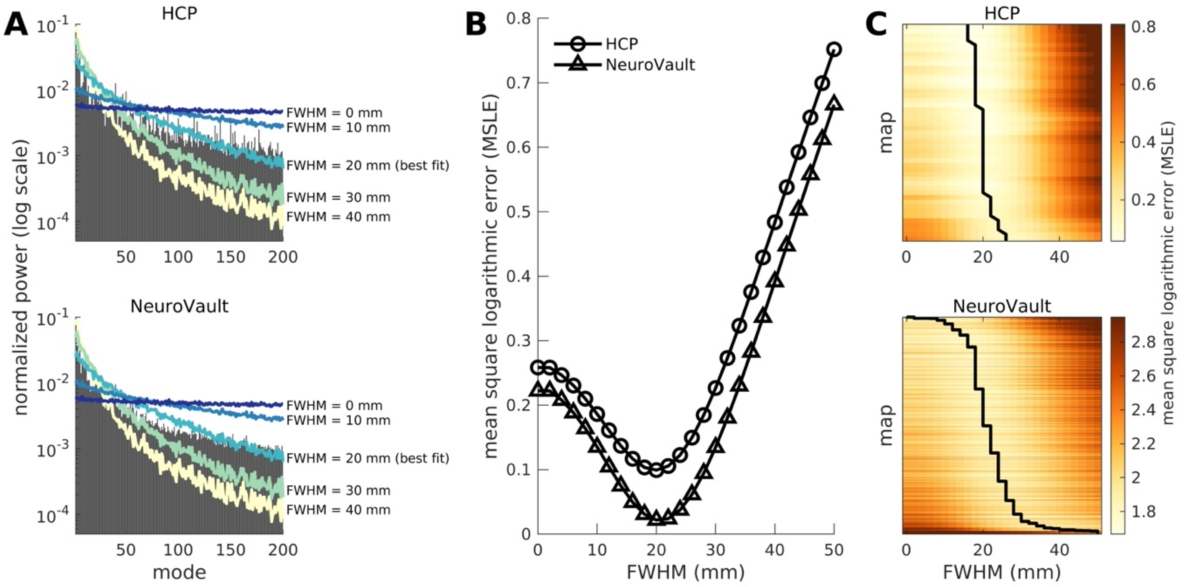


**Fig. S17. Modal power spectra of empirical task-activation maps and surrogate maps.** (**A**) Normalized mean power spectra of 47 HCP task-contrast maps (top) and 10,000 contrast maps from the NeuroVault database (bottom). The colored lines correspond to power spectra of surrogate data following the application of spatial smoothing filters with varying full-width at half-maximum (FWHM). (**B**) Average mean square logarithmic error (MSLE) as a function of FWHM between normalized mean power spectra of HCP and NeuroVault contrast maps and smoothed surrogate data. (**C**) MSLE separately obtained between the power spectra of each of the 47 HCP and 10,000 NeuroVault contrast maps and the smoothed surrogate data. Each row represents a different task-contrast map. The lines correspond to the FWHM where MSLE is minimum for each map.

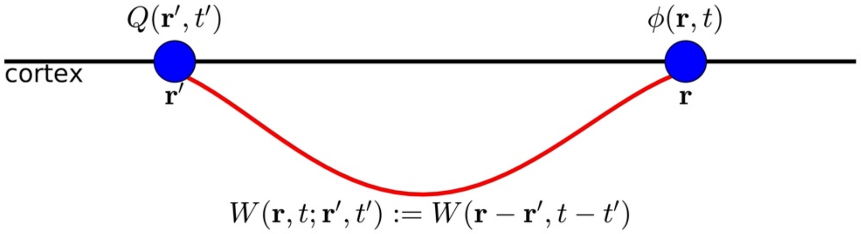


**Fig. S18. Schematic of signal propagation in neural field theory.** Two points on the cortex at locations $\boldsymbol{r}'$ and $\boldsymbol{r}$ are connected by a white-matter tract (red curve). For an isotropic medium, the activity $\phi(\boldsymbol{r}, t)$ is a convolution of the source $Q(\boldsymbol{r}', t')$ and a connectivity kernel $W(\boldsymbol{r}, t; \boldsymbol{r}', t')$ imposed by the white-matter connection, which depends only on the spatial separation, $\boldsymbol{r} - \boldsymbol{r}'$, and the time separation, $t - t'$.

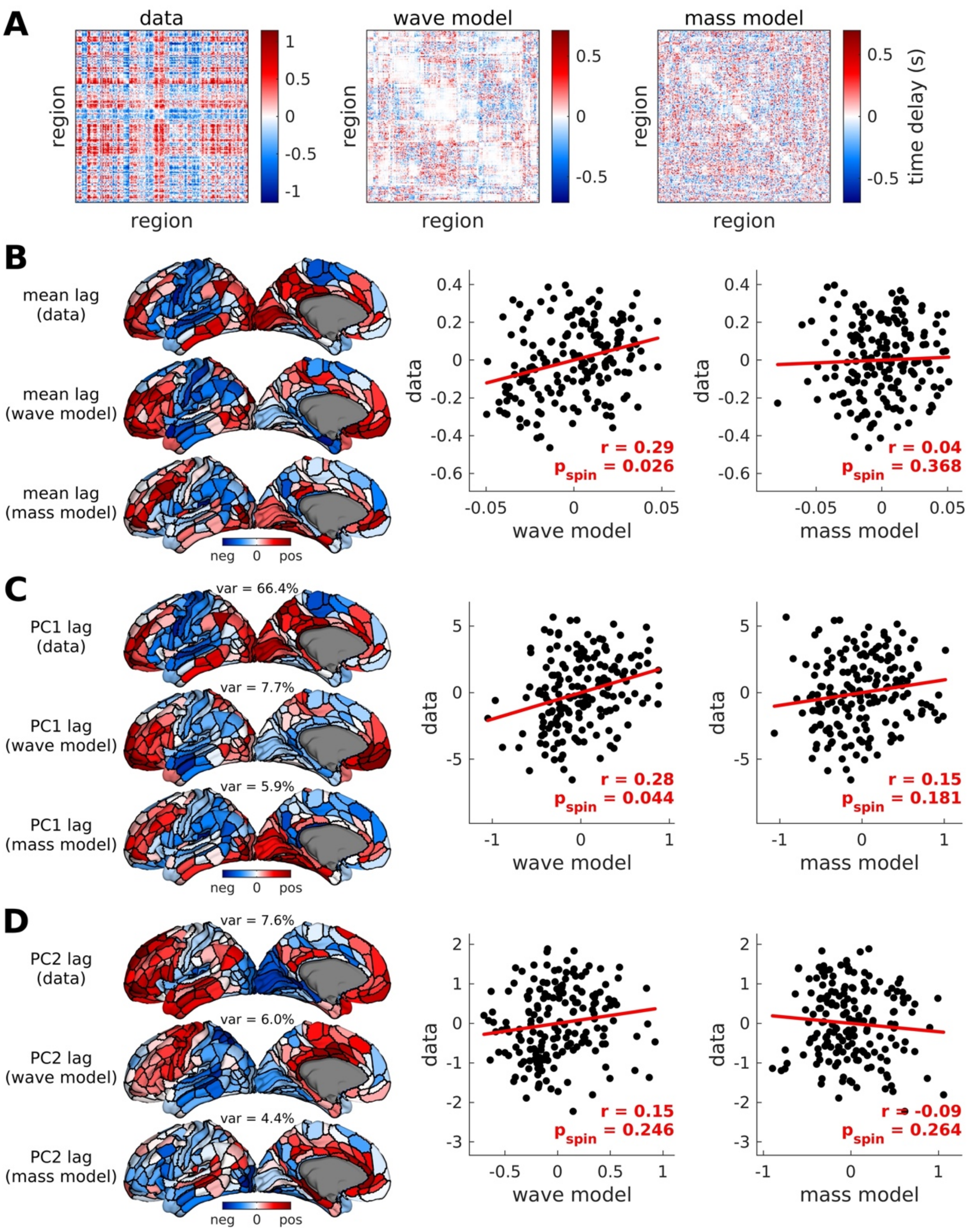


**Fig. S19. Comparison of the wave and mass models in capturing time-lagged properties of fMRI data.** (**A**) Time-delay matrices from empirical data, simulated data using the wave model, and simulated data using the mass model of the left hemisphere. Negative–zero–positive values are colored as blue–white–red. (**B**) Mean lags from the matrices in panel A (mean of each column) projected on the cortical surface. Negative–zero–positive values are colored as blue–white–red. The scatter plots show the relationship of mean lags from empirical data and simulated data from the two models. The red line represents a linear fit with Pearson correlation coefficient $r$ and spin-test $p$-value $p_{\text{spin}}$ from 10,000 permutations. (**C**) Similar to panel B but on the first principal component (PC1) of the matrices in panel A. The number above the surfaces (var) corresponds to the variance explained by the PC. (**D**) Similar to panel B but on the second PC (PC2) of the matrices in panel A.

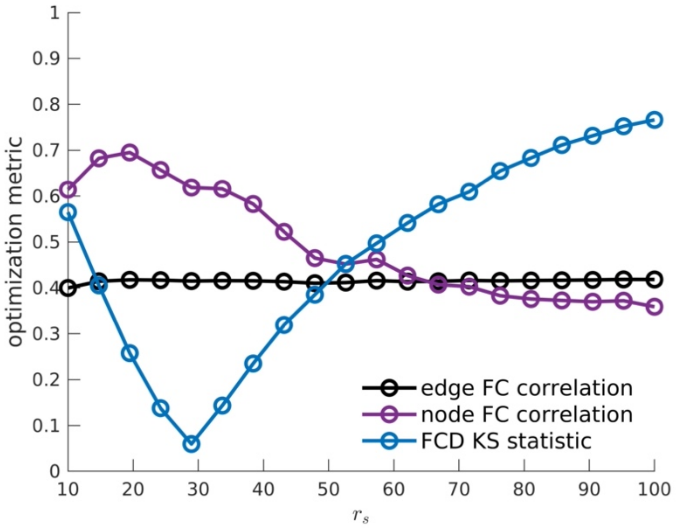


**Fig. S20. Optimization of the wave model.** The model is trained on 125 HCP individuals to find the optimal value of the parameter $r_s$ (mm). Optimization performance compares data and model FC based on the following metrics: edge FC correlation, node FC correlation, and FCD KS statistic (Section S12.4). Higher edge FC correlation, higher node FC correlation, and lower FCD KS statistic correspond to better model fit. We take $r_s = 28.9$ mm as the optimal parameter as it leads to the minimum FCD KS statistic value.

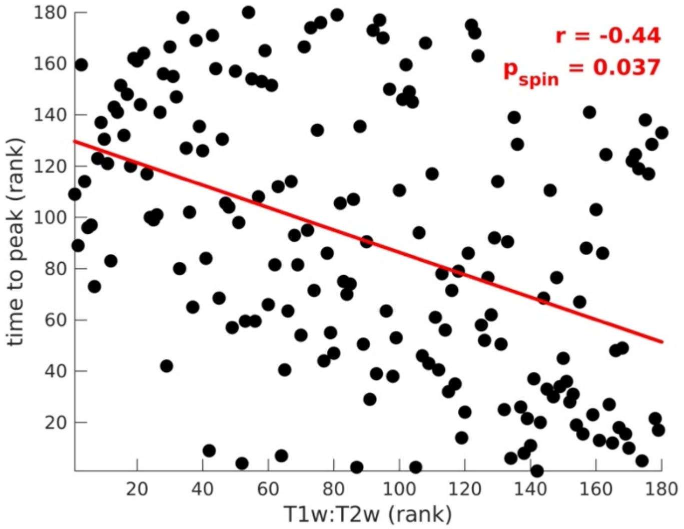


**Fig. S21. Comparison of time to peak activation and T1w:T2w of each region.** Relationship of the ranked activity profile time to peak and ranked T1w:T2w value of all brain regions. The red line represents a linear fit of the ranked variables with Spearman correlation coefficient $r$ and spin-test $p$-value $p_{\text{spin}}$ from 10,000 permutations.
